# Mammalian olfactory cortex neurons retain molecular signatures of ancestral cell types

**DOI:** 10.1101/2023.08.13.553130

**Authors:** S. Zeppilli, A. Ortega Gurrola, P. Demetci, D. H. Brann, R. Attey, N. Zilkha, T. Kimchi, S. R. Datta, R. Singh, M. A. Tosches, A. Crombach, A. Fleischmann

## Abstract

The cerebral cortex diversified extensively during vertebrate evolution. Intriguingly, the three-layered mammalian olfactory cortex resembles the cortical cytoarchitecture of non-mammals yet evolved alongside the six-layered neocortex, enabling unique comparisons for investigating cortical neuron diversification. We performed single-nucleus multiome sequencing across mouse three- to six-layered cortices and compared neuron types across mice, reptiles and salamander. We identified neurons that are olfactory cortex-specific or conserved across mouse cortical areas. However, transcriptomically similar neurons exhibited area-specific epigenetic states. Additionally, the olfactory cortex showed transcriptomic divergence between lab and wild-derived mice, suggesting enhanced circuit plasticity through adult immature neurons. Finally, olfactory cortex neurons displayed marked transcriptomic similarities to reptile and salamander neurons. Together, these data indicate that the mammalian olfactory cortex retains molecular signatures representative of ancestral cortical traits.

## Introduction

Sensory systems evolved to support behavioral adaptations in diverse ecological environments (*1–3*). For example, during the transition from aquatic to terrestrial life, animals adapted olfactory circuits to find food, detect predators, and locate mates (*4–6*). Olfactory areas are thought to have dominated the pallium (cortex) of early vertebrates (*7–9*). Over time, the pallium underwent remarkable diversification in its cellular and circuit organization to accommodate increasing demands for neural sensory processing (*2*, *7*).

A criterion historically used to infer the evolution of the pallium is its cytoarchitecture, which defines the six-layered neocortex as a novel cortical trait in mammals that emerged approximately 200 million years ago (MYA) (*10*). In contrast, two/three-layered structures are considered ancestral cortical traits, which date back at least to the last common ancestor of tetrapods, 350 MYA (*4*). Models of brain evolution propose that neocortex evolved from the expansion of an ancestral dorsal pallium, and that cell biological innovations arose from new combinations and modifications of an ancestral genetic toolkit (*7*, *11*, *12*).

Intriguingly, the three-layered cytoarchitecture of the mammalian olfactory cortex resembles the pallium of non-mammalian vertebrates, including reptiles and amphibians. This structurally conserved trait has been referred to as paleocortex, despite differences in function and developmental origin (*13–15*). Thus, the mammalian olfactory cortex provides a unique point of comparison, as its cytoarchitecture is conserved across species yet its neurons evolved in parallel with the emergence of the neocortex (**Fig. 1A**). A key question that remains is how cell types in diverse cortical structures co-evolved. We here test the premise that olfactory cortex neurons retained molecular signatures of ancestral cortical cell identity in the mammalian brain.

**Fig. 1:**
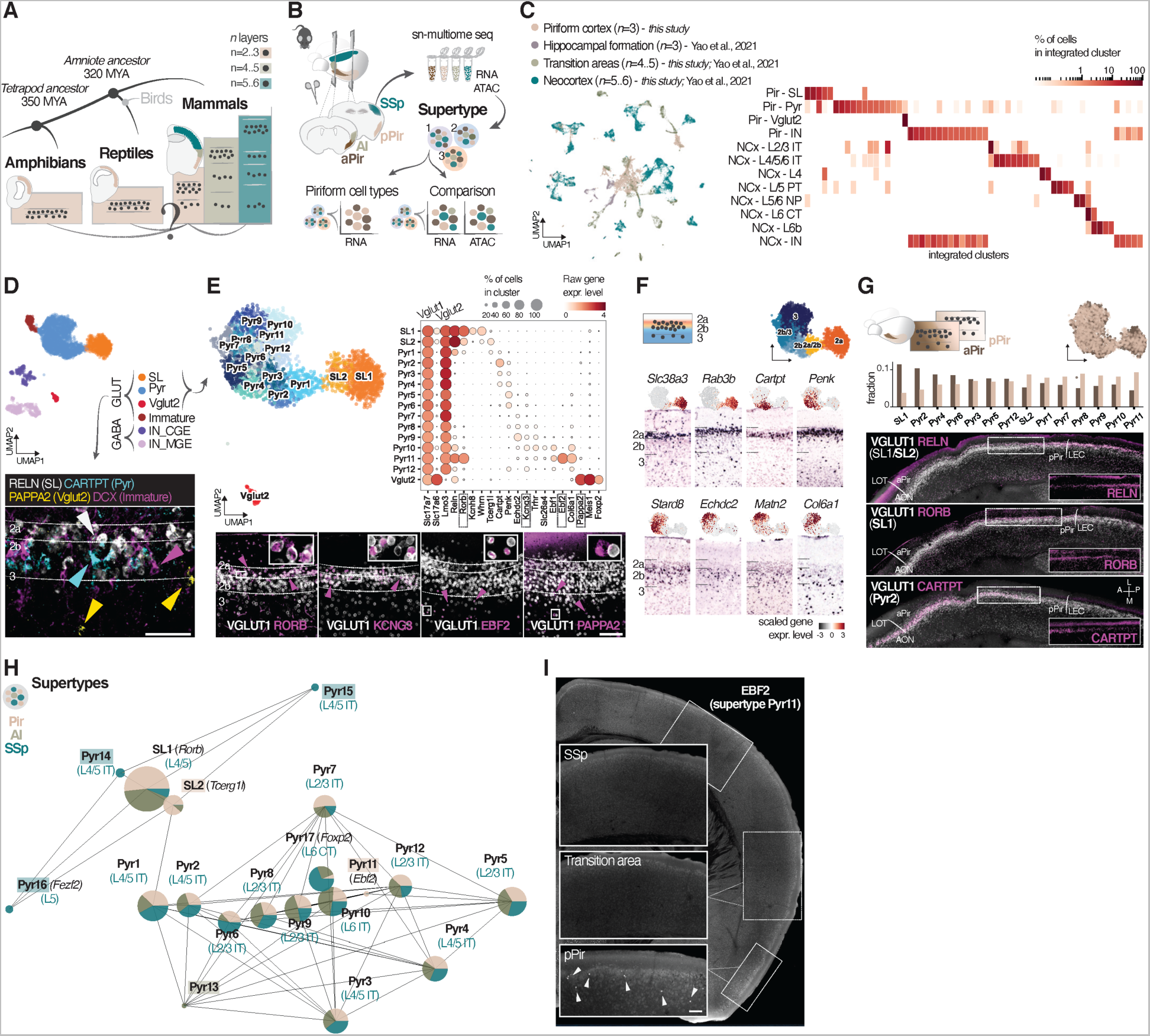
Transcriptomic diversity of piriform cortex glutamatergic neurons. **(A)** A two- to three-layered cytoarchitecture is observed throughout the pallium of amphibians and reptiles, and in the olfactory cortex of mammals, while a four- to six-layered cytoarchitecture is only present in mammals. **(B)** Overview of the experimental and computational workflow: microdissection of anterior and posterior piriform cortex (aPir and pPir), agranular insular cortex (AI) and primary somatosensory cortex (SSp) of adult mice, single-nucleus multiome sequencing (sn-multiome seq), and unsupervised clustering of all neurons into supertypes. **(C)** Integration of neurons from *this study* and from a mouse single-cell reference atlas (*17*) using Canonical Correlation Analysis (CCA). Left: UMAP representation of all integrated neurons (n=30,553). Neurons from the reference atlas are grouped by cortical regions with similar cytoarchitectures (*n*=number of layers). Right: heatmap showing quantification of co-clustering between piriform and neocortical neurons. Color of the rectangles indicates the percentage of neurons (rows) co-clustering in the seurat integrated clusters (columns). SL: semilunar cell; Pyr: Pyramidal cell; IN: inhibitory cell; L: layer; NCx: neocortex; IT: intratelencephalic; PT: pyramidal tract; NP: near projecting; CT: cortico-thalamic. **(D)** Top: UMAP representation of combined aPir and pPir neurons grouped by glutamatergic (GLUT) and GABAergic (GABA) neuron types: semilunar cells (SL); pyramidal cells (Pyr); inhibitory neurons from caudal ganglionic eminence (IN_CGE); inhibitory neurons from medial ganglionic eminence (IN_MGE). Bottom: immunohistochemistry shows expression of representative markers for piriform glutamatergic types. Scale bar, 100 μm. **(E)** Top left: UMAP representation of combined aPir and pPir neurons grouped by glutamatergic neuron subtypes (immature neurons are excluded). Top right: dot plot showing gene expression levels of laminar- and subtype-specific markers for piriform glutamatergic neuron subtypes. Bottom: immunohistochemistry using *Vglut1*-CRE/INTACT-GFP transgenic mice shows expression of representative markers for piriform glutamatergic neuron subtypes (magenta arrowheads). Higher magnification inserts show co-expression of the markers with VGLUT1. Scale bar, 100 μm. **(F)** Top: scheme of cell type gradient across cortical depth from SL (in orange) to Pyr (in blue) cells, together with UMAP representation of piriform glutamatergic neuron subtypes grouped by layers. Bottom: UMAP representations showing gene expression of markers distributed across cortical depth from layers 2a to 3, and respective *in situ* hybridization (ISH) images from the Allen Brain Atlas. Marginal zone up **(G)** Top: scheme of cell type gradient across anterior-posterior axis, together with UMAP representation color-coded by aPir and pPir datasets. Middle: bar charts showing the fraction of aPir and pPir neurons for each glutamatergic subtype. Bottom: immunohistochemistry in horizontal slices of *Vglut1*-CRE/INTACT-GFP transgenic mice shows expression of anterior- or posterior-specific markers (magenta) along the anterior-posterior axis of the piriform cortex. Inserts show the marker without VGLUT1 staining. LOT: lateral olfactory tract; AON: anterior olfactory nucleus; LEC: lateral entorhinal cortex. **(H)** Partition-based graph abstraction (PAGA) plot represents a topology-preserving map of supertypes from piriform (aPir and pPir are combined), AI, and SSp. Each supertype (node) is represented as a pie chart showing the relative contribution of the cortical areas, and edges between the nodes represent their neighborhood transcriptomic relationships. Supertype labels are based on piriform nomenclature, accompanied by their respective SSp labels based on (*17*) (**fig. S6B**). Transcription factors (TFs) specific for a particular supertype (*Ebf2* in Pyr11, *Rorb* in SL1, *Tcerg1l* in SL2), or established markers for projection neuron types (*Fezf2* in Pyr16, *Foxp2* in Pyr17) are also shown for reference. **(I)** Immunohistochemistry for the TF EBF2 corroborates the supertype Pyr11 as a small glutamatergic neuron subtype unique to Pir layer 3.

## Results

### Neuronal diversity across the mouse three- to six-layered cortical continuum

We performed single-nucleus multiome (RNA and ATAC) sequencing of four micro-dissected cortical areas of adult mice, namely the anterior and posterior three-layered piriform cortex (aPir and pPir), the four-layered agranular insular cortex (AI), and the six-layered primary somatosensory cortex (SSp) **(Fig. 1B and fig. S1)**. After stringent quality control for both RNA and ATAC data, we integrated 7,840 high-quality nuclei into a single dataset, representing various neuronal and non-neuronal cell types based on the expression of well-established markers **(Methods, figs. S2 and S3)**. We then subclustered neurons for all subsequent analyses, representing a core dataset of 5,483 high-quality nuclei across the four cortical areas.

As an initial reference, we used gene expression and canonical correlation analysis (*16*) to integrate piriform neurons with neurons from a single-cell reference atlas that included mouse neocortical areas (NCx), transition areas, such as AI and lateral entorhinal cortex (LEC), and the hippocampal formation (*17*) **(Fig. 1C and fig. S5, A to D)**. Piriform neurons were grouped into four main neuron types: *Vglut1*-expressing semilunar (SL) cells, pyramidal (Pyr) cells, *Vglut2*-expressing neurons (Vglut2), and inhibitory neurons (INs). Piriform INs co-clustered with INs from all cortical areas (**Fig. 1C and fig. S5, A and B**). In contrast, piriform glutamatergic neurons co-clustered with 83% of neurons from transition and hippocampal formation areas, and only with 8% of neurons from neocortex **(Fig. 1C, fig. S5D and Methods)**. Specifically, piriform semilunar cells co-clustered with neocortex layer (L) 4, AI, and LEC L2a neurons, while piriform pyramidal cells co-clustered with neocortex, retrohippocampal areas, AI, and LEC L2/3 intratelencephalic neurons **(Fig. 1C and fig. S5D)**. This large-scale comparison is consistent with previous studies indicating region-specific sets of glutamatergic neurons and region-invariant sets of INs across cortical areas and species (*17–19*).

Hereafter, we focus on neurons from our single-nucleus multiome sequencing dataset. Unsupervised clustering of the gene expression data identified 27 transcriptomically-defined clusters, which we refer to as supertypes (**Fig. 1, B and H**). Supertypes provide a consistent reference for comparing gene expression and chromatin accessibility states across mouse cortices. Piriform neurons were grouped into types and subtypes. Neuron types included 2 types of inhibitory GABAergic neurons, namely caudal and medial ganglionic eminence-derived INs (CGE and MGE respectively), 3 types of mature glutamatergic neurons (SL, Pyr, Vglut2), and 1 type of immature glutamatergic neurons (**Fig. 1D**). INs from CGE and MGE groups were further divided into 2 and 3 subtypes, respectively (**fig. S4G)**. Mature glutamatergic neurons were further divided into 2 subtypes of semilunar cells, 12 subtypes of pyramidal cells, and 1 subtype of Vglut2 neurons (**Fig. 1E**). We identified differentially expressed genes specific to types and subtypes, whose expression patterns were validated using immunohistochemistry and RNA *in situ* hybridization data from the Allen Brain Atlas (**Fig. 1, D and E, fig. S4**).

We also identified spatially orthogonal axes of variation of piriform *Vglut1*-expressing neurons (SL and Pyr cells) across cortical depth and the anterior-posterior axis (**Fig. 1, F and G**). Vglut1 neuron subtypes exhibited a layer-specific distribution or transition profiles between layers, accompanied by markers spanning this laminar gradient (**Fig. 1F**). L2a was composed of subtypes SL 1-2; L2b of Pyr 2-3-4; L3 of Pyr 8-9-10-11-12. Pyr1 subtype was located between L2a and L2b, and subtypes Pyr 5-6-7 between L2b and L3 (**Fig. 1F**). Along the anterior-posterior axis of piriform cortex, subtypes SL1 and Pyr2 were enriched in anterior, while subtypes SL2 and Pyr 8-9-11 in posterior piriform (**Fig. 1G**). Immunohistochemistry confirmed these anterior-posterior gradients: SL1-specific RORB and Pyr2-specific CARTPT expression were higher in anterior piriform, while RELN expression, enriched in SL2 cells, was higher in posterior piriform (**Fig. 1, E and G**).

We next quantified the co-clustering of glutamatergic neurons from the piriform cortex (Pir), the transition area AI, and the neocortical area SSp within each supertype, and inferred their connectivity profiles by integrating SSp datasets from this study and from the single-cell reference atlas (*17*) **(Fig. 1H and fig. 6B)**. Supertype SL2, selectively expressing the transcriptional regulator *Tcerg1l*, was composed of only piriform semilunar cells. In contrast, 7% of SSp L4 intratelencephalic projection neurons co-clustered with (anterior) piriform semilunar cells within the SL1 supertype, characterized by high expression levels of the thalamorecipient-specific transcription factor (TF) *Rorb,* potentially reflecting a shared molecular signature of sensory input neurons in Pir (aPir) and SSp **(Fig. 1H and fig. S6D).** 90% of pyramidal neurons co-clustered across cortical areas as intratelencephalic projection neurons **(Fig. 1H and fig. S6B).** Specifically, piriform neurons of L2a/2b co-clustered with SSp neurons of L4/5, piriform neurons of L2b/3 with SSp neurons of L2/3, and piriform neurons of L3 with SSp neurons of L6 (**Fig. 1H**). In contrast, the supertype Pyr11, selectively expressing the TF *Ebf2*, was only composed of piriform cells **(Fig. 1, E, H and I, figs. S4B and S5E)**. SSp-specific supertypes corresponded to L4/5 intratelencephalic projection neurons (Pyr 14-15), L5 extra-telencephalic projection neurons (Pyr16, *Fezf2^+^*), and L6 corticothalamic projection neurons (Pyr17, *Foxp2^+^*) (**Fig. 1H and fig. S6D**). Finally, we identified a set of TFs that were highly enriched in piriform compared to other cortical areas, including *Ebf1, Ebf2, Tfap2d, Dach1, Dach2, Glis3* and *Rarb* (**Fig. 1I and fig. S5E**). In summary, our data provide a comprehensive molecular characterization of piriform neurons. We identify piriform-specific glutamatergic neurons, as well as neurons whose molecular identities were similar across mouse cortical areas. These identities corresponded primarily to intratelencephalic projection neurons. In contrast, neocortical extra-telencephalic projection neurons did not co-cluster with any piriform subtype.

### Transcriptomically similar glutamatergic neurons differ in chromatin accessibility

The transcriptomic co-clustering of piriform and SSp neurons suggests that their chromatin accessibility states may be similarly shared. However, unsupervised clustering of the ATAC sequencing data revealed an unexpected misalignment between transcriptome- and epigenome-based supertypes. The greatest divergence was observed between Pir and SSp glutamatergic neurons, with AI neurons exhibiting an intermediate epigenetic profile. In contrast, transcriptomically-defined INs aligned well with ATAC-based clusters **(Fig. 2A and fig S7)**. To explore further such epigenomic divergence, we applied SCENIC+, a prediction framework that uses RNA and ATAC data to infer enhancer-driven gene regulatory networks (e-GRNs) (*20*). An e-GRN is composed of e-regulons, which in turn consist of TFs, their target enhancers, and their downstream target genes **(Fig. 2B)**. We computed e-GRNs for aPir, pPir, AI, and SSp neurons, resulting in 161, 150, 135, and 124 high-quality e-regulons, respectively **(fig. S8, A and B)**. We then focused on the 35 e-regulons found in each of the four cortical areas **(fig. S8, C and D)**. These e-regulons exhibited specificity either for a particular supertype, such as *Rorb* (SL1), for a particular layer, such as *Glis3* (SSp L2/3, Pir L2b/3) and *Rfx3* (SSp L2/3, Pir L2b/3), or were active in all neurons, such as *Mef2c, Tcf4,* and *Zpf148* **(fig. S9)**. Remarkably, in each of the 35 e-regulons, TFs had highly area-specific target enhancers and target genes **(Fig. 2C and fig. S10)**. For example, the TF *Rorb*, specific to the supertype SL1, shared only 7% of its target enhancers and 9% of its target genes between aPir and SSp neurons **(Fig. 2D and fig. S11, A and B)**. Nonetheless, target genes identified for one cortical area were still expressed in the other areas, contributing to the global transcriptomic similarity **(fig. S11, C to F)**. However, target genes in the other areas were predicted to be regulated by overlapping yet distinct TF combinations **(fig. S12B)**. Examples of area-specific TF combinations and enhancers for the potassium channel *Kcnk2*, a *Rorb*-specific target gene shared across cortical areas, are shown in **(Fig. 2E and fig. S12A)**.

**Fig. 2:**
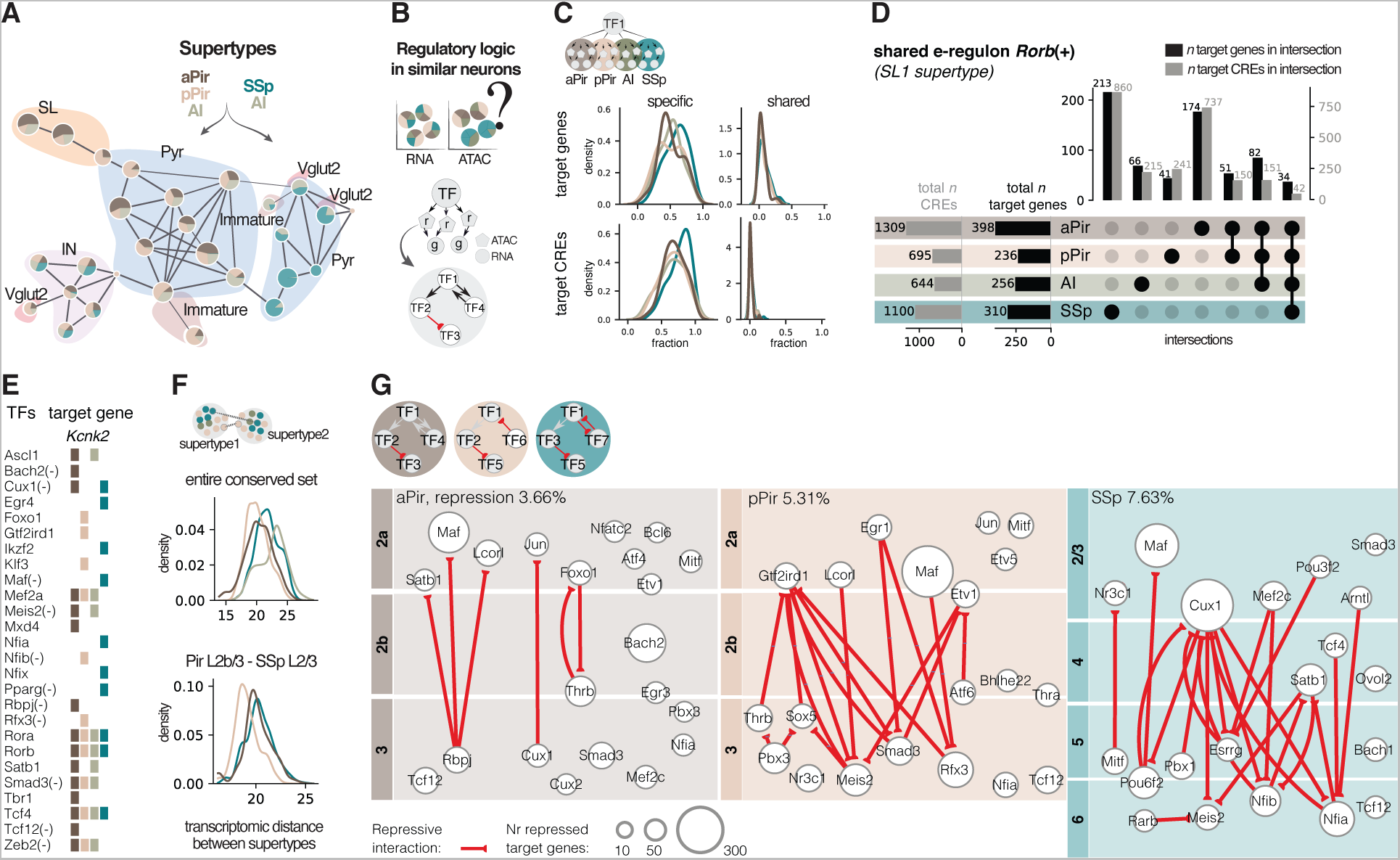
Area-specific epigenetic states distinguish transcriptomically similar glutamatergic neurons across the adult mouse three- to six-layered cortical continuum. **(A)** PAGA plot representing RNA-defined supertypes projected onto ATAC-defined clusters (nodes). Background area color-coded by main neuron type: SL, Pyr, IN, immature and Vglut2 neurons. Each ATAC cluster is represented as a pie chart that shows the relative contribution of the four cortical areas. **(B)** Top: scheme illustrating the computational strategy to investigate differential regulatory mechanisms underlying transcriptomically similar cell types across mouse areas. Bottom: scheme of an e-regulon composed of the TF, enhancer regions the TF binds to (r), and the target genes the TF regulates (g). e-Regulons are then combined into an enhancer-driven gene regulatory network (e-GRN). **(C)** Kernel-density estimate plots showing the fraction of target genes (top) and target *cis*-regulatory elements (CREs) (bottom) of the 35 shared e-regulons across areas. Left: fractions specific to a single area; right: fractions shared across areas. **(D)** Upset plot of the intersection of target genes (black bars) and target CREs (gray bars) for the e-regulon *Rorb*(+) across aPir, pPir, AI and SSp. The vertical bars show the number of target genes and CREs in the corresponding intersection of the matrix below. The horizontal bars show the total number of target genes and CREs for each cortical area. **(E)** Matrix showing an example of TF combinations for each cortical area. The potassium channel *Kcnk2* is a target gene of the TF *Rorb* in all four cortical areas. **(F)** Quantification of cell type discreteness per cortical area by considering transcriptomically similar neurons across all layers (corresponding to supertypes SL1, Pyr 1-2-3-4-5-6-7-8-9-10-12) or within a transcriptomically similar layer (e.g., Pir 2b/3 and SSp 2/3, corresponding to supertypes Pyr 5-6-7-8-9-12). Areas are downsampled to the same number of neurons and re-processed. **(G)** Quantification (%) of predicted repressive interactions between TFs (nodes) for each cortical area. Interactions are visualized as networks laid out along cortical depth, with TFs placed in the center of their expression laminar domain. Node size indicates the number of repressor-specific target genes.

Given that transcriptomic and epigenomic profiles of INs aligned well across cortical areas, the differences observed for glutamatergic neurons are unlikely to be solely driven by the sparse nature of ATAC data. Therefore, the transcriptomic similarity observed between glutamatergic neurons across mouse cortical areas likely arises from area-specific epigenetic states and the differential use of TF combinations. Conserved transcriptomic yet divergent epigenetic states in neuron types have similarly been observed across species (*20*, *21*).

### Higher overlap in gene expression and limited transcriptional repression in piriform compared to primary somatosensory cortex

A prominent feature observed in the clustering of piriform glutamatergic neurons were the graded changes in transcriptomic profiles **(Fig. 1, E to G).** We therefore compared cluster discreteness across cortical areas by measuring, for each area, the minimum distance between supertypes containing glutamatergic neurons from the four areas. This analysis was conducted across all layers, or selectively on transcriptomically similar layers (e.g., Pir L2b/3 and SSp L2/3). We found that clusters exhibited significant greater transcriptomic overlap among each other in Pir compared to SSp **(Fig. 2F)**. A high overlap in gene expression may result from co-expression of key TFs for cell type identity previously observed in piriform but not in neocortex (*22*), and from limited repression between TFs across cell types and cortical layers (*23*, *24*). We thus quantified repressive interactions in each area-specific e-GRN by considering only supertypes containing glutamatergic neurons from aPir, pPir, and SSp **(fig. S13A)**. Repression here is defined as anti-correlated expression between a TF and its target genes, while enhancers linked to the target genes are accessible (*20*). We found that the degree of transcriptional repression was lower in Pir compared to SSp (aPir: 3.66%; pPir: 5.31%; SSp: 7.63%). The majority of repressive interactions distinguished Pir and SSp neurons along cortical depth **(Fig. 2G)**. For example, in aPir and pPir, repressive interactions occurred primarily between L2a and L3. Notably, aPir and pPir neurons exhibited pronounced differences in these interactions, despite repressors being expressed in both areas **(Fig. 2G and fig. S13B)**.

Together, these findings suggest that transcriptional repression may contribute to the observed differences in discreteness between Pir and SSp clusters across cortical depth, and further highlight distinct regulatory mechanisms along the anterior-posterior axis of the piriform cortex.

### Posterior piriform cortex exhibits pronounced transcriptomic divergence between lab and wild-derived mice

Our transcriptomic analysis identified a population of immature neurons, enriched in pPir **(Fig. 3A)**. Immunohistochemistry for DCX, a widely used marker for immature neurons (*25*), revealed an anterior-posterior gradient of immature cells, with the highest levels in pPir **(Fig. 3B)**. This population exhibited a glutamatergic phenotype (predominantly *Vglut2*) and expressed canonical markers of adult immature neurons such as *Sox11* and *Sox4* (*26*), as well as specific markers such as *St8sia2*, *Igfbpl1* and *Zfp57* **(Fig. 3C and fig. S4F)**. We also observed the presence of SSp neurons in the immature neuron supertype. However, given their transcriptomic similarity to L4/5/6 intratelencephalic Car3^+^ neurons and the absence of DCX expression in SSp we do not consider these neurons as immature (**fig. S6, B and C**). Our analysis aligns with previous studies reporting the presence of embryonically-generated adult immature neurons in piriform cortex (*27*). These neurons have been suggested to enhance circuit plasticity in brain areas lacking adult neurogenesis (*28*).

**Fig. 3:**
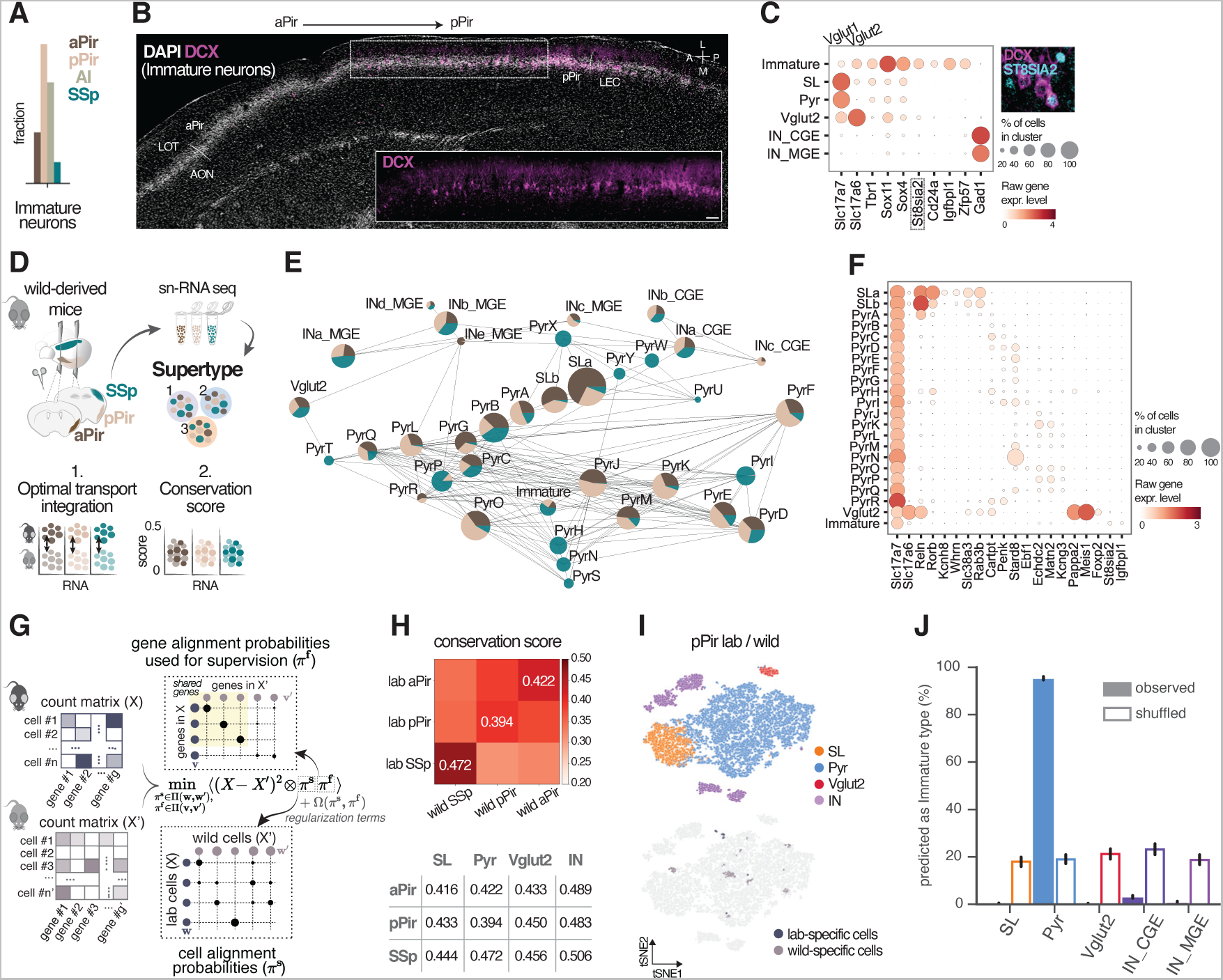
A potential link between immature neurons and the transcriptomic divergence of pyramidal cells between lab and wild-derived mice in posterior piriform cortex. **(A)** Bar chart showing the fraction of the immature neuron supertype across aPir, pPir, AI, and SSp. **(B)** Immunohistochemistry in horizontal slices shows expression of DCX (in magenta), a marker for immature neurons along the anterior-posterior axis of the piriform cortex. Insert shows DCX expression without DAPI. LOT: lateral olfactory tract; AON: anterior olfactory nucleus; LEC: lateral entorhinal cortex. **(C)** Dot plot showing gene expression levels of generic or specific markers for piriform immature neurons. Top right: immunohistochemistry in pPir shows co-expression of DCX with the immature neuron-specific marker ST8SIA2. **(D)** Overview of the experimental and computational workflow: microdissection of aPir, pPir and SSp of adult wild-derived mice, single-nucleus RNA sequencing (sn-RNA seq), and unsupervised clustering of all neurons into supertypes. For each cortical area, neurons of lab and wild-derived mice are integrated using optimal transport (OT) and their transcriptomic similarity is quantified through a conservation score. **(E)** PAGA plot representing supertypes from aPir, pPir and SSp, with pie chart showing the relative contribution of the three cortical areas. **(F)** Dot plot showing gene expression levels of laminar- and subtype-specific markers across piriform glutamatergic neuron subtypes of the wild dataset used similarly to distinguish subtypes in the lab dataset. **(G)** Scheme of the OT pipeline used to integrate sc-RNA seq count matrices of lab mice (X, top) and wild-derived mice (X’, bottom). The OT formulation finds the two alignment maps, π^s (mapping cells) and π^f (mapping genes), by minimizing the sum of squared error between the two count matrices when cells and genes of one dataset are transformed into the other by multiplying with these maps. See **Methods** for details. **(H)** Top: matrix plot of average conservation scores for aPir, pPir and SSp. The scores represent the median probability of transcriptomic similarity between lab and wild datasets quantified by considering all neurons. Bottom: table of average conservation scores for each area quantified by considering only neurons of a particular cell type: SL, Pyr, Vglut2 and IN. A conservation score of 0.5 indicates a perfect mixing between lab and wild datasets. **(I)** tSNE representations of integrated pPir neurons of lab and wild-derived mice. Top: tSNE representation color-coded by main neuron types. Bottom: tSNE representation color-coded by misaligned neurons between lab and wild datasets. **(J)** Linear Support Vector Classifier (SVC) showing the percentage of each neuron type predicted as immature neurons in combined lab and wild pPir datasets. Classification and generalization performance were at chance levels when classifiers were trained on data with permuted neuron type labels (shuffled).

To establish a potential link between piriform immature neurons and increased plasticity-driven cellular divergence, we conducted a comparison of Pir and SSp cell types between two mouse strains derived from different environments, inbred lab mice and outbred wild-derived mice (*29*, *30*). We performed single-nucleus multiome and RNA sequencing of micro-dissected aPir, pPir, and SSp of adult wild-derived mice **(Fig. 3D and fig. S14)**. After stringent quality control, we integrated 26,975 high-quality nuclei into a single dataset, referred to here as the wild dataset, comprising neuronal and non-neuronal cell types with good alignment with the lab dataset **(figs. S15 and S16, Methods**). We subclustered mature and immature neurons, resulting in a core dataset of 24,901 high-quality nuclei across aPir, pPir and SSp. Unsupervised clustering of gene expression data identified 36 transcriptomically-defined supertypes **(Fig. 3E)**. The classification of piriform neuron subtypes was largely consistent between lab and wild datasets **(Fig. 3, E and F).**

We then asked to what extent cortical areas of lab and wild-derived mice differed in their cellular components. We quantified, for each cortical area, the correspondence probability for each pair of cells between lab and wild datasets using optimal transport (*31*). These probabilities were used to co-embed the lab and wild datasets into a shared low-dimensional space **(Fig. 3, D and G, Methods)**. To measure the degree of neuron-neuron similarity, we defined a conservation score, which is a modification of the local inverse Simpson’s index (*32*). A score of 0.5 indicates perfect overlap between lab and wild datasets **(Fig. 3D, Methods)**. We found that pPir exhibited the lowest conservation score, while SSp the highest (aPir: 0.422 (95% CI: 0.405-0.439), pPir: 0.394 (95% CI: 0.378-0.433), SSp: 0.472 (95% CI: 0.456-0.489)) **(Fig. 3H, top)**. In pPir, pyramidal cells had the lowest conservation score **(Fig. 3H, bottom)**, and misaligned neurons were distributed across distinct pyramidal subtypes, rather than forming a transcriptomically-defined cluster **(Fig. 3I).** To corroborate these results, we integrated the lab and wild pPir datasets using the Single-cell Variational Inference (scVI) model (*33*). In the scVI latent space, pyramidal cells showed the least similarity between datasets **(fig. S17A)**, and OT-identified misaligned pyramidal cells (wild-specific) were less likely to share neighbors with neurons from lab mice **(fig. S17B)**.

Finally, to further assess a potential relatedness between pyramidal and immature neurons, we trained linear classifiers on main neuron types (SL, Pyr, Vglut2, INs) from combined lab and wild pPir datasets, achieving 98.8% accurate type distinction **(fig. S17C)**. We then applied the classifiers to immature neurons, which predicted immature neurons as pyramidal cells with 95.4% accuracy **(Fig. 3J)**. Immunohistochemistry further supported this relatedness by revealing co-expression of DCX with CUX1, a marker for pyramidal cells, but not with RELN and GABA, markers for semilunar cells and inhibitory neurons, respectively **(fig. S17D**). These results are consistent with previous studies showing that adult immature neuron differentiate into glutamatergic neurons in piriform layer 2 (*34*). Together, our data reveal greater cell type variation in Pir compared to SSp, potentially linked to a plasticity-driven differentiation of adult immature neurons into molecularly diverse pyramidal cells.

### Piriform glutamatergic neurons display marked transcriptomic similarities to neurons of non-mammals

Piriform cortex is central to the historical definition of paleocortex, as its three-layered cytoarchitecture resembles the pallium of amphibians and reptiles (*14*, *15*, *19*). This raises the question whether neurons from conserved cytoarchitectures exhibit similar molecular profiles **(Fig. 4A)**, as speculated based on single-cell RNA sequencing studies in non-mammals (*19*, *35*). We used gene expression and canonical correlation analysis (*16*) to integrate our piriform dataset with published single-cell RNA sequencing datasets of the pallium of representative tetrapod groups, namely mammals (*17*), reptiles (*36–38*), and amphibians (*19*). Published single-cell RNA sequencing data comprised mouse neocortical areas including SSp, motor, and visual cortex, transition areas including AI and LEC, and the hippocampal formation; turtle and lizard medial (MCtx), dorsal (DCtx), and lateral (LCtx) cortex, and the anterior dorsal ventricular ridge (aDVR); salamander medial (MP), dorsal (DP), lateral (LP), and ventral (VP) pallium **(Fig. 4B and fig. S18)**. Data integration generates new clusters, enabling comparisons of neuron types across areas and species by quantifying the co-clustering of neurons within each integrated cluster **(Fig. 4A)**. Our analysis yielded 39 integrated clusters, whose composition was consistent with previously identified transcriptomic similarities and differences (Woych et al., 2022) **(figs. S18, S19, S20, S21)**. For example, INs co-clustered across cortical areas, neurons of the mouse hippocampal formation co-clustered with those of medial regions of reptiles and salamander, and neocortex extra-telencephalic neurons did not co-cluster with any other cell type **(figs. S19, S20, S21)**. Surprisingly, however, the inclusion of non-mammalian datasets revealed that piriform glutamatergic neurons exhibited greater transcriptomic similarity to glutamatergic neurons of turtle, lizard and salamander than to those of the neocortex **(Fig. 1C and fig. S22A)**. The majority of piriform glutamatergic neurons were present in integrated clusters 1 (25% of all piriform neurons), 4 (14%), 5 (17%), and 14 (22%), with clusters 1 and 5 also composed of neurons from neocortex and non-mammalian species **(fig. S22A)**. In cluster 1, piriform neurons co-clustered only with 2% of neocortex neurons, in contrast to over 20% of reptile DCtx, LCtx, and aDVR neurons, and more than 50% of salamander DP and VP neurons. In cluster 5, piriform neurons co-clustered primarily with neurons from lateral olfactory areas of reptiles (10%) and salamander (49%) **(fig. S22A)**.

**Fig. 4:**
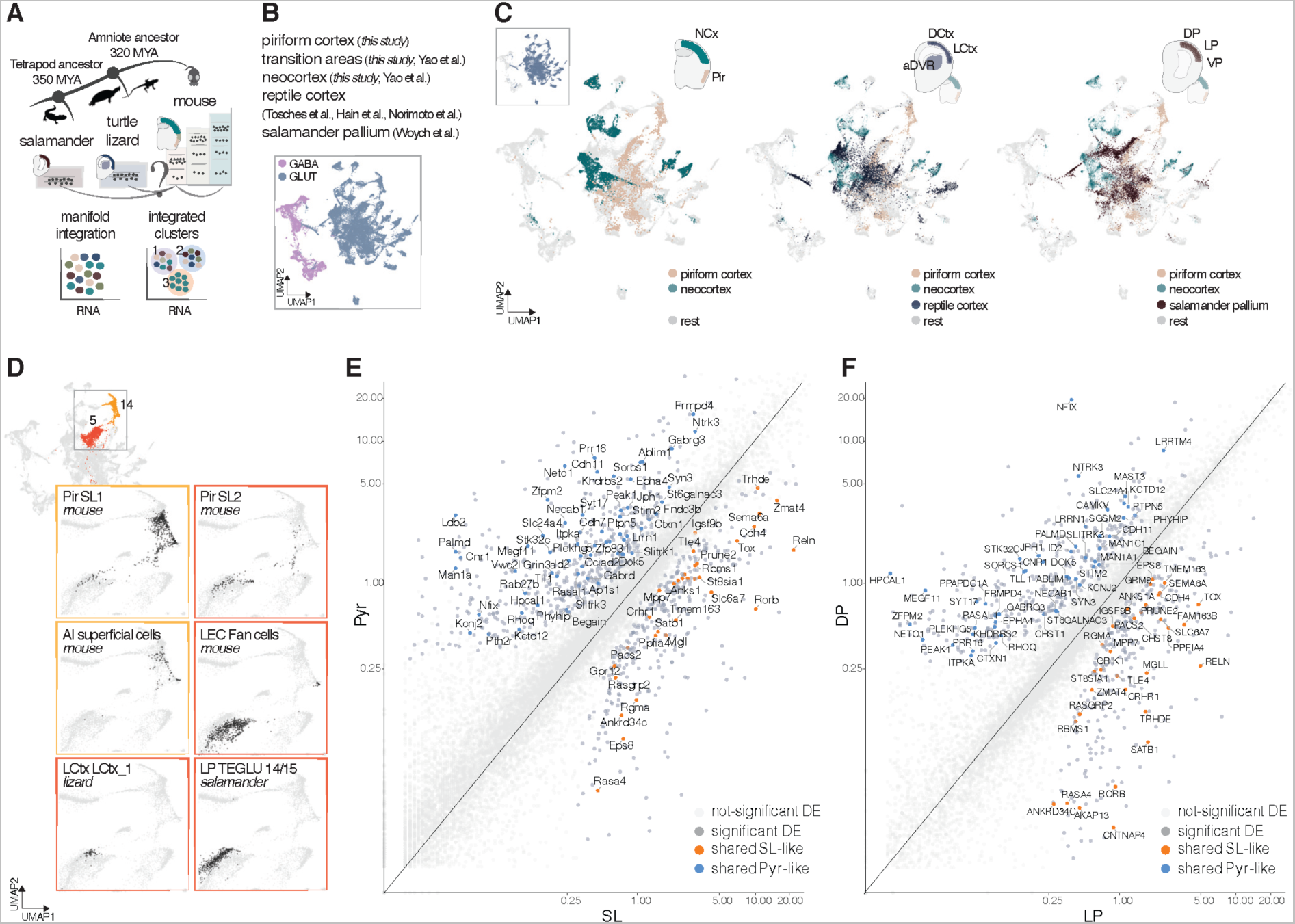
Piriform glutamatergic neurons display marked transcriptomic similarity to reptile and salamander neurons. **(A)** Sc-RNA seq datasets from salamander, turtle, lizard and mouse (*this study*, single-cell reference atlas) are integrated using CCA in seurat. The analysis generates new integrated clusters, in which is quantified the co-clustering between datasets to infer conserved or novel neuron types across tetrapods. **(B)** UMAP representation of integrated neurons color-coded by glutamatergic (GLUT) or GABAergic (GABA) phenotype. **(C)** UMAP representations of the integration and visualization of only glutamatergic neurons from the dorsolateral cortical continuum of each species. From left to right: piriform and neocortex glutamatergic neurons only; neurons of the reptile dorsolateral cortex (turtle and lizard are combined) visualized over piriform and neocortex glutamatergic neurons; neurons of salamander dorsal to ventral pallium visualized over piriform and neocortex glutamatergic neurons. NCx: neocortex; DCtx: dorsal cortex; LCtx: lateral cortex; aDVR: anterior dorsal ventricular ridge; DP, LP and VP: dorsal, lateral and ventral pallium. **(D)** Top insert: UMAP representation colored by the integrated clusters 5 and 14 comprising SL cells. Rest: UMAP representations of neurons from mouse and non-mammalian cortical areas that co-clustered with piriform SL cells in integrated clusters 5 and 14. **(E)** Scatter plot of differentially expressed genes (DEGs) between SL and Pyr cells. Each dot represents the mean gene expression value. Orange and blue dots depict shared genes enriched both in SL and LP neurons, or in Pyr and DP neurons, respectively. Other statistically significant DEGs are shown in dark gray, while non-statistically significant DEGs in light gray. **(F)** Scatter plot as shown in **(E)** but of DEGs between LP and DP glutamatergic neurons.

We next focused on the two main types of piriform glutamatergic neurons, semilunar (SL) and pyramidal (Pyr) cells. Semilunar cells co-clustered with functionally and developmentally related neurons of lateral olfactory cortical areas. Specifically, integrated cluster 14 comprised only mouse glutamatergic neurons, including 92% of piriform SL1 and 21% of SL2 cells, 96% AI cells, and 4% of LEC fan cells **(Fig. 4D and fig. S22B)**. In contrast, integrated cluster 5 comprised glutamatergic neurons from all species, including 7.5% of piriform SL1 and 76% of SL2 cells, 96% of LEC fan cells, 94% of lizard LCtx superficial cells (Reln^+^, putative bowl cells (*39*)), and 63% and 86% of two salamander LP clusters (Reln^+^, superficial cells) **(Fig. 4D and fig. S22B)**. These results suggest the presence of ancestral molecular signatures of intratelencephalic input neurons in lateral olfactory cortical areas across tetrapods, see also (*40*), and the specialization of a subset of intratelencephalic input neurons in mammals, situated in the most anterior part of the olfactory cortex.

In contrast to semilunar cells, piriform pyramidal cells co-clustered with a heterogenous mixture of pyramidal neurons from dorsal, lateral, and ventral cortical areas of different developmental origins, such as mouse neocortex and LEC L2/3 intratelencephalic neurons, reptile LCtx and aDVR neurons, and salamander DP and VP deep layer neurons **(fig. S23)**. These results suggest the presence of ancestral molecular signatures of intratelencephalic non-input neurons in dorsal, lateral and ventral cortical areas across tetrapods.

Finally, we independently computed differentially expressed genes between piriform pyramidal *versus* semilunar cells, and between salamander neurons that co-clustered with them, namely deep DP or VP *versus* LP neurons, respectively. We identified genes that were shared between piriform pyramidal and DP or VP neurons, and between piriform semilunar and LP neurons **(Fig. 4, E and F, fig. S24A)**. These genes were then used to define a SL-like or a Pyr-like gene module. The SL-like module showed specific enrichment in neurons of lateral olfactory areas expressing *Reln*, namely piriform semilunar cells, LEC fan cells, and LP neurons, as well as in neurons of the retrosplenial cortex L4 **(fig. S24B)**. This module included TFs such as *Rorb, Satb1* and *Tox*. In contrast, consistent with the mixed clustering of pyramidal neurons, the Pyr-like module showed enrichment in neurons of medial, dorsal, lateral and ventral cortical areas, representing a generic signature of pyramidal neurons across mouse areas and species **(fig. S24B)**.

Together, our analysis indicates distinct evolutionary origins of piriform pyramidal and semilunar cells in the last common ancestor of tetrapods.

## Discussion

We here provide a comprehensive molecular characterization of cell types in the adult mouse olfactory (piriform) cortex. We characterize distinct glutamatergic subtypes of piriform semilunar, pyramidal, Vglut2-expressing and immature cells, organized along continuous transcriptomic gradients across cortical depth and the anterior-posterior axis **(Fig. 1 and fig. S4)**. We speculate that the transcriptomic differences between semilunar, pyramidal, Vglut2 and immature cells reflect molecular signatures of distinct developmental origin, connectivity and function (*22*, *41–44*).

We then compare the transcriptomes and epigenomes of glutamatergic neurons across three- to six-layered mouse cortices, as well as the transcriptomes of glutamatergic neurons across mouse, reptile and salamander pallia (cortices). Altogether, these comparisons reveal that olfactory cortex glutamatergic neurons retain distinctive molecular signatures of ancestral cortical cells in the mammalian brain. These signatures include area-specific epigenetic states, enhanced circuit plasticity, and pronounced transcriptomic similarity with neurons of non-mammals.

Comparative enhancer-driven gene regulatory network (e-GRNs) analysis across three- to six-layered mouse cortices reveals highly area-specific epigenetic states and the differential use of transcription factor (TF) combinations within transcriptomically similar glutamatergic neurons **(Fig. 2, C and E, figs. S7, S10, S11, S12)**. Differences in regulatory logic in the absence of major changes in global gene expression may be interpreted as an example of evolvability through weak regulatory linkage, a concept proposed to play a key role in the development and evolution of morphological traits (*11*, *45*, *46*). The stark difference in epigenetic states we observe may represent a solution to diversify glutamatergic neurons across the cortical continuum with a limited set of TFs. Furthermore, comparative e-GRN analysis provides evidence for an expansion of transcriptional repression in the neocortex **(Fig. 2G)**. Cross-repressive interactions between TFs represent a mechanism to establish boundaries between cell types and cortical layers (*23*, *24*). The lower degree of transcriptional repression we found in piriform compared to somatosensory cortex may thus explain the observed pronounced overlap in gene expression between piriform neurons **(Fig. 2F)**. A high overlap in gene expression may result from the co-expression of key TFs for cell type identity previously observed in piriform and in the pallium of non-mammals, but not in neocortex (*22*, *47*, *48*). Such overlap may give rise to less distinct and functionally specialized neurons, potentially representing a shared ancestral signature of cortices with fewer layers. Together, our study points to evolutionary strategies in global gene regulatory mechanisms that shape glutamatergic neuronal identity and diversity.

A distinctive feature of the adult piriform cortex is the presence of embryonically-generated immature neurons, which have been proposed to contribute to circuit plasticity, facilitating adaptive changes to novel environments (*27*, *28*, *34*). We here characterize their transcriptomic profiles and explore a potential relationship between piriform adult immature neurons and plasticity-driven cellular divergence. Upon comparing the transcriptomes of anterior and posterior piriform and somatosensory cortex neurons between adult inbred (lab) and outbred wild-derived mice, we find the greatest cellular divergence in posterior piriform cortex, while the lowest in somatosensory cortex **(Fig. 3 and fig. S17)**. These results suggest enhanced adaptation of olfactory circuits, potentially driven by plasticity-driven maturation of adult immature neurons into pyramidal cells. One intriguing possibility is that these embryonically-generated piriform immature neurons represent remnants of active neural progenitors found in lateral olfactory areas of non-mammals (*49*).

Finally, within- and cross-species transcriptomic comparisons reveal the presence of molecular signatures of ancestral glutamatergic (projection) neurons in the mammalian olfactory cortex, namely signatures of intratelencephalic input and non-input neurons. Some of the transcriptomic similarities between intratelencephalic input neurons may reflect homologous relationships between sister cell types, as their alignment is also consistent with a common developmental origin. These intratelencephalic input neurons, including piriform semilunar cells, receive sensory input from the olfactory bulb and are located in superficial layers of lateral olfactory cortical areas in mice, reptiles and salamander **(Fig. 4D and fig. S22B),** see also (*40*). In contrast, transcriptomic similarities between intratelencephalic non-input neurons are not consistent with a common developmental origin in extant tetrapods, suggesting these similarities may have arisen either through convergent evolution or through duplication and divergence of homologous intratelencephalic non-input neurons in earlier vertebrates. These intratelencephalic non-input neurons, including piriform pyramidal cells, are located in deeper layers of ventral, lateral, and dorsal cortical areas in mice, reptiles and salamander **(fig. S23)**.

More generally, our cross-species transcriptomic analysis reveals unexpected similarities between piriform glutamatergic neurons and glutamatergic neurons of reptiles and salamander. For example, salamander dorsal pallium neurons exhibited greater co-clustering with piriform neurons than with neocortex neurons **(Fig. 4C, fig. S22A).** The *superimpositio lateralis* model posits that an ancestral dorsal pallium co-opted developmental programs from dorsolateral areas, leading to cellular and molecular rearrangements that may have occurred in response to novel terrestrial pressures (*50*). Interestingly, olfactory inputs also target dorsal regions in lower vertebrates, but shifted to more lateral areas in reptiles and mammals (*51*). The transcriptomic similarity observed among a mixture of intratelencephalic projection neurons from dorsal, lateral, and ventral pallial areas in extant tetrapods may be attributed to the retention of ancestral GRNs in these areas. These ancestral GRNs may have been exapted for accommodating olfactory-driven behavioral adaptations.

In conclusion, our study provides a roadmap to explore the evolutionary logic underlying the diversification of cortical cell types and circuits in the mammalian brain.

## Acknowledgments

We thank Zach Herbert and Maura Berkeley from the Molecular Biology Core Facilities at the Dana-Farber Cancer Institute for sequencing services. We thank Nicole Eckart from 10xGenomics for excellent technical support. We thank Kelsey Babcock, Federica Mosti, Debra Silver, Jason Ritt, Stuart Firestein, Hugues Berry, Kevin Franks, Sophie Pantalacci, Guillaume Beslon, Hynek Wichterle, and Telmo Pievani for critical comments on the manuscript. We thank Andrea Pierre for software support and Carmen Bravo González-Blas for analysis support with SCENIC+. We thank Noa Nisim for technical assistance with the wild-derived mice and the Brown and Weizmann animal facilities for animal care.

## Author contributions

Conceptualization: SZ, MAT, AC, AF

Methodology: SZ, AOG, PD, DHB, RA, NZ, RS, MAT, AC

Investigation: SZ, AOG, PD, DHB, RS, MAT, AC, AF

Visualization: SZ, AOG, PD, DHB, RA, RS, MAT, AC

Funding acquisition: SZ, TK, RS, MAT, AC, AF

Project administration: AF

Supervision: SZ, TK, SRD, RS, MAT, AC, AF

Writing - original draft: SZ, AC, AF

Writing - review & editing: SZ, AOG, PD, DHB, NZ, TK, SRD, RS, MAT, AC, AF

## Data and materials availability

Raw and processed single nucleus multiome and RNA sequencing data have been deposited in Gene Expression Omnibus (GEO) under the accession number GSE239477 and available upon request. All other data are included in the main paper or the supplementary materials. The R and Python analysis scripts used for this paper are available at the GitLab link https://gitlab.com/fleischmann-lab/papers/zeppilli-et-al-2023.

## Supplementary Materials

Materials and Methods

Supplementary text: abbreviations

Figs. S1 to S24

References (52 to 61)

## References and Notes

1. D. M. Bear, J.-M. Lassance, H. E. Hoekstra, S. R. Datta, The Evolving Neural and Genetic Architecture of Vertebrate Olfaction. Current Biology. 26, R1039–R1049 (2016).

2. P. Cisek, Evolution of behavioural control from chordates to primates. Phil. Trans. R. Soc. B. 377, 20200522 (2022).

3. R. J. V. Roberts, S. Pop, L. L. Prieto-Godino, Evolution of central neural circuits: state of the art and perspectives. Nat Rev Neurosci. 23, 725–743 (2022).

4. M. A. MacIver, B. L. Finlay, The neuroecology of the water-to-land transition and the evolution of the vertebrate brain. Phil. Trans. R. Soc. B. 377, 20200523 (2022).

5. Y. Niimura, Evolutionary dynamics of olfactory receptor genes in chordates: interaction between environments and genomic contents. Human Genomics. 4, 107 (2009).

6. L. Weiss, I. Manzini, T. Hassenklöver, Olfaction across the water–air interface in anuran amphibians. Cell Tissue Res. 383, 301–325 (2021).

7. J. H. Kaas, The evolution of brains from early mammals to humans. WIREs Cognitive Science. 4, 33–45 (2013).

8. T. B. Rowe, T. E. Macrini, Z.-X. Luo, Fossil Evidence on Origin of the Mammalian Brain. Science. 332, 955–957 (2011).

9. T. B. Rowe, G. M. Shepherd, Role of ortho-retronasal olfaction in mammalian cortical evolution. Journal of Comparative Neurology. 524, 471–495 (2016).

10. J. H. Kaas, Evolution of the neocortex. Curr Biol. 16, R910–914 (2006).

11. S. B. Carroll, Endless Forms: The Evolution of Gene Regulation and Morphological Diversity. Cell. 101, 577–580 (2000).

12. G. F. Striedter, The Telencephalon of Tetrapods in Evolution; pp. 179–194. Brain Behavior and Evolution. 49, 179–194 (2008).

13. M. Á. García-Cabezas, B. Zikopoulos, H. Barbas, The Structural Model: a theory linking connections, plasticity, pathology, development and evolution of the cerebral cortex. Brain Struct Funct. 224, 985–1008 (2019).

14. C. A. Kappers, The Phylogenesis of the Palaeo-cortex and Archi-cortes Compared with the Evolution of the Visual Neo-cortex (J. Truscott and Son, 1909).

15. G. Laurent, J. Fournier, M. Hemberger, C. Müller, R. Naumann, J. M. Ondracek, L. Pammer, S. Reiter, M. Shein-Idelson, M. A. Tosches, T. Yamawaki, “Cortical Evolution: Introduction to the Reptilian Cortex” in Micro-, Meso- and Macro-Dynamics of the Brain, G. Buzsáki, Y. Christen, Eds. (Springer, Cham (CH), 2016; http://www.ncbi.nlm.nih.gov/books/NBK435755/).

16. Y. Hao, S. Hao, E. Andersen-Nissen, W. M. Mauck, S. Zheng, A. Butler, M. J. Lee, A. J. Wilk, C. Darby, M. Zager, P. Hoffman, M. Stoeckius, E. Papalexi, E. P. Mimitou, J. Jain, A. Srivastava, T. Stuart, L. M. Fleming, B. Yeung, A. J. Rogers, J. M. McElrath, C. A. Blish, R. Gottardo, P. Smibert, R. Satija, Integrated analysis of multimodal single-cell data. Cell. 184, 3573–3587.e29 (2021).

17. Z. Yao, C. T. J. van Velthoven, T. N. Nguyen, J. Goldy, A. E. Sedeno-Cortes, F. Baftizadeh, D. Bertagnolli, T. Casper, M. Chiang, K. Crichton, S.-L. Ding, O. Fong, E. Garren, A. Glandon, N. W. Gouwens, J. Gray, L. T. Graybuck, M. J. Hawrylycz, D. Hirschstein, M. Kroll, K. Lathia, C. Lee, B. Levi, D. McMillen, S. Mok, T. Pham, Q. Ren, C. Rimorin, N. Shapovalova, J. Sulc, S. M. Sunkin, M. Tieu, A. Torkelson, H. Tung, K. Ward, N. Dee, K. A. Smith, B. Tasic, H. Zeng, A taxonomy of transcriptomic cell types across the isocortex and hippocampal formation. Cell. 184, 3222–3241.e26 (2021).

18. J. M. Kebschull, E. B. Richman, N. Ringach, D. Friedmann, E. Albarran, S. S. Kolluru, R. C. Jones, W. E. Allen, Y. Wang, S. W. Cho, H. Zhou, J. B. Ding, H. Y. Chang, K. Deisseroth, S. R. Quake, L. Luo, Cerebellar nuclei evolved by repeatedly duplicating a conserved cell-type set. Science. 370, eabd5059 (2020).

19. J. Woych, A. Ortega Gurrola, A. Deryckere, E. C. B. Jaeger, E. Gumnit, G. Merello, J. Gu, A. Joven Araus, N. D. Leigh, M. Yun, A. Simon, M. A. Tosches, Cell-type profiling in salamanders identifies innovations in vertebrate forebrain evolution. Science. 377, eabp9186 (2022).

20. C. Bravo González-Blas, S. De Winter, G. Hulselmans, N. Hecker, I. Matetovici, V. Christiaens, S. Poovathingal, J. Wouters, S. Aibar, S. Aerts, SCENIC+: single-cell multiomic inference of enhancers and gene regulatory networks. Nat Methods, 1–13 (2023).

21. T. E. Bakken, N. L. Jorstad, Q. Hu, B. B. Lake, W. Tian, B. E. Kalmbach, M. Crow, R. D. Hodge, F. M. Krienen, S. A. Sorensen, J. Eggermont, Z. Yao, B. D. Aevermann, A. I. Aldridge, A. Bartlett, D. Bertagnolli, T. Casper, R. G. Castanon, K. Crichton, T. L. Daigle, R. Dalley, N. Dee, N. Dembrow, D. Diep, S.-L. Ding, W. Dong, R. Fang, S. Fischer, M. Goldman, J. Goldy, L. T. Graybuck, B. R. Herb, X. Hou, J. Kancherla, M. Kroll, K. Lathia, B. van Lew, Y. E. Li, C. S. Liu, H. Liu, J. D. Lucero, A. Mahurkar, D. McMillen, J. A. Miller, M. Moussa, J. R. Nery, P. R. Nicovich, S.-Y. Niu, J. Orvis, J. K. Osteen, S. Owen, C. R. Palmer, T. Pham, N. Plongthongkum, O. Poirion, N. M. Reed, C. Rimorin, A. Rivkin, W. J. Romanow, A. E. Sedeño-Cortés, K. Siletti, S. Somasundaram, J. Sulc, M. Tieu, A. Torkelson, H. Tung, X. Wang, F. Xie, A. M. Yanny, R. Zhang, S. A. Ament, M. M. Behrens, H. C. Bravo, J. Chun, A. Dobin, J. Gillis, R. Hertzano, P. R. Hof, T. Höllt, G. D. Horwitz, C. D. Keene, P. V. Kharchenko, A. L. Ko, B. P. Lelieveldt, C. Luo, E. A. Mukamel, A. Pinto-Duarte, S. Preissl, A. Regev, B. Ren, R. H. Scheuermann, K. Smith, W. J. Spain, O. R. White, C. Koch, M. Hawrylycz, B. Tasic, E. Z. Macosko, S. A. McCarroll, J. T. Ting, H. Zeng, K. Zhang, G. Feng, J. R. Ecker, S. Linnarsson, E. S. Lein, Comparative cellular analysis of motor cortex in human, marmoset and mouse. Nature. 598, 111–119 (2021).

22. A. Diodato, M. Ruinart de Brimont, Y. S. Yim, N. Derian, S. Perrin, J. Pouch, D. Klatzmann, S. Garel, G. B. Choi, A. Fleischmann, Molecular signatures of neural connectivity in the olfactory cortex. Nat Commun. 7 (2016), doi:10.1038/ncomms12238.

23. M. J. Delás, J. Briscoe, Repressive interactions in gene regulatory networks: When you have no other choice. Curr Top Dev Biol. 139, 239–266 (2020).

24. L. C. Greig, M. B. Woodworth, M. J. Galazo, H. Padmanabhan, J. D. Macklis, Molecular logic of neocortical projection neuron specification, development and diversity. Nat Rev Neurosci. 14, 755–769 (2013).

25. R. Knoth, I. Singec, M. Ditter, G. Pantazis, P. Capetian, R. P. Meyer, V. Horvat, B. Volk, G. Kempermann, Murine Features of Neurogenesis in the Human Hippocampus across the Lifespan from 0 to 100 Years. PLoS One. 5, e8809 (2010).

26. L. Mu, L. Berti, G. Masserdotti, M. Covic, T. M. Michaelidis, K. Doberauer, K. Merz, F. Rehfeld, A. Haslinger, M. Wegner, E. Sock, V. Lefebvre, S. Couillard-Despres, L. Aigner, B. Berninger, D. C. Lie, SoxC Transcription Factors Are Required for Neuronal Differentiation in Adult Hippocampal Neurogenesis. J Neurosci. 32, 3067–3080 (2012).

27. M. Á. Gómez-Climent, E. Castillo-Gómez, E. Varea, R. Guirado, J. M. Blasco-Ibáñez, C. Crespo, F. J. Martínez-Guijarro, J. Nácher, A Population of Prenatally Generated Cells in the Rat Paleocortex Maintains an Immature Neuronal Phenotype into Adulthood. Cereb Cortex. 18, 2229–2240 (2008).

28. C. La Rosa, F. Cavallo, A. Pecora, M. Chincarini, U. Ala, C. G. Faulkes, J. Nacher, B. Cozzi, C. C. Sherwood, I. Amrein, L. Bonfanti, Phylogenetic variation in cortical layer II immature neuron reservoir of mammals. eLife. 9, e55456 (2020).

29. R. A. Miller, R. Dysko, C. Chrisp, R. Seguin, L. Linsalata, G. Buehner, J. M. Harper, S. Austad, Mouse (Mus musculus) stocks derived from tropical islands: new models for genetic analysis of life-history traits. Journal of Zoology. 250, 95–104 (2000).

30. N. Zilkha, S. G. Chuartzman, Y. Sofer, Y. Pen, M. Cum, A. Mayo, U. Alon, T. Kimchi, Sex-dependent control of pheromones on social organization within groups of wild house mice. Current Biology. 33, 1407–1420.e4 (2023).

31. Q. H. Tran, H. Janati, N. Courty, R. Flamary, I. Redko, P. Demetci, R. Singh, Unbalanced CO-optimal Transport. Proceedings of the AAAI Conference on Artificial Intelligence. 37, 10006–10016 (2023).

32. I. Korsunsky, N. Millard, J. Fan, K. Slowikowski, F. Zhang, K. Wei, Y. Baglaenko, M. Brenner, P. Loh, S. Raychaudhuri, Fast, sensitive and accurate integration of single-cell data with Harmony. Nat Methods. 16, 1289–1296 (2019).

33. R. Lopez, J. Regier, M. B. Cole, M. I. Jordan, N. Yosef, Deep Generative Modeling for Single-cell Transcriptomics. Nat Methods. 15, 1053–1058 (2018).

34. P. Rotheneichner, M. Belles, B. Benedetti, R. König, D. Dannehl, C. Kreutzer, P. Zaunmair, M. Engelhardt, L. Aigner, J. Nacher, S. Couillard-Despres, Cellular Plasticity in the Adult Murine Piriform Cortex: Continuous Maturation of Dormant Precursors Into Excitatory Neurons. Cerebral Cortex. 28, 2610–2621 (2018).

35. B. M. Colquitt, D. P. Merullo, G. Konopka, T. F. Roberts, M. S. Brainard, Cellular transcriptomics reveals evolutionary identities of songbird vocal circuits. Science. 371 (2021), doi:10.1126/science.abd9704.

36. D. Hain, T. Gallego-Flores, M. Klinkmann, A. Macias, E. Ciirdaeva, A. Arends, C. Thum, G. Tushev, F. Kretschmer, M. A. Tosches, G. Laurent, Molecular diversity and evolution of neuron types in the amniote brain. Science. 377, eabp8202 (2022).

37. H. Norimoto, L. A. Fenk, H.-H. Li, M. A. Tosches, T. Gallego-Flores, D. Hain, S. Reiter, R. Kobayashi, A. Macias, A. Arends, M. Klinkmann, G. Laurent, A claustrum in reptiles and its role in slow-wave sleep. Nature. 578, 413–418 (2020).

38. M. A. Tosches, T. M. Yamawaki, R. K. Naumann, A. A. Jacobi, G. Tushev, G. Laurent, Evolution of pallium, hippocampus, and cortical cell types revealed by single-cell transcriptomics in reptiles. Science. 360, 881–888 (2018).

39. P. S. Ulinski, W. T. Rainey, Intrinsic organization of snake lateral cortex. Journal of Morphology. 165, 85–116 (1980).

40. M. A. Tosches, From Cell Types to an Integrated Understanding of Brain Evolution: The Case of the Cerebral Cortex. Annu Rev Cell Dev Biol. 37, 495–517 (2021).

41. C.-F. F. Chen, D.-J. Zou, C. G. Altomare, L. Xu, C. A. Greer, S. J. Firestein, Nonsensory target-dependent organization of piriform cortex. Proc. Natl. Acad. Sci. U.S.A. 111, 16931–16936 (2014).

42. Y. Chen, X. Chen, B. Baserdem, H. Zhan, Y. Li, M. B. Davis, J. M. Kebschull, A. M. Zador, A. A. Koulakov, D. F. Albeanu, High-throughput sequencing of single neuron projections reveals spatial organization in the olfactory cortex. Cell. 185, 4117–4134.e28 (2022).

43. E. Martin-Lopez, K. Ishiguro, C. A. Greer, The Laminar Organization of Piriform Cortex Follows a Selective Developmental and Migratory Program Established by Cell Lineage. Cereb Cortex. 29, 1–16 (2019).

44. C. Mazo, J. Grimaud, Y. Shima, V. N. Murthy, C. G. Lau, Distinct projection patterns of different classes of layer 2 principal neurons in the olfactory cortex. Sci Rep. 7, 8282 (2017).

45. J. Gerhart, M. Kirschner, The theory of facilitated variation. Proceedings of the National Academy of Sciences. 104, 8582–8589 (2007).

46. P. J. Wittkopp, G. Kalay, Cis-regulatory elements: molecular mechanisms and evolutionary processes underlying divergence. Nat Rev Genet. 13, 59–69 (2012).

47. J. Dugas-Ford, J. J. Rowell, C. W. Ragsdale, Cell-type homologies and the origins of the neocortex. Proc Natl Acad Sci U S A. 109, 16974–16979 (2012).

48. M. A. Tosches, G. Laurent, Evolution of neuronal identity in the cerebral cortex. Curr. Opin. Neurobiol. 56, 199–208 (2019).

49. J. Kaslin, J. Ganz, M. Brand, Proliferation, neurogenesis and regeneration in the non-mammalian vertebrate brain. Philos Trans R Soc Lond B Biol Sci. 363, 101–122 (2008).

50. F. Luzzati, A hypothesis for the evolution of the upper layers of the neocortex through co-option of the olfactory cortex developmental program. Front Neurosci. 9 (2015), doi:10.3389/fnins.2015.00162.

51. G. F. Striedter, R. G. Northcutt, The Independent Evolution of Dorsal Pallia in Multiple Vertebrate Lineages. Brain Behavior and Evolution. 96, 200–211 (2021).

52. S. Zeppilli, T. Ackels, R. Attey, N. Klimpert, K. D. Ritola, S. Boeing, A. Crombach, A. T. Schaefer, A. Fleischmann, Molecular characterization of projection neuron subtypes in the mouse olfactory bulb. eLife. 10, e65445 (2021).

53. F. A. Wolf, P. Angerer, F. J. Theis, SCANPY: large-scale single-cell gene expression data analysis. Genome Biology. 19, 15 (2018).

54. S. L. Wolock, R. Lopez, A. M. Klein, Scrublet: Computational Identification of Cell Doublets in Single-Cell Transcriptomic Data. Cell Systems. 8, 281–291.e9 (2019).

55. A. T. L. Lun, K. Bach, J. C. Marioni, Pooling across cells to normalize single-cell RNA sequencing data with many zero counts. Genome Biology. 17, 75 (2016).

56. V. A. Traag, L. Waltman, N. J. van Eck, From Louvain to Leiden: guaranteeing well-connected communities. Sci Rep. 9, 5233 (2019).

57. F. A. Wolf, F. K. Hamey, M. Plass, J. Solana, J. S. Dahlin, B. Göttgens, N. Rajewsky, L. Simon, F. J. Theis, PAGA: graph abstraction reconciles clustering with trajectory inference through a topology preserving map of single cells. Genome Biology. 20, 59 (2019).

58. Y. Zhang, T. Liu, C. A. Meyer, J. Eeckhoute, D. S. Johnson, B. E. Bernstein, C. Nusbaum, R. M. Myers, M. Brown, W. Li, X. S. Liu, Model-based Analysis of ChIP-Seq (MACS). Genome Biology. 9, R137 (2008).

59. K. Cao, Y. Hong, L. Wan, Manifold alignment for heterogeneous single-cell multi-omics data integration using Pamona. Bioinformatics. 38, 211–219 (2021).

60. P. Demetci, Q. H. Tran, I. Redko, R. Singh, Jointly aligning cells and genomic features of single-cell multi-omics data with co-optimal transport (2022), p. 2022.11.09.515883, doi:10.1101/2022.11.09.515883.

61. F. Pedregosa, G. Varoquaux, A. Gramfort, V. Michel, B. Thirion, O. Grisel, M. Blondel, A. Müller, J. Nothman, G. Louppe, P. Prettenhofer, R. Weiss, V. Dubourg, J. Vanderplas, A. Passos, D. Cournapeau, M. Brucher, M. Perrot, É. Duchesnay, Scikit-learn: Machine Learning in Python (2018), doi:10.48550/arXiv.1201.0490.

