## Supplementary Material for "Mammalian olfactory cortex neurons retain molecular signatures of ancestral cell types"

**The PDF file includes:**

Materials and Methods
Supplementary text: abbreviations
Fig.s. S1 to S24
References (52 to 61)

### Materials and Methods

#### Experimental model and subject details

A total of 12 adult C57Bl/6 mice (6 females and 6 males) of age between 6 and 8 weeks, 10 adult wild mice (5 females and 5 males) derived from mice trapped in the fields near livestock barns (Idaho, USA) and used as an outbred stock of pathogen-free wild mice ("wild-derived mice") of age between 7 and 8 weeks (*29*, *30*), and 3 adult Vglut1-Cre/INTACT-GFP transgenic mice of age between 6 and 8 weeks, were used in this study and obtained by in-house breeding. All animal protocols were approved by the Brown University's Institutional Animal Care and Use Committee (protocol number: 21-03-0004) followed by the guidelines provided by the National Institutes of Health, as well as approved by the Institutional Animal Care and Use Committee of the Weizmann Institute of Science.

#### Tissue microdissection and single-nuclei isolation

*Tissue microdissection*

Mice were deeply anesthetized with 2.5% of 250 mg/kg Avertin and transcardially perfused with 10 ml of ice-cold phosphate-buffered saline (PBS). The brains were dissected and immediately manually sliced into 500-700 µm coronal sections using the adult mouse brain slicer matrix (Zivic Instruments, BSMAS001-1). Anterior piriform cortex (aPir), posterior piriform cortex (pPir), agranular insular cortex (AI), and primary somatosensory cortex (SSp) were microdissected under a stereo microscope. With respect to the dorsolateral boundaries, the rhinal fissure was used as a visual landmark for the microdissections: about half millimeter below the rhinal fissure for aPir and pPir, right below the fissure for AI, and about half millimeter above for SSp. With respect to the anterior-posterior axis, based on the Paxinos and Franklin Mouse Brain Atlas, 1-3 coronal slices were cut within a 2.20mm to 0.14mm window from Bregma, and 1-2 coronal slices within a -1.06mm to -2.06mm window from Bregma. aPir and AI were microdissected from anterior slices, while pPir and SSp from posterior slices. Both hemispheres were included for each cortical area. One slice in between aPir and pPir dissections was always removed to avoid piriform anterior-posterior border inconsistency across mice and strains. The remaining tissue was fixed overnight at 4°C in 4% paraformaldehyde (PFA) for *post-hoc* histological validation of the microdissected areas **(see histology method section and figs. S1 and S14)**. Potential cells from neighboring regions were identified in the transcriptomic analysis and removed from the datasets (see data pre-processing, QC, normalization and clustering method section).

28 individual biological replicates from lab mice and 12 individual biological replicates from wild-derived mice were sequenced: 9 (anterior piriform), 10 (posterior piriform), 5 (agranular insular) and 4 (primary somatosensory) replicates from lab mice, and 6 (anterior piriform), 4 (posterior piriform) and 2 (primary somatosensory) replicates from wild-derived mice.

*Single-nuclei isolation*

We isolated single nuclei suspensions from fresh tissue by adapting previously described procedures in Zeppilli et al., 2021 (*51*) for ATAC sequencing (seq) experiments. Each biological replicate was minced separately and placed into a tube containing cold Nuclei PURE Lysis Buffer and 10% Triton X-100 (Sigma, NUC201-1KT). The minced tissue was transferred into a 7 ml ice-cold tissue grinder (Sigma, D9063), homogenized up and down 20-25 times, and filtered on ice through cell strainers of 100µm, 70µm and 40 µm (Pluriselect, 43-10040). RNasin Plus diluted 1/200 (Promega, N2611) was added in all solutions.

*Single-nuclei isolation for multiome sequencing experiments*
After centrifuging at 500 rpm for 5 min at 4°C, the supernatant was aspirated and gently resuspended in 100 µl of a cold lysis solution containing Nuclei PURE Lysis Buffer, 10% Triton X-100, 0.01% of digitonin (ThermoFisher, BN2006), and 1% nuclease-free UltraPure™ BSA (ThermoFisher, AM2616). After 1 minute of incubation, 200 µl of cold 1X Nuclei Buffer (10x Genomics, 2000153/2000207) was added, the suspension was filtered again with a 40 µm cell strainer, centrifuged at 500 rpm for 5 min at 4°C, and gently resuspended into a final volume of 50 µl of cold 1X Nuclei Buffer.

*Single-nuclei isolation for RNA sequencing experiments*
After centrifuging at 500 rpm for 5 min at 4°C, the supernatant was aspirated and gently resuspended in 400 µl of a cold wash solution containing 1X Hanks' Balanced Salt Solution HBSS and 1% nuclease-free UltraPure™ BSA. This step was repeated for a total of two times, and nuclei were resuspended into a final volume of 50 µl of cold wash solution.

**Library preparation and single-nucleus ATAC and RNA sequencing**
*Library preparation*
Single-nuclei libraries were generated using the Single Cell Multiome ATAC + Gene Expression kit (10x Genomics, PN-1000283). Manufacturer's instructions were followed for *Tn5*-based transposition, cell capture, barcoding, reverse transcription, cDNA amplification, and ATAC and RNA libraries construction. For wild-derived mice, 4 of 6 biological replicates for aPir, 1 of 4 for pPir, and 1 of 2 for SSp were processed using the Single Cell 3' Reagent Kits v3.1 dual index (10x Genomics, PN-1000268). Final libraries (40 libraries from gene expression and 32 libraries from ATAC experiments) were evaluated for quality and quantified using the Qubit fluorometer. The fragment size distribution was evaluated by Agilent TapeStation 2200 (Agilent Technologies). Libraries were further evaluated for proper incorporation of the Illumina adaptors on the Roche LightCycler 480 using the Roche Kapa library quant assay according to manufacturer's protocol.

*Sequencing*
Libraries were pooled and sequenced on an Illumina NovaSeq6000 instrument. ATAC libraries were sequenced using the read lengths of R1: 50 bp R2: 49 bp I1: 8 bp I2: 24 bp. RNA libraries were sequenced using the read lengths of R1: 28 bp R2: 90 bp I1: 10 bp I2: 10 bp. The mean raw read pairs per cell achieved across the 72 libraries were 295,832.59 reads/nucleus for the Multiome GEX libraries, 335,692 reads/nucleus for the Multiome ATAC libraries, and 54,838.75 reads/nucleus for the Single Cell 3' v3.1 gene expression libraries.

**Pre-processing, quality control, normalization and clustering of RNA and ATAC data**
*Genome alignment*
The generated FASTQ files were processed with Cell Ranger ARC (v2.0.0) for Multiome ATAC + Gene expression experiments and with Cell Ranger (v6.0.0) for Single Cell 3' v3.1 experiments (10x Genomics). Reads were aligned to the mouse (*Mus musculus*) pre-mRNA reference genome (cellranger-arc-mm10-2020-A-2.0.0) for both lab and wild-derived mice.

*Transcriptome analysis*
Individual biological replicates of all cortical areas were merged into a single dataset from the start (from pre-processing) for each mouse strain, to provide a consistent reference for comparing gene

expression and chromatin accessibility of neurons across mouse cortical areas. The main two datasets were the lab dataset (which includes only 10x Multiome experiments from all areas) and the wild dataset (which includes 10x Multiome and Single Cell 3' v3.1 experiments from all areas). Scanpy (v1.8.2) was used to perform most of the transcriptomic analysis (*52*). Only nuclei that had between 2,000 and 10,000 genes per nucleus (average 4,874 and 2,900 genes/nucleus for lab and wild datasets, respectively), and a percentage of mitochondrial counts below 2.5% were retained (**figs. S2 and S15**). Note that more glia cells than neurons were filtered out as glia cells contain less number of genes per nucleus. Genes expressed in less than 7 nuclei were also removed. Doublets were detected using Scrublet (v0.2.3) with default parameters (*53*). After quality filtering, lab and wild datasets comprised 7,840 and 26,975 high-quality nuclei, respectively. Processing for each dataset consisted of normalization of the expression matrix using the R package SCRAN (v3.12) called from Python with default parameters (*54*), identification of the top 2,000 highly variable genes (HVG) amongst replicates, regression of the percentage of mitochondrial content and number of counts, and scaling of the expression values per replicate. Principal Component Analysis (PCA) linear dimensionality reduction was then performed on the scaled data, and the first 200 principal components (PCs) were selected for the generation of a k-nearest-neighbor (knn) graph. The knn graph served as input for unsupervised clustering using the graph-based Leiden algorithm (*55*), and for visualization in low dimensional spaces using Uniform Manifold Approximation and Projection (UMAP) or Partition-based graph abstraction (PAGA) (*56*). PAGA plot generates a topology-preserving map of groups of cells whose nodes correspond to the groups (clusters) and weighted edges to the transcriptomic relationship between groups (clusters). To correct for batch effects, we used Harmony on the selected PCs (200) (*32*). For both lab and wild-derived datasets, we performed a first coarse clustering and identified potential cells from neighboring areas using differential expression (DE) analysis and available RNA in situ hybridization (ISH) data from the Allen Brain Atlas. We located these neighboring cells primarily to endopiriform nucleus/claustrum based on a combination of genes such as *Npsr1, Rorb, Rspo2, Fezf2, Nr4a2, Slc26a4, Reln* and *Pou6f2*. These genes were not expressed together in clusters clearly identified as piriform cells using other established markers (for example *Reln* combined with *Rorb* for SL cells). After the removal of 'neighboring cells', we repeated all the computational steps above, excluding quality filtering. Leiden clustering was run using the following neighbors and resolution parameters: 50 - 2.5 (coarse, lab dataset) and 50 - 1.5 (coarse, wild dataset). In both datasets, we identified classes of neuronal and non-neuronal cells. These classes included mature neurons (*Syt1, Syn1, Rbfox3, Slc17a7, Gad1, Gad2*), immature neurons (higher levels of *Dcx*, *Sox4, Sox11* compared to mature neurons), microglia (*Tmem119, Siglech*), oligodendrocytes (*Mog, Mbp, Mobp*), oligodendrocyte precursors (*Pdgfra*), astrocytes (*Gfap, Sox9, Slc6a11*) and vascular leptomeningeal cells (*Vtn, Dcn, Egfl7*) **(figs. S3 and S16).** Mature and immature neurons were further sub-clustered by repeating all the computational steps above, excluding quality filtering. For sub-clustering, we used neighbors and resolution parameters: 50 - 3.7 (neurons, lab dataset) and 50 - 2.9 (neurons, wild dataset). This resulted in 5,553 high-quality nuclei grouped in 27 supertypes (lab dataset), and 24,901 high-quality nuclei grouped in 36 supertypes (wild dataset). Supertypes are defined as clusters derived from the unsupervised clustering of neurons from all cortical areas. A supertype may include neurons from all areas or neurons from only some areas. While the number of nuclei was higher in the wild dataset due to the additional use of the Single Cell 3' v3.1 kit (Single Cell 3' v3.1 kit enables to input a higher volume of single-nucleus suspension), the fundamental cellular components did not differ between the two datasets (**fig. S16C**).

*Epigenome analysis*

The corresponding ATAC seq data of the lab dataset (multiome sub-clustered dataset, which includes only mature and immature neurons) were processed using pycisTopic (v1.0.2.dev8+g848f78b) (*20*). Note that the ATAC data of the wild dataset (multiome replicates) were not processed within the context of this study, thus hereafter we refer only to the lab dataset for the epigenomic analysis. Nuclei with > 3.5 log unique fragments per nucleus (average 53,470 fragments/nucleus), FRIP > 0.2, and TSS enrichment > 4.0 were retained, resulting in 5,190 high-quality nuclei that passed both transcriptome- and epigenome-specific quality control (QC) metrics (**fig. S2**). For each cortical area, we downsampled cell numbers to the same number by maintaining the original supertype proportions to avoid interpretations based on asymmetric data. The downsampled dataset comprised 3,430 nuclei from all areas. In all analyses, we used both the original and downsampled dataset to verify robustness of the results. In-depth analyses were only performed on the downsampled dataset. Next, we generated pseudo-bulk ATAC seq datasets by combining, for each cortical area, fragment reads for the transcriptome-based neuronal clusters. Peak calling was performed on these pseudo-bulk data using MACS2 (v2) (*57*) with default parameters: shift=73, ext_size=146, keep_dup='all', q_value=0.05. To generate a list of consensus peaks, we used pycisTopic's iterative peak calling algorithm, which resulted in 479,584 chromatin accessibility regions across the four cortical areas. We used Mallet (v2.0.8) for topic modeling through Latent Dirichlet Allocation (500 iterations). Models were selected based on the stabilization of Arun_2010, Cao_Juan_2009, Minmo_2011 and log likelihood quality metrics. A single model of 50 topics (without downsampling) and 75 topics (with downsampling) was selected for cortical areas together, and area-specific models of 32, 32, 35, and 37 topics were selected for aPir, pPir, AI, and SSp, respectively (all with downsampling). Batch correction was applied on scaled topic distributions using the Python implementation of Harmony (*32*), and clustering was performed using the Leiden algorithm (*55*) with neighbors and resolution parameters 50 and 2.0, respectively. Differentially Accessible Regions (DARs) were calculated between cortical areas, between supertypes within and across cortical areas, and between cortical layers within each cortical area, using default parameters: adjpval_thr=0.05, log2fc_thr=1.0. Next, to enable transcription factor (TF) binding motif predictions on the ATAC seq data, custom motif rankings and scores databases were generated using the protocols provided on https://github.com/aertslab/create_cisTarget_databases. Using these databases, motif enrichment analysis was performed on the different sets of DARs and on binarized topics (Otsu thresholding) using the cistarget and DEM methods of pycisTarget (v1.0.2.dev8+g48af509.d20220905); with http://sep2019.archive.ensembl.org as biomart host, motif annotations v10nr_clust (public version), ctx_auc_threshold=0.005, ctx_nes_threshold=3.0, ctx_rank_threshold=0.05, dem_log2fc_thr=1.0, dem_motif_hit_thr=3.0, and dem_max_bg_regions=500.

**Supertype annotation and quantification across cortical areas**

*Supertype annotation*

We used a piriform-based cell type taxonomy across cortical areas. We assigned supertype labels based on the expression of well-established marker genes, such as *Reln, Cux1, Bcl11b Ctip2*), as well as based on genes identified through DE analysis. DE analysis on piriform lab data (raw data) was performed in scanpy (*52*) using the functions *tl.rank_genes_groups* with method t-test, penalty L2, and *tl.filter_rank_genes_groups* based on minimum log-fold-change of 3. *In situ* hybridization

(ISH) images from the Allen Brain Institute and immunohistochemical experiments for the DE genes were then used to attribute piriform layer specificity.
For SSp, we further matched the piriform-based supertype labels with the standard SSp nomenclature based on the Allen Brain Institute as described in the method section below named 'integration of single cell RNA sequencing datasets from *this study* and a mouse single-cell reference atlas'.

*Supertype quantification across cortical areas*
For quantification across cortical areas: for each cortical area, relative contributions to each supertype were quantified by normalizing the number of nuclei of a single area to 1. This made supertypes comparable between areas for quantification. We defined enrichment of supertype $s$ in area $a$ with respect to area $b$ as the ratio between normalized $s_a$ / normalized $s_b$ > 1.75. For visualization in PAGA plots with pie charts representing the (relative) contributions of each area to a supertype, we took per-cortical-area normalizations of a supertype and re-normalized these to add up to 1 (a full pie chart). Within cortical areas, we quantified the contribution of supertypes with respect to cortical layer assignments using fractions. E.g. in SSp, L4 is assigned to SL1, Pyr 1-2-3-4-14-15. Thus, the contribution of SL1 to L4 is 7%, which is 22 / (22 + 80 + 54 + 54 + 40 + 32 + 22). Similarly, the contribution of Pyr neurons in supertypes shared across the four cortical areas was computed (per cortical area) as the fraction of shared-supertype Pyr neurons over the total number of Pyr neurons (in shared and area-specific supertypes).

**Integration of single-cell RNA sequencing datasets from *this study* and a mouse single-cell reference atlas**
*Integration between SSp datasets*
We integrated our SSp dataset (from the lab dataset) with the SSp dataset from a single-cell reference atlas (*17*), whose cell types also included connectivity- and layer-specific information. We integrated the two SSp datasets using the R package Seurat (v4.3) (*16*). After subsetting to only SSp cells for each dataset, the two subsets were normalized using Seurat's *SCTransform* v2 function, which corrects for differences in sequencing depth. Each SSp subset was regressed by the number of counts. The two subsets were then merged together in a list class object, from which the integration features were calculated. We computed 2000 HVGs. Pairs of mutual nearest neighbors (anchors) were identified using the *FindIntegrationAnchors* function with the arguments reduction="CCA" (canonical correlation analysis) and normalization method= SCT. Integration was carried out with the Seurat function *IntegrateData*, normalizing with SCT using the anchor sets and 200 PCs. Dimensionality reduction was performed by calculating PCA with 200 PCs. Unsupervised clustering was performed with the default method using the SLM algorithm and clustering parameters 25 (neighbors), 3 (resolution). The resulting integrated object was then visualized in a low dimensional space using the UMAP algorithm. New clusters that result from the unsupervised clustering of the two integrated datasets, also referred to as integrated clusters, were used to assess the co-clustering between the two datasets and to assign the neuron type identity to the SSp clusters of our dataset (**fig. S6B**).

*Integration between all mouse cortical areas*
We integrated our entire lab dataset (aPir, pPir, AI, and SSp) with the mouse single-cell reference atlas (*17*), whose cell types also included connectivity- and layer-specific information. We used the R package Seurat (v4.3) (*16*), and the same computational pipeline and parameters used for the

integration between the SSp datasets described above. The reference atlas was subsampled to 25,000 cells from the original dataset and included glutamatergic and inhibitory neurons from the entire mouse cortex, excluding piriform and cortical amygdala. Areas included the hippocampal formation, including subiculum and retrohippocampal regions, the neocortex, including primary and secondary motor, visual, somatosensory and auditory cortex, and transition areas, including lateral and medial entorhinal cortex. New clusters that result from the unsupervised clustering of the integrated datasets, also referred to as integrated clusters, were used to assess and quantify the co-clustering between piriform and neocortical glutamatergic neurons, and between piriform glutamatergic neurons and glutamatergic neurons from transition areas (ENT, ENTl, ENTm, TPE-ENT) and hippocampal formation areas (Sub, Sub-ProS, PPP, RHP, DG, CA1-2-3). The quantification was based on integrated clusters having at least 8 cells from both piriform and the other areas (i.e. minimum cluster size 16). The fraction of piriform glutamatergic neurons co-clustering with the other neurons is then the sum of piriform cells in the integrated clusters divided by the total number of piriform cells in the integrated object.

**Inference of enhancer-driven Gene Regulatory Networks (e-GRNs)**

*e-GRNs computation*

To infer e-GRNs across cortical areas, we applied SCENIC+ (*20*) on the lab dataset (on the sub-clustered and downsampled dataset as described in the method section epigenome analysis). We computed e-GRNs for each cortical area by including glutamatergic and inhibitory neurons (INs), namely SL, Pyr, Vglut2 and IN cells. Immature neurons were excluded to restrict our analysis on mature neurons. We used a genomic search region around genes of +/-500kb, http://sep2019.archive.ensembl.org as biomart host, and otherwise default parameters. Promoter regions were excluded from the analysis as they are ubiquitously open and tend to provide little discriminatory information (*20*). SCENIC+ identifies e-regulons, which consist of TFs, their target enhancers, and their downstream target genes. High quality e-regulons were selected by keeping those with a correlation >0.4 between the areas under the curve (AUC) of target gene activity and target enhancer activity, and with ≥10 number of target genes. Given inhibitory neurons were transcriptomically and epigenetically well-conserved across cortical areas, they were used as an internal control. e-Regulons specific for inhibitory neurons were determined using Regulon Specificity Score (RSS) and removed for analyses specific to glutamatergic neurons. IN-specific e-regulons were Arx(+), Dlx1(+), Dlx2(+), Dlx5(+), Dlx6(+), Lhx6(+), Maf(+), Mafb(+), and Sox2(+), where (+) indicates an activating role for the e-regulon's TF.

*Transcription factor binding site motifs*

To assess which binding motifs were identified for each TF, we ran pycisTarget on cistromes of each e-regulon (*20*). We then ordered the resulting binding motifs by their Normalized Enrichment Score (NES), a metric that captures how enriched a motif is in comparison to the average presence of all motifs. Only TFs (both e-regulon and non-e-regulon TFs) expressed in at least 20% of neurons belonging to a particular cortical layer were considered. We then examined the number of CREs in which the binding motif was found in relation to the total number of CREs for the TF. For the shared e-regulons, we compared across area-specific e-GRNs which binding motif was the top motif. We visualized these motifs for selected e-regulons.

*Transcription factor combinations*

For each area-specific e-GRN, we quantified TF combinations for each target gene. We grouped all CREs of a gene and interrogated them for the TFs that are capable of binding at these genomic sites. Next, in a pairwise fashion, TF-TF combinations were counted and visualized using clustered heatmaps. We performed this analysis both for shared e-regulons and for all e-regulons of an e-GRN. Moreover, single genes were inspected for differential accessibility of their coding sequence and their surrounding chromatin, including for differential importance of their CREs.

*Quantification of supertype distances (discreteness)*
To determine if cell types tend to form a graded or a more discrete molecular identity, we reasoned that the minimal distance between two clusters should be smaller for graded cell types than for discrete cell types. We thus quantified and compared cluster discreteness between cortical areas by including only supertypes composed of glutamatergic neurons from all areas. We performed this analysis at two levels: on transcriptomically similar neurons from all cortical layers, and selectively on transcriptomically similar neurons within a particular layer. The within-layer analysis makes sure results are not based solely on differences between layers. We also used the downsampled datasets as done in the ATAC analysis to avoid interpretations based on asymmetric datasets. We first re-processed the transcriptome of the downsampled datasets. Next, we defined the distance between clusters as the minimal pairwise euclidean distance between supertypes for each area, using their PCA representation (number of PCs = 200). As this measure may be sensitive to outliers for each cortical area and for all pairs of cell types of interest, we sampled 100 times 90% of the cells in each cluster and computed the distribution of cluster distances. We visualized results with kernel-density-estimate plots using kdeplot from Seaborn (v0.11.2) and we confirmed observations of one distribution being less than another using Mann-Whitney rank tests. Across all cortical layers: aPir < pPir: $p = 0.999$; aPir < SSp: $p = 3.571e-109$; pPir < SSp: $p = 5.785e-206$; AI < SSp: $p = 1.0$. Within the single layer Pir 2b/3, SSp L2/3: aPir < pPir: $p = 1.0$; aPir < SSp: $p = 3.501e-5$; pPir < SSp: $p = 5.233e-124$; AI < SSp: $p = 0.459$.

*Repression networks and repression quantification*
We quantified and compared transcriptional repression across cortical areas by quantifying activating and repressive interactions in each area-specific e-GRN. Area-specific supertypes were excluded to better compare predicted regulatory interactions within transcriptomically similar neurons (supertypes SL1, Pyr 11-13-14-15-16, and immature neuron were excluded). Using for each network its high-quality e-regulons, a percentage of repressive interactions was calculated using the formula 100 * *repr / (act + repr)*, where *act* indicated activating TF-TF interactions and *repr* repressive ones. Only TFs (both e-regulon and non-e-regulon TFs) expressed in at least 20% of neurons belonging to a particular cortical layer were considered. Non-e-regulon TFs could only be the recipient of a regulatory interaction. For visualization and to understand how repressive interactions were distributed across the e-GRN, we focused on cortical layers. Layer-specific e-regulons were determined using the Regulon Specificity Score (RSS) and by keeping only e-regulons with their TFs expressed in >10% of glutamatergic neurons. Note that we did not restrict TF expression to the layer for which we computed specificity, since repressors tended to be expressed outside of the layer where their target genes were expressed. Due to lack of layer information on AI, we omitted this cortical area. The union of layer-specific e-regulons resulted in networks of size 53, 51, and 51 e-regulons, for aPir, pPir and SSp respectively. From these networks, we visualized the subgraph of all repressing e-regulons and their interactions as

repressing. Placement of each TF along cortical depth was determined by the center of their expression domain along the cortical layer axis and by network layout constraints.

**Integration of single-cell RNA sequencing data of lab and wild-derived mice using optimal transport**

To compare the degree of similarity between lab and wild-derived mice across cortical areas (aPir, pPir, and SSp), we aligned gene expression data in a shared computational space and quantified the degree of overlap between lab and wild datasets. All computational steps are carried out separately for each cortical area.To perform the alignment, we used an optimal transport (OT) framework (see method section below), treating single-cells measurements as probability distributions. The method first finds cell-to-cell correspondence probabilities between lab and wild datasets, and then co-embeds the two datasets in a shared space based on these probabilities. Finally, we calculated a 'conservation score between the co-embedded lab and wild datasets (see method section below). We used inhibitory neurons (INs) as internal control as being transcriptomically and epigenetically well-conserved across cortical areas compared to glutamatergic neurons. We also excluded immature neurons from this analysis as they are hypothesized to be the potential source of variation.

*Finding cell-cell similarities using optimal transport*

Optimal transport is a mathematical framework for matching probability distributions or discrete measures to one another. It has been previously employed to align single-cell datasets of different genomic measurement types (*58*, *59*). Here, we leveraged it to align gene expression data from lab and wild-derived mice. We treated the two count matrices, $X = [x_1, \ldots, x_n]^T \in R^{n \times d}$ and $X' = [x'_1, \ldots, x'_{n'}]^T \in R^{n' \times d'}$, where $n$ and $n'$ are the number of cells and, $d$ and $d'$ are the number of features (genes), as discrete measures by defining probabilistic weights over both the cells and the features. Here, $R$ represents real values. We defined the weights associated with the cells as uniform distributions $w = [w_1, \ldots, w_n]^T$ *and* $w' = [w'_1, \ldots, w'_{n'}]^T$ for the lab and wild datasets, respectively, where $w_1$=...=$w_n = 1/n$ and $w'_1$=...=$w'_n = 1/n'$. Similarly, we defined the uniform weights associated with the genes $v = [v_1, \ldots, v_d]^T$ *and* $v' = [v'_1, \ldots, v'_{d'}]^T$, where $v_1$=...=$v_d = 1/d$ and $v'_1$=...=$v'_d = 1/d'$. These weights can be seen as the histograms of probabilities for observing a cell or gene, and by initializing them as uniform distribution, we assumed no prior knowledge, as has been done by other optimal transport applications on single-cell datasets (*58*, *59*). Given these distributions and the count matrices, we used a specific optimal transport framework, called the unbalanced co-optimal transport (UCOOT) algorithm (*31*), to jointly solve for two probabilistic alignment matrices – $\pi^s \in [0,1]^{n \times n'}$ between the cells, and $\pi^f \in [0,1]^{d \times d'}$ between the genes that will attain:

$$min_{\pi^s \epsilon \Pi^s, \pi^f \epsilon \Pi^f} \quad \sum_{i,j,k,l} \quad \left(X_{i,k} - X'_{j,l}\right)^2 \pi^s_{i,j} \pi^f_{k,l} \qquad (+ \text{ regularization term})$$

such that $\Pi^s$={$\pi \epsilon\ R_+^{n \times n'} \quad \cup\ 0$ } $\Pi^f = \{\pi \epsilon\ R_+^{d \times d'} \quad \cup\ 0\}$

Intuitively, UCOOT finds the two alignment maps, $\pi^s$and $\pi^f$, that will minimize the sum of squared error between the two count matrices when the cell and the gene space of one dataset is transformed into the other according to these maps. Our motivation behind picking this specific optimal transport formulation was to leverage the feature alignment matrix, $\pi^f$, to provide supervision on shared features (genes) and to improve the quality of dataset alignment. We used the shared genes between the lab and wild datasets to fix the entries in the feature alignment map that correspond to these features, enforcing their alignment, which also informs cell alignments

through the joint formulation defined above. Following other works in optimal transport literature (*31*, *58*, *59*), we added two regularization terms to the objective function. The first is an entropic regularization term $\varepsilon_1 H(\pi^s) + \varepsilon_2 H(\pi^f)$, where $H(\pi) = -\sum_{i,j} \pi_{ij} log(\pi_{ij})$ is the Shannon entropy. This entropic regularization term is commonly used in practical applications of optimal transport as it leads to more robust and fast estimation of mappings while preventing them from being too sparse. Additionally, we included a "weight relaxation" term:
$\lambda_1[KLD(\pi^s 1_n|w) + KLD(n^{s^T} 1_{n'}|w')] + \lambda_2[KLD(\pi^f 1_d|v) + KLD(n^{f^T} 1_{d'}|v')]$,
where $1_n$ is a n-length vector of ones, KLD is the Kullback-Leibler divergence, quantifying the difference between the input probabilistic weights defined over the cells and features and the weights distributed in the alignment maps. Practically, this relaxation term allows for diverging from the uniform weight distributions, so weights could be locally adapted in alignment, and smaller probabilities could be distributed over dataset-specific cells that do not have well-matching correspondences in the other dataset. With these relaxation terms, the full objective function becomes:

$$min_{\pi^s,\pi^f \epsilon \Pi} \sum_{i,j,k,l} \left(X_{i,k} - X'_{j,l}\right)^2 \pi^s_{i,j}\pi^f_{k,l} + \varepsilon_1 H(\pi^s) + \varepsilon_2 H(\pi^f) + \rho_1[KLD(\pi^s 1_n|w) + KLD(n^{s^T} 1_{n'}|w')] + \rho_2[KLD(\pi^f 1_d|v) + KLD(n^{f^T} 1_{d'}|v')],$$

with the entries of alignment matrices constrained to be non-negative. The terms $\varepsilon_1, \varepsilon_2, \rho_1$, and $\rho_2$ are hyperparameters. We defined the following grids over these$\varepsilon_1, \varepsilon_2 = [0.005, 0.001, 0.05, 0.01, .., 0.1]$ , $\rho_1, \rho_2 = [0.005, 0.001, 0.05, 0.01, .., 1]$, and then pick the hyperparameter combination that leads to the lowest non-zero sum of squared difference between the aligned count matrices (zero leads to a degenerate solution of empty alignment matrices).

*Co-embedding datasets based on cell-cell similarities:*
For each cortical area, we integrated the gene expression data from lab and wild-derived mice into a shared space by co-embedding them according to their alignment probabilities obtained from the optimal transport step. To do so, we followed the procedure by Cao *et al.* (*58*) and first constructed a weighted k-nearest neighbor (k-nn) graph for each dataset, where the nodes are cells and edges are weighted by the pairwise Euclidean distance between cells in the same dataset. Then, we computed the Laplacian matrices, $L_X$ and $L_{X'}$ where $L_X = D_X - A_X$, $D_X$ is the diagonal matrix of node degrees, and $A_X$ is the adjacency matrix of the k-nn graph of the dataset $X$. The Laplacian matrices take into account each dataset's original structure. When co-embedding datasets, we not only want to position cells with high alignment probabilities close together, but we also want to keep cells that were close to each other in the original dataset nearby. Given these Laplacian matrices $L_X$, $L_{X'}$, the cell alignment map $\pi^s$ , and the count matrices $X, X'$, we solved for the two co-embedded datasets in $p$-dimensional space, $X^e \epsilon R^{n \times p}, X'^e \epsilon R^{n' \times p}$, that will attain

$$max_{X^e, X'^e} \ tr(X^e \pi^s X'^{eT} ),$$
such that $X^e S_{XX} X^{eT} = I$ and $X'^e S_{X'X'} X'^{eT} = I$ ,

where $S_{xx} = L_x + \lambda \Sigma_x$ and $S_{x'x'} = L_{x'} + \lambda \Sigma_{x'}$ , $\Sigma_x = diag(\pi^s 1_n)$ and $\Sigma_{x'} = diag(1_{n'}{}^T \ \pi^s)$, $tr(\cdot)$ denotes the trace of matrix and $diag(\cdot)$ denotes a diagonal matrix. Similarly to Cao *et al.*, we carried out the optimization using the eigenvalue decomposition method. This process has three hyperparameters: $k$, $\lambda$, and $p$. We defined a grid over these hyperparameters $k \epsilon [5, 10, 15, 20, 25, 30]$, $\lambda \epsilon [0.001, 0.005, 0.01, 0.05, \ldots, 10]$, $p \epsilon [2, 3, 5, 10, 15, 20, 25, 30]$. Among these, we selected the hyperparameter combination that led to the highest dataset mixing (conservation score defined in the following section) of INs, as we observed the strongest alignment probabilities to be assigned between INs and determined them to be an "out group".

*Conservation score*
To compare the extent of similarity between lab and wild datasets across cortical areas, we quantified the overlap between the two datasets in the shared space. We expect neurons of more conserved cortical areas to be well-mixed, and neurons of less conserved cortical areas to be less mixed. To quantify the level of overlap, we defined a metric inspired by the local inverse Simpson's index (iLISI) used by the batch correction method, Harmony (*32*). Korsunsky *et al.* use iLISI to quantify the level of mixing between different batches after batch correction. Similar to their method, we built a Gaussian kernel-based distribution of neighborhoods, with a fixed perplexity (a bandwidth setting) parameter of 30. Then, for each cell from the lab dataset we computed $p(w)$, and for each cell from the wild dataset we computed $p(l)$. These values correspond to the probability of finding a neighbor from the opposing (wild or lab) dataset in the local neighborhood of a cell. We computed the mixing probability score as: $1/\sum \quad p(l)^2\, p(w)^2$. This score differs from the iLISI scores. Our motivation behind this change is the following observation: iLISI scores are low if a cell's neighborhood is mostly composed of one dataset, indicating poor mixing. However, iLISI score does not differentiate between whether these neighbors are from the cell's own dataset or the opposite dataset. While cells locally co-embedded with other cells from their own dataset indicate low level of conservation (and potentially dataset-specific cells), the opposite (i.e. cells mostly co-embedded with cells from the other dataset) does not necessarily indicate a lack of conservation. Therefore, to account for this difference, we computed the probability of having a neighbor from the opposite dataset. Furthermore, to take dataset size imbalance into account, we downsampled the number of cells in the wild dataset to the number of cells of the lab dataset. We maintained the original supertype proportions in the downsampling. We repeated this analysis 20 times and computed the dataset mixing probabilities for each cell in the shared space. Then, we reported the median probability as the cortical area conservation score, along with its 95% confidence intervals from the 20 downsampling repeats. We additionally reported the conservation score per main cell types (SL, Pyr, IN and Vglut2). Confidence interval (CI) of conservation scores for main cell types were: aPir SL (95% CI: 0.400-0.422), Pyr (95% CI: 0.405-0.439), Vglut2 (95% CI: 0.417-0.438), IN (95% CI: 0.456- 0.494); pPir SL (95% CI: 0.417-0.439), Pyr (95% CI: 0.378-0.433), Vglut2 (95% CI: 0.445-0.461), IN (95% CI: 0.461-0.489); SSp SL-like (95% CI: 0.439-0.467), Pyr (95% CI: 0.456-0.489), Vglut2 (95% CI:0.433-0.467), IN (95% CI: 0.483-0.511).

**Single-cell Variational Inference (scVI) integration between lab and wild pPir datasets**
Gene expression data from lab and wild pPir datasets were integrated using scVI-tools (v0.17.3), and the Single-cell Variational Inference (scVI) model (*60*). In brief, the top 3000 high variable genes (identified via the *scvi poisson_gene_selection* function) were used to train an scVI model with default parameters (n_hidden=128, n_latent=10, n_layers=1, dropout_rate=0.1) and a negative-binomial gene likelihood. Each biological replicate and mouse strain were encoded as categorical covariates and the log-total counts were used as a continuous covariate. The model was trained for 200 epochs until the ELBO converged. To evaluate whether neuronal types (SL, Pyr, INs, Vglut2) were aligned in this shared latent space, the pairwise cosine distances were calculated for each cell and then averaged depending on the neuron type and mouse strain of the cells in each pair. To evaluate whether OT-identified misaligned neurons were also less aligned in the scVI

latent space, a nearest neighbor graph was constructed for n=5–100 neighbors and, at each value of n, the average percent of neighbors that were from lab mice were calculated for both wild-specific neurons compared to other aligned Pyr neurons.

*Linear Support Vector Classifier*
Classification was performed on the log-normalized gene expression data from both lab and wild pPir datasets using scikit-learn (v1.3.0) (*61*). A classification pipeline was constructed to distinguish between Pyr, SL, Vglut2, IN_CGE, and IN_MGE neurons by identifying the top 100 genes (*SelectKBest* with k=100), z-scoring their expression, reducing the dimensionality of the data with PCA, and fitting a linear Support Vector Classifier (SVC) with the regularization parameter C=0.1 on the top 25 PCs. Immature neurons were excluded from the model training. These models were fit with stratified k-fold (k=5) cross-validation, such that the entire pipeline (gene selection, scaling, PCA, and classifiers) was only fit using the training data. Classification performance was evaluated on held-out cells. The models were fit by subsampling equal numbers of cells (250) for each of the above neuron types, and this procedure was repeated 100 times. To evaluate which of the above neuron types were most similar to immature neurons, the fitted models on each restart were then applied to the immature cells and the percentage of immature cells that were predicted to be each neuron type were recorded. Classification and generalization performance were at chance levels when classifiers were trained on data with permuted neuron type labels.

**Integration of single-cell RNA sequencing datasets across species**
*Orthologous gene alignment*
One-to-one orthologues were used to determine which genes were to be included for the cross-species comparison. EggNOG orthology assignments for lizard (*Pogona vitticeps*) and turtle (*Trachemys scripta*) were taken from (*37*), whereas salamander (*Pleurodeles waltl*) and mouse (*Mus musculus*) were taken from (*19*).

*Cortical areas and species included in the cross-species comparison*
We integrated single cell RNA sequencing (sc-RNA seq) data from lizard *Pogona vitticeps* (*35*, *36*), turtle *Trachemys scripta* (*37*), salamander *Pleurodeles waltl* (*19*) and mouse *Mus musculus* (*this study* - lab dataset; (*17*)). We subsetted the original datasets to include only cells from ontogenetically equivalent brain regions. The lizard dataset included glutamatergic neurons from medial cortex (MCtx), dorsal cortex (DCtx), lateral cortex (LCtx), anterior dorsal ventricular ridge (aDVR), and inhibitory neurons (INs). Note for the nomenclature: the reptilian cortex includes two hippocampal regions, called medial and dorsomedial cortex. In lizard, clusters from the hippocampus could be identified, but were not mapped precisely to these hippocampal subdivisions, therefore we use here the term MCtx to annotate cells of both subdivisions. For the turtle dataset, we used the same approach described in (*19*), which in short is keeping glutamatergic and INs, and excluding unidentified clusters. The salamander dataset included glutamatergic neurons from medial (MP), dorsal (DP), lateral (LP) and ventral (VP) pallia, and INs. The mouse dataset included glutamatergic and inhibitory neurons from the lab dataset of *this study* (including aPir, pPir, AI and SSp), and glutamatergic and inhibitory neurons from Yao et al., 2021 (*17*). The latter was subsampled to 25,000 cells from the original dataset and included glutamatergic and inhibitory neurons from the entire mouse cortex, excluding piriform and cortical amygdala. Areas included the hippocampal formation, including subiculum and retrohippocampal regions, the

neocortex, including primary and secondary motor, visual, somatosensory and auditory cortex, and transition areas, including lateral and medial entorhinal cortex.

*Cross-species integration*
Only one-to-one orthologs in all species analyzed were used for cross-species comparisons. We used the R package Seurat (v4.3) (*16*) to integrate gene expression data across species using one-to-one orthologues. After subsetting to keep only cells from ontologically equivalent brain regions, each dataset was normalized independently using Seurat's *SCTransform* v2 function, which corrects for differences in sequencing depth. Each dataset was regressed by percent of mitochondrial genes and animal of origin, except for the Yao et al. dataset in which the only variable to regress was external_donor_name_label. The datasets were then merged together in a list class object, from which HVGs for integration were computed by performing DE analysis on individual objects using the *FindAllMarkers* function in seurat with default parameters. From the DE analysis, we kept all genes of each cluster that had at least a positive 0.2 difference in the percent of expressed cells. We then combined dataset-specific gene lists into a single list containing 6,193 genes, and kept only genes that were present across all the datasets, resulting in a final list of 3,548 genes used for integration. Pairs of mutual nearest neighbors (anchors) were identified using the *FindIntegrationAnchors* function with the arguments reduction="CCA" (Canonical Correlation Analysis) and normalization method= SCT. Integration was carried out with the seurat function *IntegrateData*, normalizing with SCT, using the anchor sets obtained from *FindIntegrationAnchors* and 120 PCs. Dimensionality reduction was performed by performing PCA with 200 PCs. Unsupervised clustering was performed with the default method using the SLM algorithm and clustering res= 0.5. The resulting integrated object consisted of 53,823 cells from the four species, visualized in a low dimensional space using the UMAP algorithm. New clusters that result from the unsupervised clustering of the integrated datasets, also referred to as integrated clusters, were used to assess and quantify the co-clustering between the datasets.

*Differential gene expression analysis and gene module scores in the cross-species analysis*
To independently reconstruct the gene signatures shared between glutamatergic neurons that co-clustered in the cross-species integration, we reasoned to first compute differentially expressed genes (DEGs) between neurons in individual datasets, and then intersect these genes to find shared DEGs across datasets. Specifically, we ran in seurat (v4.3) the *FindMarkers* function between piriform Pyr versus SL cells, and between salamander neurons that co-clustered with them, namely deep layer neurons of DP/VP versus neurons of LP, respectively. Different subtypes of piriform Pyr and SL cells, as well as different subtypes of salamander DP, VP and LP cells were merged into a single category in the DE analysis. Categories were Pyr, SL, DP, LP, VP. The test.use for *FindMarkers* function was used with default settings (Wilcox). To get the top DEGs, we calculated the difference between percentage 1 and percentage 2 for all comparisons, we then subsetted the DEG list to keep only genes expressed in at least 20% of the cells from each category (Pyr, SL, DP, LP, VP). We next intersected the DEGs between 1) LP vs dDP, 2) LP vs dVP, and 3) SL vs Pyr, and kept only the genes that were present across all three DGE analyses. We repeated this approach for the reverse combinations (for example Pyr vs SL). The final lists consisted of 36 shared genes between SL and LP (out of which 3 were TFs), making a SL-like gene module, and 61 shared genes between Pyr and DP/VP (out of which 3 were TFs), making a Pyr-like gene module. To visualize these results, we obtained the average expression matrix from the SCT assay with the *AverageExpression* function in datasets that were subset to the cell types of interest. We

finally used the *AddModuleScore* function to examine the SL-like and Pyr-like modules across cortical areas in mouse and across species. A positive score indicates that the set of genes in the module are expressed in a particular cluster more highly compared to the average expression across all clusters.

**Histology**

*Histological validation of the microdissected cortical areas*

The remaining tissue from micro-dissections for sequencing experiments was fixed in 4% PFA at 4°C overnight. Coronal sections (200 µm thick) were prepared using a vibrating-blade Leica VT100S Vibratome and incubated in PBS, 0.1% Triton X-100 and Neurotrace counterstain diluted 1:1000 (ThermoFisher, N21483) at 4°C overnight.

*Immunohistochemistry*

3 adult Vglut1-Cre/INTACT-GFP transgenic mice, 2 adult wild-type mice, and 4 adult wild-derived mice were deeply anesthetized with 2.5% of 250 mg/kg Avertin and transcardially perfused with 10 ml of ice-cold PBS followed by 10 ml of 4% PFA. Brains were dissected and post-fixed in 4% PFA at 4°C for 5 hr. Coronal and horizontal sections (100 µm thick) were prepared using a Vibratome, permeabilized in PBS and 0.1% Triton X-100 for 1 h, blocked in PBS, 0.1% Triton X-100 and 2% heat-inactivated horse serum at 4°C for 4 h, and incubated with primary antibodies at 4°C overnight (see the list of primary antibodies used below). Note that for horizontal sections we specifically combined, in the same section, two piriform-specific markers with opposite gradients (DCX/posterior with RORB/anterior, and RELN/posterior with CARTPT/anterior) to ensure that the tissue within the section was piriform cortex throughout the anterior-posterior axis. Sections were rinsed in PBS and 0.1% Triton X-100 three times for 15 min, blocked in PBS, 0.1% Triton X-100 and 2% heat-inactivated horse serum at 4°C for 4h, and incubated with appropriate secondary antibodies (1/1000, Donkey IgG (H+L)) conjugated to 405, 488, Cy3 and Cy5 (Jackson Labs) together with DAPI (note that DAPI was not used in all experiments) at 4°C overnight. All histology samples were rinsed with PBS and mounted on SuperFrost Premium microscope slides (Fisher, 12-544-7) in Fluorescent Vectashield Mounting Medium (Vector, H-1900), and imaged at 10X and 20X using a Nikon A1R-HD confocal microscope.

Primary antibodies used were: RELN (mouse, 1/500, MAB5364 Millipore Sigma), RORB (mouse, 200, PP-N7-927-00 Perseus), CARTPT (rabbit, 1/2000, H-0003-62 Phoenix pharmaceutical), KCNG3 (rabbit, 1/1000, TA351316 Origene), EBF1 (rabbit, 1/800, AB10523 Millipore Sigma), EBF2 (sheep, 1/100, AF7006-SP R&D systems), PAPPA2 (goat, 1/500, AF1668-SP R&D systems), MEIS1 (rabbit, 1/1000, ab19867 Abcam), ST8SIA2 (rabbit, 1/200, 19736-1-AP Proteintech), DCX (guinea pig, 1/500, AB2253 Millipore Sigma), SATB1 (rabbit, 1/1500, ab109122 Abcam), CUX1 (rabbit, 1/500, discontinuous-Santacruz), GABA (rabbit, 1/500, A2052 Sigma), TBR1 (rabbit, 1/500, ab31940 Abcam), SOX11 (rabbit, 1/1000, ab134107 Abcam), BARHL1 (rabbit, 1/500, HPA 004809 Sigma).

**Supplementary text: abbreviations**

A: aPir biological replicates

ACA: anterior cingulate cortex

aDVR: anterior dorsal ventricular ridge

AI: agranular insular cortex

AON: anterior olfactory nucleus

aPir: anterior piriform cortex

Astro: astrocytes

AUD: auditory cortex

CA 1/2/3: hippocampal fields

CT: cortico-thalamic projection neurons

DCtx: dorsal cortex

DG: dentate gyrus

DP: dorsal pallium

ENT: entorhinal cortex

FC: fasciola cinereal area

GU: gustatory cortex

IG: induseum griseum area

ILA: infralimbic cortex

IN_CGE: inhibitory neurons from caudal ganglionic eminence

IN_MGE: inhibitory neurons from medial ganglionic eminence

INs: inhibitory neurons

IT: intratelencephalic projection neurons

L: layer

LCtx: lateral cortex

LEC: lateral entorhinal cortex

LOT: lateral olfactory tract

LP: lateral pallium

MCtx: medial cortex

MEC: medial entorhinal cortex

Micro: microglia

Mop: primary motor cortex

MOs_FRP: secondary motor cortex and frontal pole

MP: medial pallium

N: SSp biological replicates

NCx: neocortex

NP: near projecting projection neurons

Oligo: oligodendrocytes

OPC_diff: differentiating oligodendrocytes

OPC: oligodendrocyte precursors cells

ORB: orbital cortex

P: pPir biological replicates

PERI: perirhinal cortex

PL: prelimbic area

POE: post-olfactory eminence

pPir: posterior piriform cortex

PPP: para/post/pre subiculum

ProS: prosubiculum

PT: pyramidal tract projection neurons

PTLp: posterior parietal association cortex

Pyr: pyramidal cells

RHP: retrohippocampal area

RSP: retrosplenial cortex

SL: semilunar cells

SSp: primary somatosensory cortex

SSs: secondary somatosensory cortex

SUB: subiculum

T: AI biological replicates

Tea: temporal association cortex

TPE: Temporal association areas, Perirhinal area, Ectorhinal area

VIS: visual cortex

VISC: visceral area

VISl: lateral visual cortex

VISp: primary visual cortex

VLMC: vascular leptomeningeal cells

VP: ventral pallium

W: wild mice

**Supplementary figures S1 to S24**

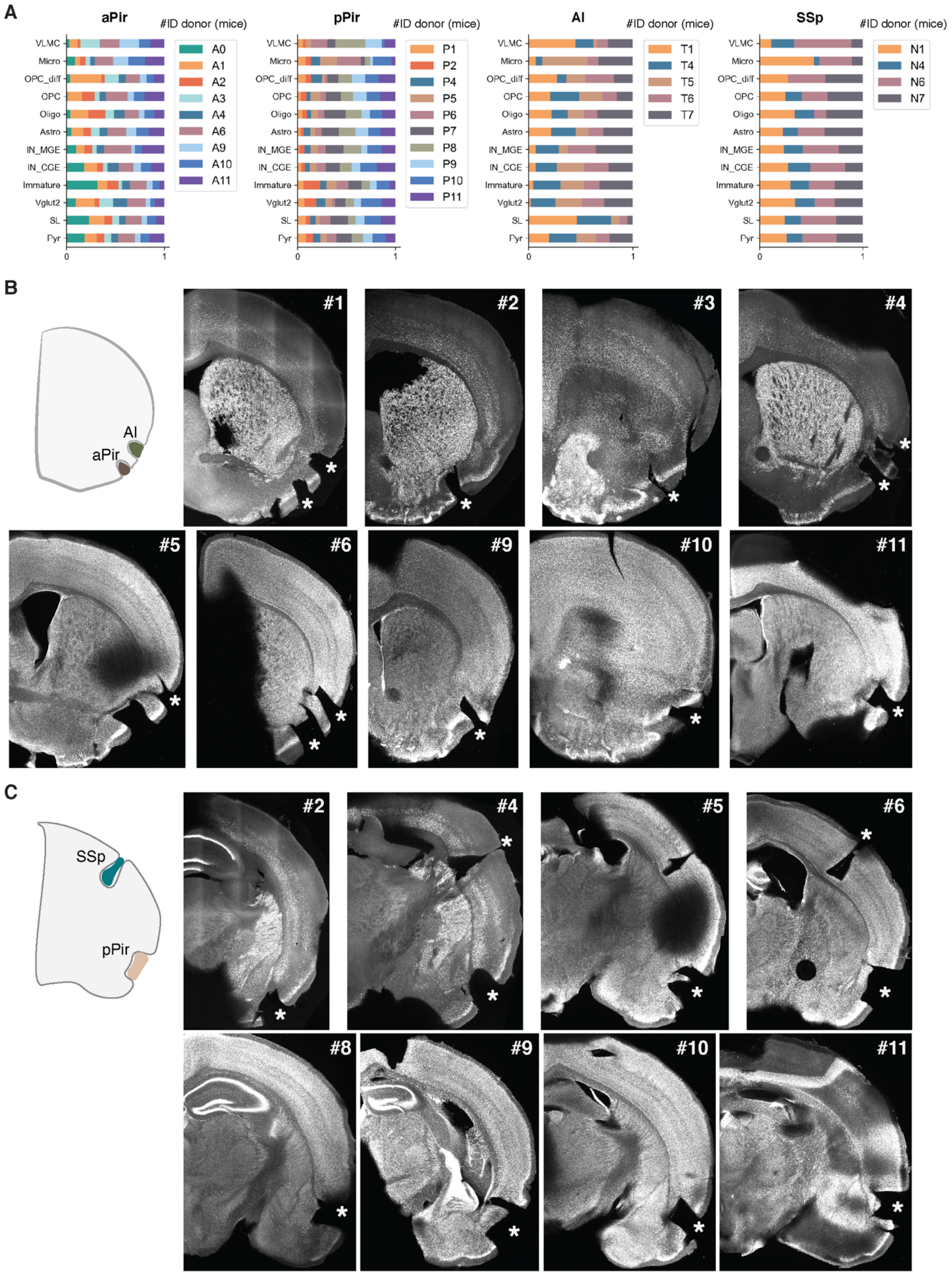
A
aPir
#ID donor (mice)
A0
A1
A2
A3
A4
A6
A9
A10
A11
pPir
P1
P2
P4
P5
P6
P7
P8
P9
P10
P11
AI
T1
T4
T5
T6
T7
SSp
N1
N4
N6
N7
VLMC
Micro
OPC_diff
OPC
Oligo
Astro
IN_MGE
IN_CGE
Immature
Vglut2
SL
Pyr
0
1
B
AI
aPir
#1
#2
#3
#4
#5
#6
#9
#10
#11
C
SSp
pPir
#2
#4
#5
#6
#8
#9
#10
#11

**Fig. S1. Relative abundance of cell types and histological assessment across biological replicates of lab mice.**

**(A)** Stacked bar plots representing the relative abundance of main cell types across biological replicates (donors) integrated from single-nucleus multiome sequencing (sn-multiome seq) experiments. From left to right: anterior piriform cortex (aPir) replicates, indicated by A; posterior piriform cortex (pPir) replicates, indicated by P; agranular insular cortex (AI) replicates, indicated by T; primary somatosensory cortex (SSp) replicates, indicated by N. Numbers correspond to the ID of mice. VLMC: vascular leptomeningeal cells; Micro: microglia; OPC_diff: differentiating oligodendrocytes; OPC: oligodendrocyte precursors; Oligo: oligodendrocytes; Astro: astrocytes; IN_MGE and IN_CGE: inhibitory neurons from medial and caudal ganglionic eminence, respectively; SL: semilunar cells; Pyr: pyramidal cells. **(B)** Post-hoc histological assessment of aPir and AI dissections from anterior coronal sections of adult lab mice ordered by ID mouse number. Asterisks indicate the micro-dissected area. Neurotrace counterstain in gray. **(C)** Same as in **(B)** but for pPir and SSp.

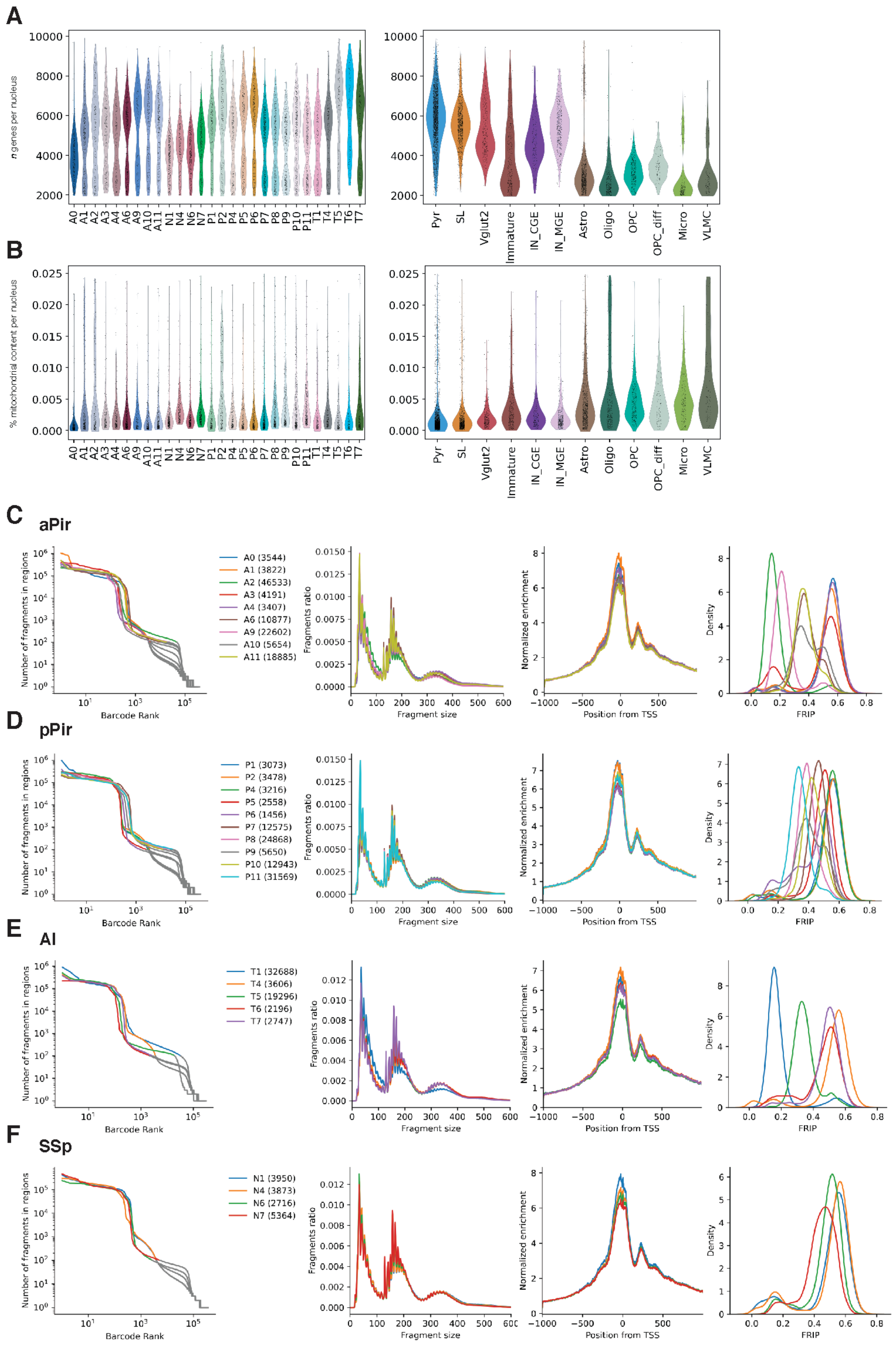
A
B
C
aPir
D
pPir
E
AI
F
SSp

**Fig. S2. Quality control of single-nucleus multiome (RNA and ATAC) sequencing data of lab mice.**

**(A)** Transcriptome quality control. Left: number of genes per nucleus quantified for each biological replicate. A indicates aPir replicates, P indicates pPir, T indicates AI, N indicates SSp. Numbers correspond to the ID of mice. Right: number of genes per nucleus identified for each main cell type. **(B)** Same as in **(A)** but for the percentage of mitochondrial content per nucleus. **(C)** Epigenome quality control for aPir biological replicates. From left to right: barcode rank plot, fragment size distribution, Transcription Start Site (TSS) enrichment and Fraction of Reads In Peaks (FRIP). **(D)** Same as in **(C)** but for pPir biological replicates. **(E)** Same as in **(C)** but for AI biological replicates. **(F)** Same as in **(C)** but for SSp biological replicates.

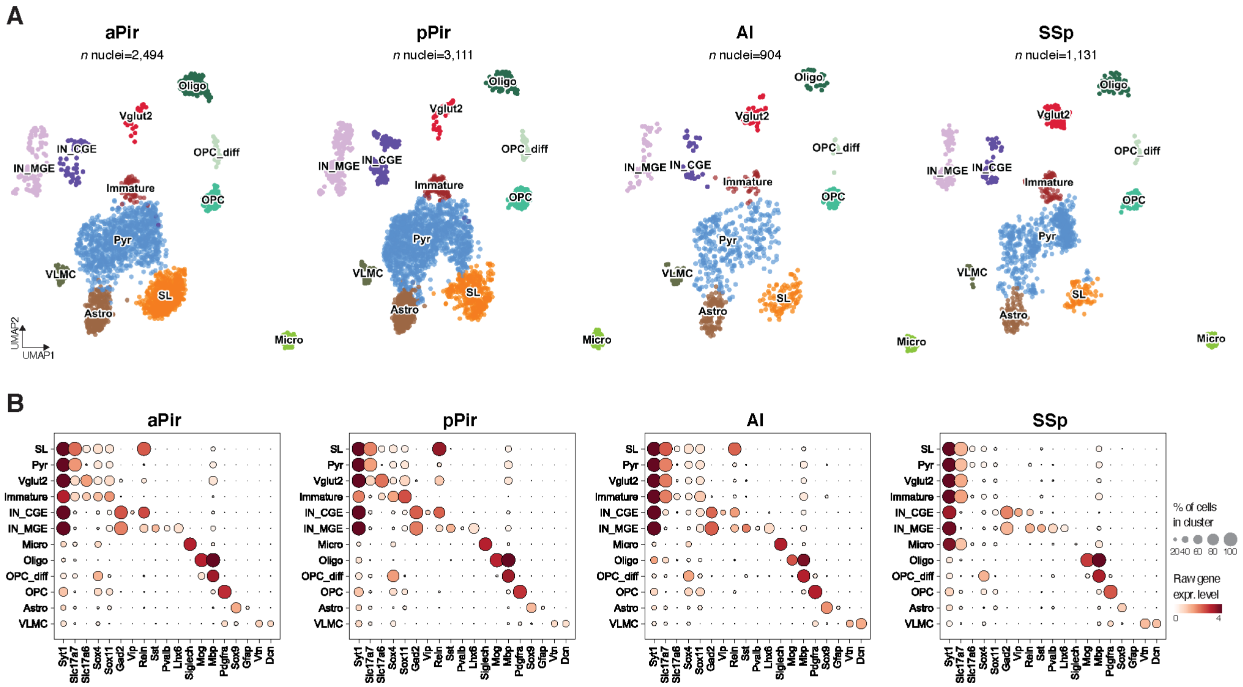


**Fig. S3. Transcriptomically-defined cell types across cortical areas of lab mice.**

**(A)** From left to right: UMAP representations of aPir, pPir, AI, and SSp datasets color-coded by main cell types. **(B)** Dot plots showing gene expression levels of representative markers for each cell type across the four cortical areas, from left to right: aPir, pPir, AI, and SSp.

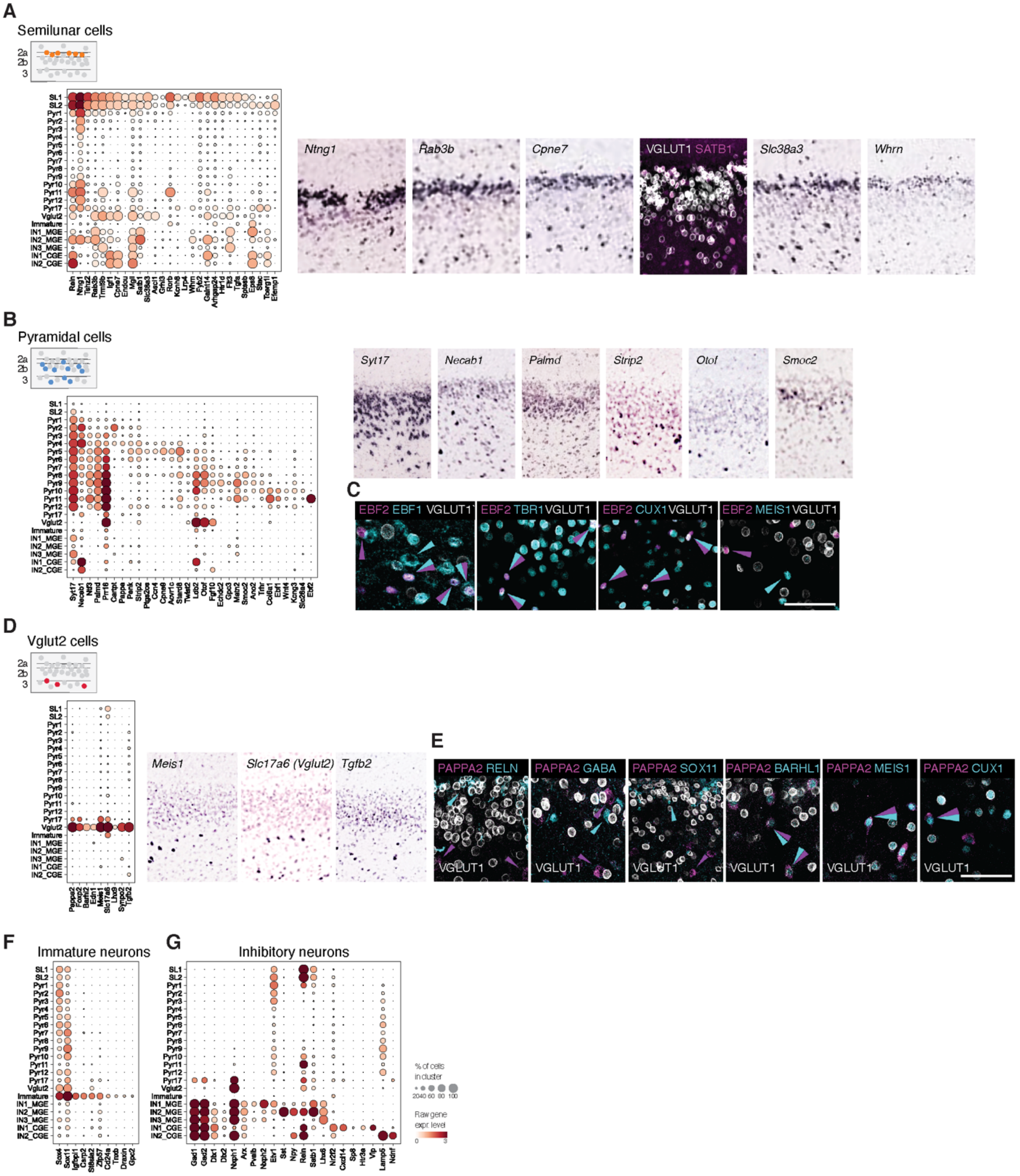

A
Semilunar cells
Ntng1
Rab3b
Cpne7
VGLUT1 SATB1
Slc38a3
Whrn
B
Pyramidal cells
Syt17
Necab1
Palmd
Strip2
Otof
Smoc2
C
EBF2 EBF1 VGLUT1
EBF2 TBR1 VGLUT1
EBF2 CUX1 VGLUT1
EBF2 MEIS1 VGLUT1
D
Vglut2 cells
Meis1
Slc17a6 (Vglut2)
Tgfb2
E
PAPPA2 RELN
PAPPA2 GABA
PAPPA2 SOX11
PAPPA2 BARHL1
PAPPA2 MEIS1
PAPPA2 CUX1
VGLUT1
F
Immature neurons
G
Inhibitory neurons
% of cells in cluster
Raw gene expr level

**Fig. S4. Piriform cortex-specific markers and histological validation.**

**(A)** Left: dot plot showing gene expression levels of generic or subtype-specific markers for semilunar cells in the combined aPir and pPir datasets. Right: *in situ* hybridization images from the Allen Brain Atlas for some of the markers shown in the dotplot at the left, with the exception of the marker SATB1, which was validated using immunohistochemistry in *Vglut1*-CRE/INTACT-GFP transgenic mice. **(B)** Same as **(A),** but for generic or subtype-specific markers for pyramidal cells in the combined aPir and pPir datasets. **(C)** High magnifications in Pir layer 3 of immunohistochemical experiments using *Vglut1*-CRE/INTACT-GFP transgenic mice showing combinatorial expression of the specific marker for Pyr11, EBF2 (in magenta), with specific or generic markers for glutamatergic neurons (in cyan): EBF1 (Pyr10-11-specific); TBR1 and CUX1 (pan-excitatory/pyramidal neuron marker); MEIS1 (Vglut2-specific). Scale bar, 100 µm. **(D)** Same as **(A),** but for generic or subtype-specific markers for Vglut2 cells in the combined aPir and pPir datasets. **(E)** High magnifications in Pir layer 3 of immunohistochemical experiments using *Vglut1*-CRE/INTACT-GFP transgenic mice showing combinatorial expression of the specific marker for Vglut2 cells, PAPPA2 (in magenta), with generic markers for semilunar cells (RELN), INs (GABA), immature neurons (SOX11), and pyramidal cells (CUX1), or with markers highly specific to Vglut2 cells (MEIS1), or expressed in Pir layer 3 (BARHL1) (in cyan), to understand the identity of the uncharacterized Vglut2-expressing neuronal population. Scale bar, 100 µm. **(F)** Same as **(A),** but for generic or subtype-specific markers for immature neurons in the combined aPir and pPir datasets. **(G)** Same as **(A),** but for generic or subtype-specific markers for INs in the combined aPir and pPir datasets.

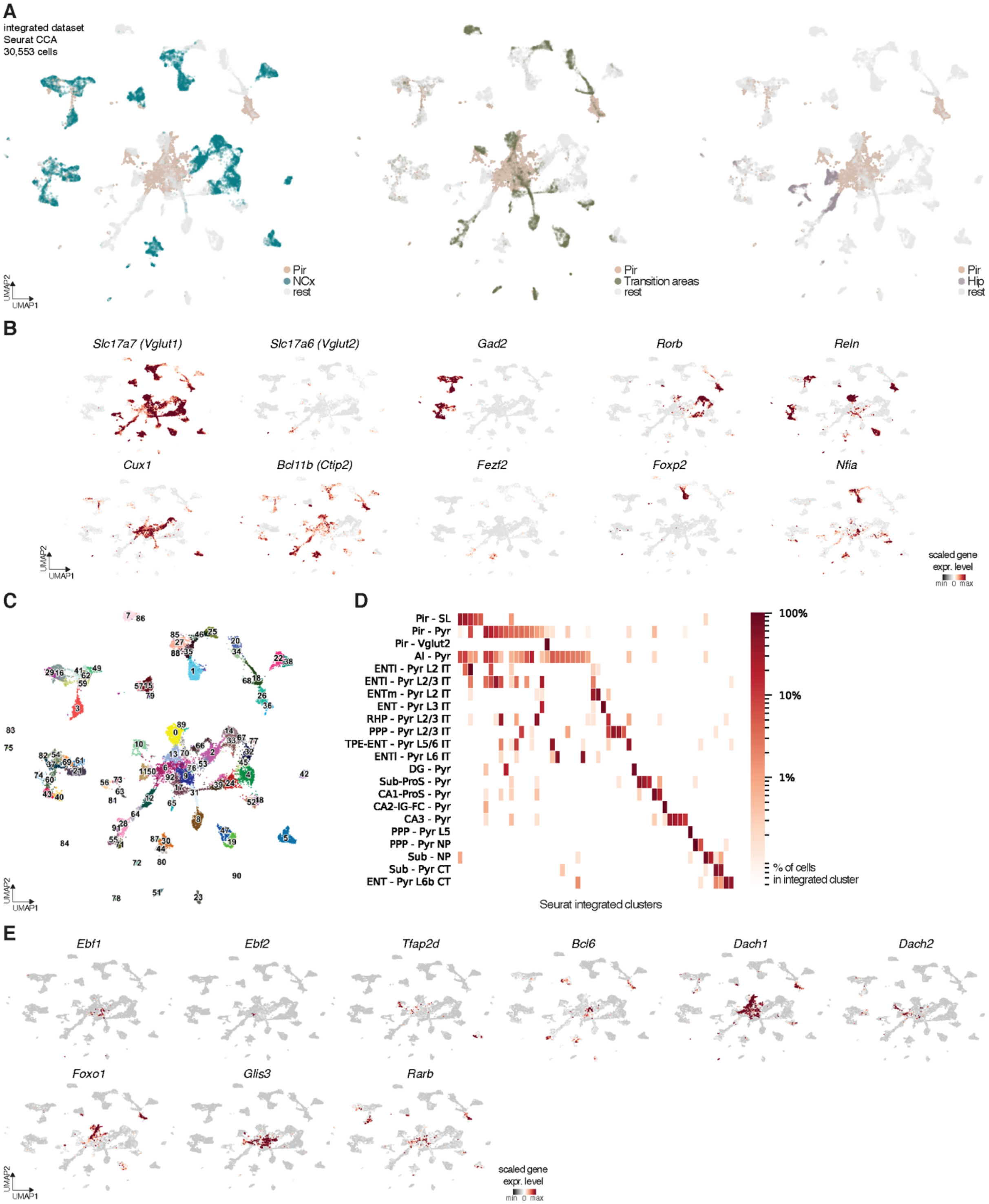
A
integrated dataset
Seurat CCA
30,553 cells
Pir
NCx
rest
Pir
Transition areas
rest
Pir
Hip
rest
UMAP2
UMAP1
B
Slc17a7 (Vglut1)
Slc17a6 (Vglut2)
Gad2
Rorb
Reln
Cux1
Bcl11b (Ctip2)
Fezf2
Foxp2
Nfia
scaled gene expr. level
min 0 max
C
D
Pir - SL
Pir - Pyr
Pir - Vglut2
AI - Pyr
ENTl - Pyr L2 IT
ENTl - Pyr L2/3 IT
ENTm - Pyr L2 IT
ENT - Pyr L3 IT
RHP - Pyr L2/3 IT
PPP - Pyr L2/3 IT
TPE-ENT - Pyr L5/6 IT
ENTl - Pyr L6 IT
DG - Pyr
Sub-ProS - Pyr
CA1-ProS - Pyr
CA2-IG-FC - Pyr
CA3 - Pyr
PPP - Pyr L5
PPP - Pyr NP
Sub - NP
Sub - Pyr CT
ENT - Pyr L6b CT
Seurat integrated clusters
100%
10%
1%
% of cells in integrated cluster
E
Ebf1
Ebf2
Tfap2d
Bcl6
Dach1
Dach2
Foxo1
Glis3
Rarb

**Fig. S5. Integration of neurons from piriform cortex, transition areas, hippocampal formation and neocortical areas.**

**(A)** UMAP representations of the integration between neurons from *this study,* which includes Pir, AI and SSp, and from (*17*), which includes transition areas, such as entorhinal cortex and AI, hippocampal formation, and neocortical areas, such as SSp, motor and visual cortex (n=30,553). From left to right: Pir and NCx neurons are visualized; Pir and transition areas neurons are visualized; Pir and hippocampal formation (Hip) neurons are visualized. Datasets were integrated using Canonical Correlation Analysis (CCA). **(B)** UMAP representations of gene expression for the integrated datasets shown in **(A)** of generic markers for glutamatergic neurons (*Slc17a7* and *Slc17a6*), inhibitory neurons (INs) (*Gad2*), and other established markers for neocortical projection neurons: *Rorb* (layer 4), *Cux1* (layer 2/3), *Ctip2* (layer 5), *Fezf2* (layer 5), *Foxp2* (layer 6), *Nfia* (layer 6). **(C)** UMAP representation of the integrated datasets shown in **(A)** color-coded by the new seurat integrated clusters. **(D)** Heatmap showing quantification of co-clustering between piriform, hippocampal formation and transition areas neurons from the integration shown in **(A)**. Neurons grouped into main types for each cortical area. Rectangles indicate co-clustering of neurons in the new seurat integrated clusters. Color of the rectangle represents the percentage of cells in the integrated cluster. L: layer; IT: intratelencephalic; NP: near projecting; CT: cortico-thalamic; Sub: subiculum; ProS: prosubiculum; PPP: para/post/pre subiculum; RHP: retrohippocampal region; DG: dentate gyrus; CA 1/2/3: hippocampal fields; IG: induseum griseum; FC: fasciola cinereal; AI: agranular insular; ENT: entorhinal (medial and lateral); TPE: Temporal association areas, Perirhinal area, Ectorhinal area. **(E)** UMAP representations of gene expression of transcription factors (TFs) highly enriched in piriform cortex compared to other cortical areas.

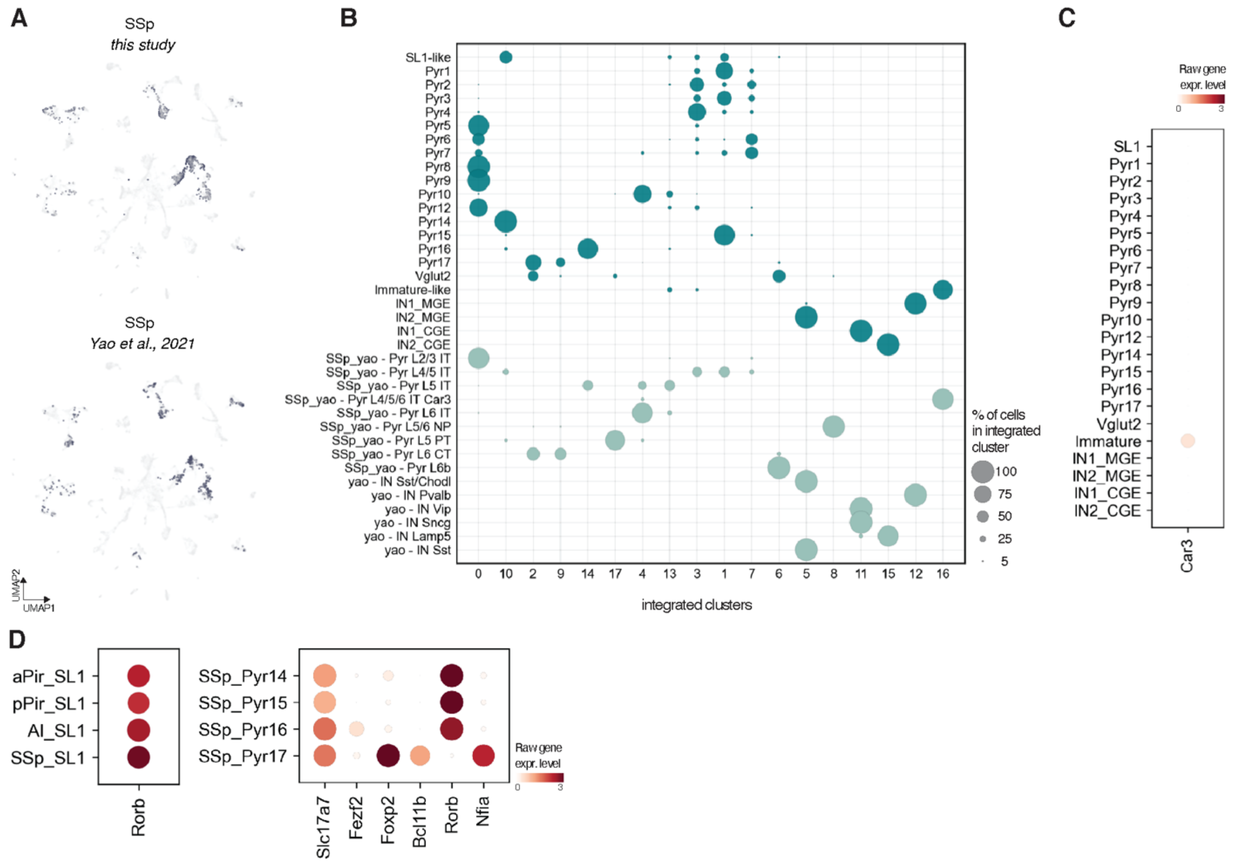


**Fig. S6. Comparison of neurons from SSp datasets from *this study* and a mouse single-cell reference atlas**

**(A)** UMAP representations of integrated neurons from *this study* and from the mouse single-cell reference atlas (bottom) (*17*), and visualization of only SSp neurons from the two studies (top: from *this study,* bottom: from single-cell reference atlas). **(B)** Dotplot showing quantification of co-clustering between SSp neurons of *this study* and of the single-cell reference atlas. Only SSp datasets were included in the integration to transfer projection neuron type nomenclature to *this study*. SSp_yao indicates neurons from (*17*), while everything else corresponds to SSp clusters of *this study.* Dots indicate co-clustering of neurons in the new seurat integrated clusters. Size of the dots represents the percentage of cells in the integrated cluster. **(C)** Dot plots showing gene expression levels of *Car3* in the SSp dataset. *Car3* is expressed in SSp neurons co-clustering with piriform neurons in the immature neuron supertype. The expression of *Car3* is consistent with the co-clustering of SSp immature-like neurons of our dataset with L4/5/6 intratelencephalic *Car3* neurons of the single-cell reference atlas. **(D)** Dot plots showing gene expression levels of the TF *Rorb*, which is highly enriched in the SL1 supertype comprising all four cortical areas (left), and of established TFs present in the SSp-specific supertypes (right). Pyr 14-15-16 of layers 4 and 5 are characterized by *Rorb* and *Fezf2*, Pyr17 is characterized by *Foxp2*, *Bcl11b* (*Ctip2*) and *Nfia*, corresponding to CT neurons of layers 6.

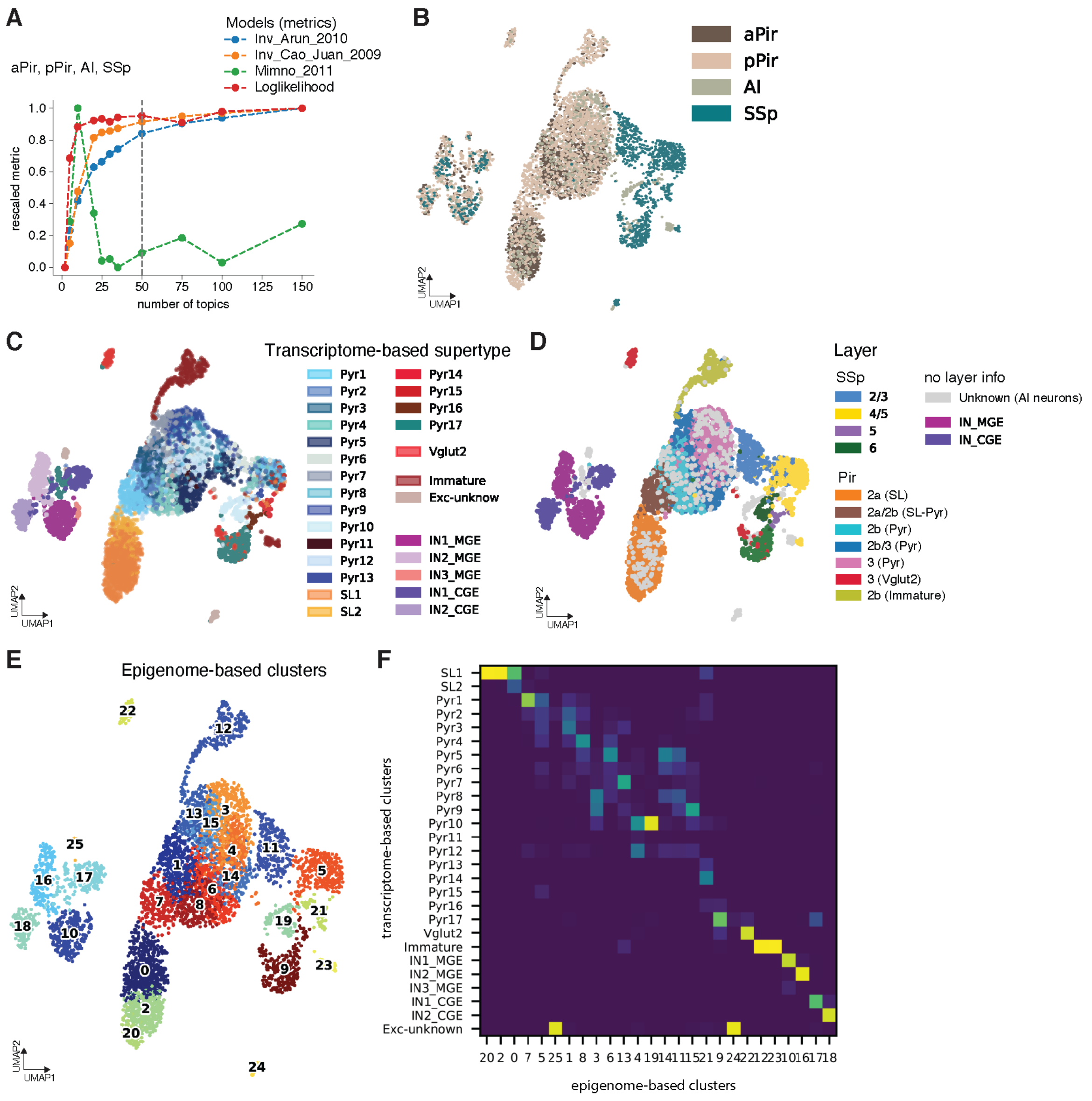


**Fig. S7. Epigenetic divergence of transcriptome-based supertypes.**

**(A)** An optimal number of 50 topics (gray vertical line) are used in the PycisTopic model selection for imputation, dimensionality reduction, and clustering the corresponding ATAC seq data of combined neurons from the four cortical areas (aPir, pPir, AI, SSp). **(B)** UMAP representation of the corresponding ATAC seq data color-coded by aPir, pPir, AI, and SSp datasets. Neurons are integrated using Harmony. **(C)** UMAP representation of the ATAC seq data color-coded by the corresponding transcriptome-based supertype. Note that areas are mixed for INs, while piriform separates from SSp in the glutamatergic neurons, with AI overlapping with both (see **(B)**). **(D)** UMAP representation of the ATAC seq data color-coded by cortical layers. Layers are assigned based on gene expression and histology for piriform, and based on the mouse single-cell reference atlas for SSp. For AI, cortical layer information was not available (unknown). **(E)** UMAP representation of the ATAC seq data color-coded by ATAC clusters (leiden). **(E)** Mapping of transcriptome-based clusters (RNA, supertypes) to epigenome-based clusters (ATAC, leiden). Note that for glutamatergic neurons multiple area-specific epigenome-based clusters correspond to a single supertype. Adjusted Rand Indices (ARIs) are computed, which quantify the cluster overlap. ARIs are: for all neurons= 0.43; for INs= 0.88; for glutamatergic neurons= 0.37.

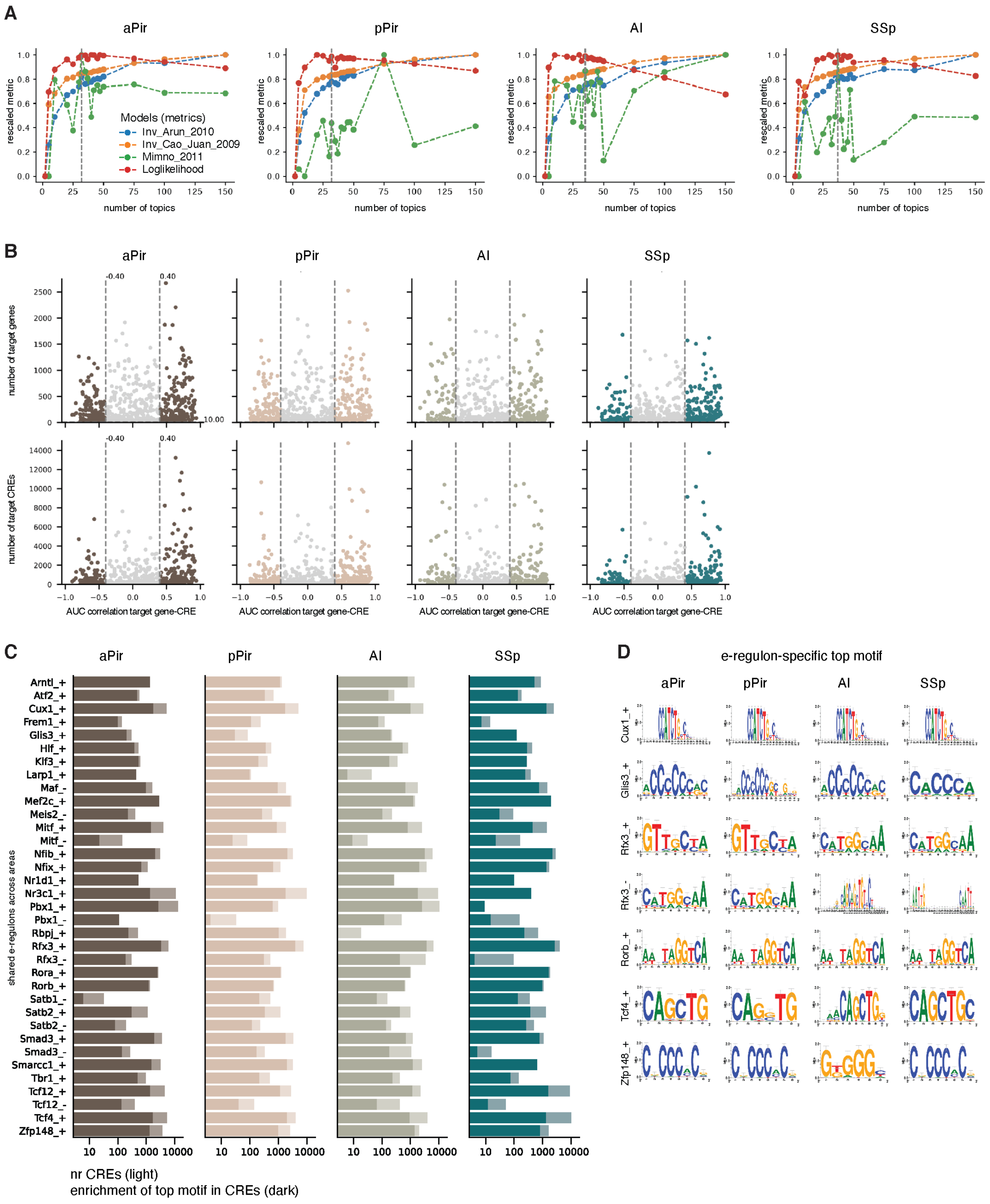
A
aPir
pPir
AI
SSp
rescaled metric
number of topics
Models (metrics)
Inv_Arun_2010
Inv_Cao_Juan_2009
Mimno_2011
Loglikelihood
B
number of target genes
number of target CREs
AUC correlation target gene-CRE
-0.40
0.40
C
shared e-regulons across areas
Arntl_+
Atf2_+
Cux1_+
Frem1_+
Glis3_+
Hlf_+
Klf3_+
Larp1_+
Maf_-
Mef2c_+
Meis2_-
Mitf_+
Mitf_-
Nfib_+
Nfix_+
Nr1d1_+
Nr3c1_+
Pbx1_+
Pbx1_-
Rbpj_+
Rfx3_+
Rfx3_-
Rora_+
Rorb_+
Satb1_-
Satb2_+
Satb2_-
Smad3_+
Smad3_-
Smarcc1_+
Tbr1_+
Tcf12_+
Tcf12_-
Tcf4_+
Zfp148_+
nr CREs (light)
enrichment of top motif in CREs (dark)
D
e-regulon-specific top motif
Cux1_+
Glis3_+
Rfx3_+
Rfx3_-
Rorb_+
Tcf4_+
Zfp148_+

**Fig. S8. Enhancer-driven gene regulatory networks (e-GRNs) of aPir, pPir, AI, and SSp.**

**(A)** An optimal number of 32, 32, 35, and 37 topics for aPir, pPir, AI, and SSp respectively (gray vertical lines), are used in the PycisTopic model selection for imputation, dimensionality reduction, and clustering the corresponding ATAC seq data of neurons from each area. **(B)** High-quality e-regulons are selected for downstream analysis based on the correlation between AUC scores for target genes and target CREs. A correlation cut-off of 0.4 and a minimum number of target genes of 10 are used. CRE: *cis* regulatory element; AUC: Area Under the Curve. **(C)** Bar charts showing number of CREs and enrichment score of the top TF binding motif in these CREs for the 35 e-regulons shared across areas, which are specific for glutamatergic neurons. The 35 e-regulons were found to use multiple binding site motifs, with on average the top motif (with the highest normalized enrichment score) being found in more than 54% of the CREs associated with the respective e-regulon. **(D)** From left to right, it is shown the top area-specific TF binding site motif for a few example e-regulons shared across areas.

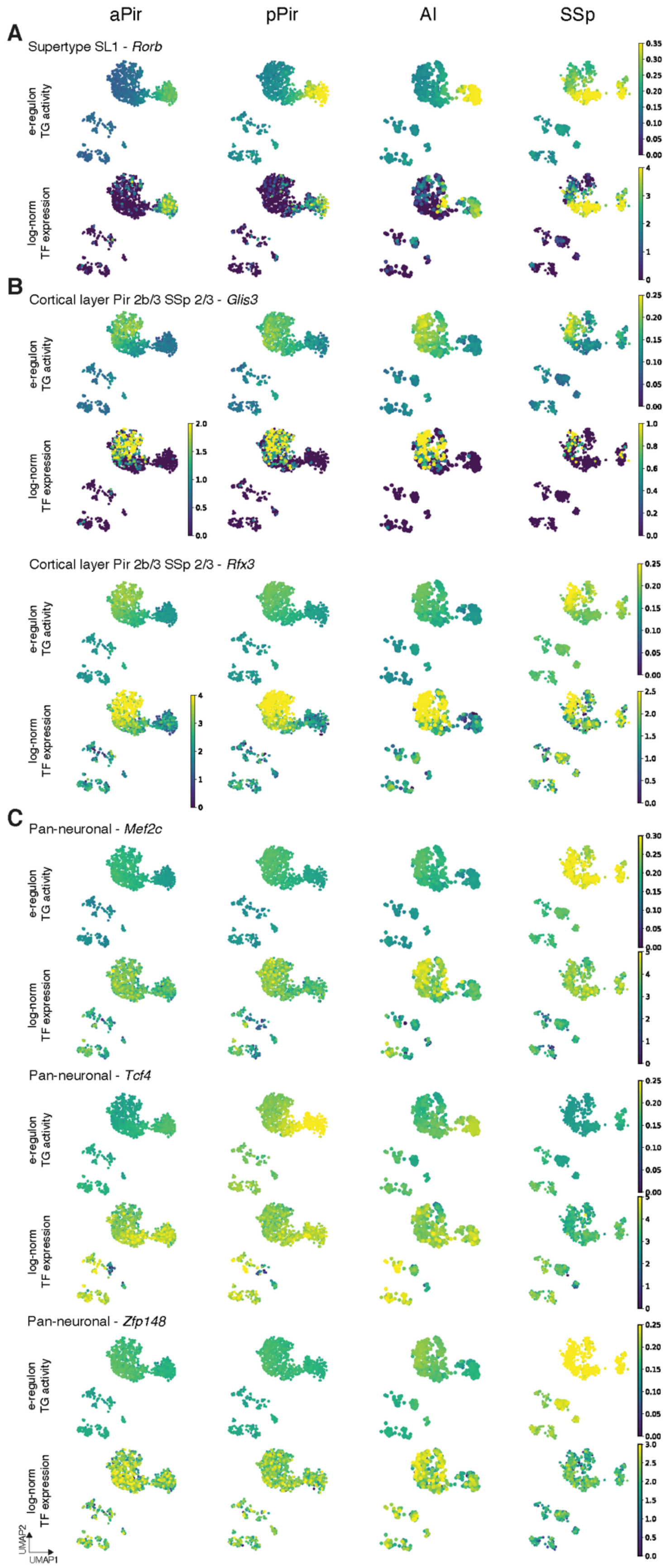
aPir
pPir
AI
SSp
A
Supertype SL1 - *Rorb*
e-regulon TG activity
log-norm TF expression
B
Cortical layer Pir 2b/3 SSp 2/3 - *Glis3*
Cortical layer Pir 2b/3 SSp 2/3 - *Rfx3*
C
Pan-neuronal - *Mef2c*
Pan-neuronal - *Tcf4*
Pan-neuronal - *Zfp148*
UMAP2
UMAP1

**Fig. S9. Examples of e-regulons shared across cortical areas that are specific for a particular supertype, layer, or active in all neurons.**

**(A)** UMAP representations of target gene (TG) activity expressed as an AUC score and log-normalized TF expression of the TF that is part of a given e-regulon. Here is shown as an example the shared e-regulons *Rorb* across aPir, pPir, AI, and SSp, which exhibits specificity for the supertype SL1. **(B)** UMAP representations of TG activity and log-normalized TF expression of the e-regulons *Glis3* and *Rfx3*, as examples of shared e-regulons across aPir, pPir, AI, and SSp that exhibit specificity for piriform layers 2b/3 and SSp layer 2/3. **(C)** UMAP representations of TG activity and log-normalized TF expression of the e-regulons *Mef2c, Tcf4*, and *Zfp148*, as examples of shared e-regulons across aPir, pPir, AI, and SSp that are active in all neurons (pan-neuronal).

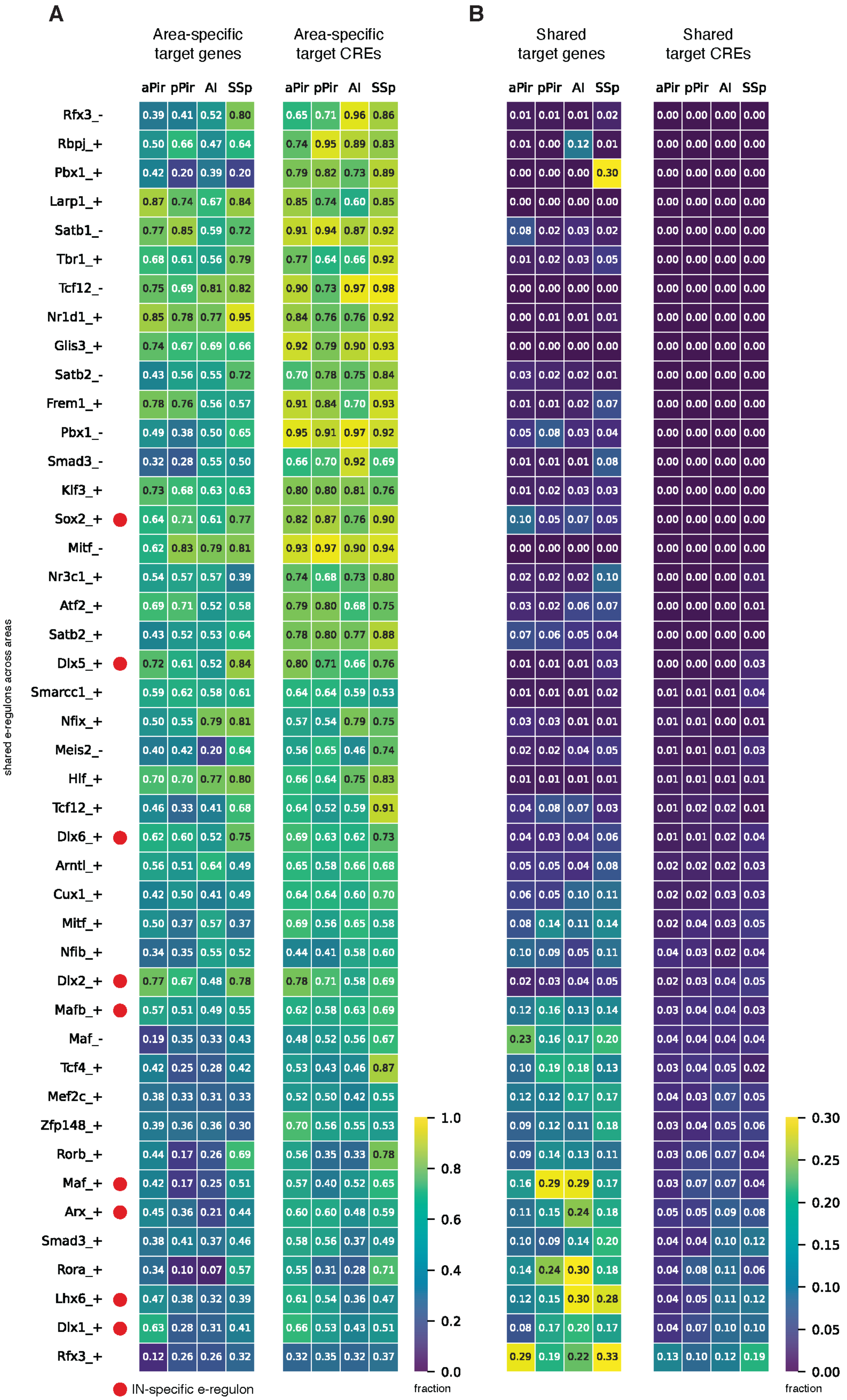
A
Area-specific target genes
Area-specific target CREs
B
Shared target genes
Shared target CREs
aPir pPir AI SSp
shared e-regulons across areas
Rfx3_- 0.39 0.41 0.52 0.80 | 0.65 0.71 0.96 0.86 | 0.01 0.01 0.01 0.02 | 0.00 0.00 0.00 0.00
Rbpj_+ 0.50 0.66 0.47 0.64 | 0.74 0.95 0.89 0.83 | 0.01 0.00 0.12 0.01 | 0.00 0.00 0.00 0.00
Pbx1_+ 0.42 0.20 0.39 0.20 | 0.79 0.82 0.73 0.89 | 0.00 0.00 0.00 0.30 | 0.00 0.00 0.00 0.00
Larp1_+ 0.87 0.74 0.67 0.84 | 0.85 0.74 0.60 0.85 | 0.00 0.00 0.00 0.00 | 0.00 0.00 0.00 0.00
Satb1_- 0.77 0.85 0.59 0.72 | 0.91 0.94 0.87 0.92 | 0.08 0.02 0.03 0.02 | 0.00 0.00 0.00 0.00
Tbr1_+ 0.68 0.61 0.56 0.79 | 0.77 0.64 0.66 0.92 | 0.01 0.02 0.03 0.05 | 0.00 0.00 0.00 0.00
Tcf12_- 0.75 0.69 0.81 0.82 | 0.90 0.73 0.97 0.98 | 0.00 0.00 0.00 0.00 | 0.00 0.00 0.00 0.00
Nr1d1_+ 0.85 0.78 0.77 0.95 | 0.84 0.76 0.76 0.92 | 0.00 0.01 0.01 0.01 | 0.00 0.00 0.00 0.00
Glis3_+ 0.74 0.67 0.69 0.66 | 0.92 0.79 0.90 0.93 | 0.00 0.00 0.00 0.00 | 0.00 0.00 0.00 0.00
Satb2_- 0.43 0.56 0.55 0.72 | 0.70 0.78 0.75 0.84 | 0.03 0.02 0.02 0.01 | 0.00 0.00 0.00 0.00
Frem1_+ 0.78 0.76 0.56 0.57 | 0.91 0.84 0.70 0.93 | 0.01 0.01 0.02 0.07 | 0.00 0.00 0.00 0.00
Pbx1_- 0.49 0.38 0.50 0.65 | 0.95 0.91 0.97 0.92 | 0.05 0.08 0.03 0.04 | 0.00 0.00 0.00 0.00
Smad3_- 0.32 0.28 0.55 0.50 | 0.66 0.70 0.92 0.69 | 0.01 0.01 0.01 0.08 | 0.00 0.00 0.00 0.00
Klf3_+ 0.73 0.68 0.63 0.63 | 0.80 0.80 0.81 0.76 | 0.01 0.02 0.03 0.03 | 0.00 0.00 0.00 0.00
Sox2_+ 0.64 0.71 0.61 0.77 | 0.82 0.87 0.76 0.90 | 0.10 0.05 0.07 0.05 | 0.00 0.00 0.00 0.00
Mitf_- 0.62 0.83 0.79 0.81 | 0.93 0.97 0.90 0.94 | 0.00 0.00 0.00 0.00 | 0.00 0.00 0.00 0.00
Nr3c1_+ 0.54 0.57 0.57 0.39 | 0.74 0.68 0.73 0.80 | 0.02 0.02 0.02 0.10 | 0.00 0.00 0.00 0.01
Atf2_+ 0.69 0.71 0.52 0.58 | 0.79 0.80 0.68 0.75 | 0.03 0.02 0.06 0.07 | 0.00 0.00 0.00 0.01
Satb2_+ 0.43 0.52 0.53 0.64 | 0.78 0.80 0.77 0.88 | 0.07 0.06 0.05 0.04 | 0.00 0.00 0.00 0.00
Dlx5_+ 0.72 0.61 0.52 0.84 | 0.80 0.71 0.66 0.76 | 0.01 0.01 0.01 0.03 | 0.00 0.00 0.00 0.03
Smarcc1_+ 0.59 0.62 0.58 0.61 | 0.64 0.64 0.59 0.53 | 0.01 0.01 0.01 0.02 | 0.01 0.01 0.01 0.04
Nfix_+ 0.50 0.55 0.79 0.81 | 0.57 0.54 0.79 0.75 | 0.03 0.03 0.01 0.01 | 0.01 0.01 0.00 0.01
Meis2_- 0.40 0.42 0.20 0.64 | 0.56 0.65 0.46 0.74 | 0.02 0.02 0.04 0.05 | 0.01 0.01 0.01 0.03
Hlf_+ 0.70 0.70 0.77 0.80 | 0.66 0.64 0.75 0.83 | 0.01 0.01 0.01 0.01 | 0.01 0.01 0.01 0.01
Tcf12_+ 0.46 0.33 0.41 0.68 | 0.64 0.52 0.59 0.91 | 0.04 0.08 0.07 0.03 | 0.01 0.02 0.02 0.01
Dlx6_+ 0.62 0.60 0.52 0.75 | 0.69 0.63 0.62 0.73 | 0.04 0.03 0.04 0.06 | 0.01 0.01 0.02 0.04
Arntl_+ 0.56 0.51 0.64 0.49 | 0.65 0.58 0.66 0.68 | 0.05 0.05 0.04 0.08 | 0.02 0.02 0.02 0.03
Cux1_+ 0.42 0.50 0.41 0.49 | 0.64 0.64 0.60 0.70 | 0.06 0.05 0.10 0.11 | 0.02 0.02 0.03 0.03
Mitf_+ 0.50 0.37 0.57 0.37 | 0.69 0.56 0.65 0.58 | 0.08 0.14 0.11 0.14 | 0.02 0.04 0.03 0.05
Nfib_+ 0.34 0.35 0.55 0.52 | 0.44 0.41 0.58 0.60 | 0.10 0.09 0.05 0.11 | 0.04 0.03 0.02 0.04
Dlx2_+ 0.77 0.67 0.48 0.78 | 0.78 0.71 0.58 0.69 | 0.02 0.03 0.04 0.05 | 0.02 0.03 0.04 0.05
Mafb_+ 0.57 0.51 0.49 0.55 | 0.62 0.58 0.63 0.69 | 0.12 0.16 0.13 0.14 | 0.03 0.04 0.04 0.03
Maf_- 0.19 0.35 0.33 0.43 | 0.48 0.52 0.56 0.67 | 0.23 0.16 0.17 0.20 | 0.04 0.04 0.04 0.04
Tcf4_+ 0.42 0.25 0.28 0.42 | 0.53 0.43 0.46 0.87 | 0.10 0.19 0.18 0.13 | 0.03 0.04 0.05 0.02
Mef2c_+ 0.38 0.33 0.31 0.33 | 0.52 0.50 0.42 0.55 | 0.12 0.12 0.17 0.17 | 0.04 0.03 0.07 0.05
Zfp148_+ 0.39 0.36 0.36 0.30 | 0.70 0.56 0.55 0.53 | 0.09 0.12 0.11 0.18 | 0.03 0.04 0.05 0.06
Rorb_+ 0.44 0.17 0.26 0.69 | 0.56 0.35 0.33 0.78 | 0.09 0.14 0.13 0.11 | 0.03 0.06 0.07 0.04
Maf_+ 0.42 0.17 0.25 0.51 | 0.57 0.40 0.52 0.65 | 0.16 0.29 0.29 0.17 | 0.03 0.07 0.07 0.04
Arx_+ 0.45 0.36 0.21 0.44 | 0.60 0.60 0.48 0.59 | 0.11 0.15 0.24 0.18 | 0.05 0.05 0.09 0.08
Smad3_+ 0.38 0.41 0.37 0.46 | 0.58 0.56 0.37 0.49 | 0.10 0.09 0.14 0.20 | 0.04 0.04 0.10 0.12
Rora_+ 0.34 0.10 0.07 0.57 | 0.55 0.31 0.28 0.71 | 0.14 0.24 0.30 0.18 | 0.04 0.08 0.11 0.06
Lhx6_+ 0.47 0.38 0.32 0.39 | 0.61 0.54 0.36 0.47 | 0.12 0.15 0.30 0.28 | 0.04 0.05 0.11 0.12
Dlx1_+ 0.63 0.28 0.31 0.41 | 0.66 0.53 0.43 0.51 | 0.08 0.17 0.20 0.17 | 0.04 0.07 0.10 0.10
Rfx3_+ 0.12 0.26 0.26 0.32 | 0.32 0.35 0.32 0.37 | 0.29 0.19 0.22 0.33 | 0.13 0.10 0.12 0.19
1.0 0.8 0.6 0.4 0.2 0.0
fraction
0.30 0.25 0.20 0.15 0.10 0.05 0.00
fraction
IN-specific e-regulon

**Fig. S10. Target genes and target enhancers of the e-regulons shared across cortical areas.** **(A)** Heatmaps showing, for each cortical area and for each shared e-regulons across areas, the fractions of area-specific target genes and target CREs (only enhancers are included) over the total number of target genes and target CREs of each area. For example, the e-regulon *Rbpj* (second row) has 50% of its target genes specific to aPir, meaning these target genes are not identified as predicted to be regulated by *Rbpj* in pPir, AI, and SSp. IN-specific e-regulons are marked with a red dot. **(B)** Heatmaps showing, for each cortical area and for each shared e-regulons across areas, the fractions of target genes and target CREs (only enhancers are included) shared across areas over the total number of target genes and target CREs of each area.

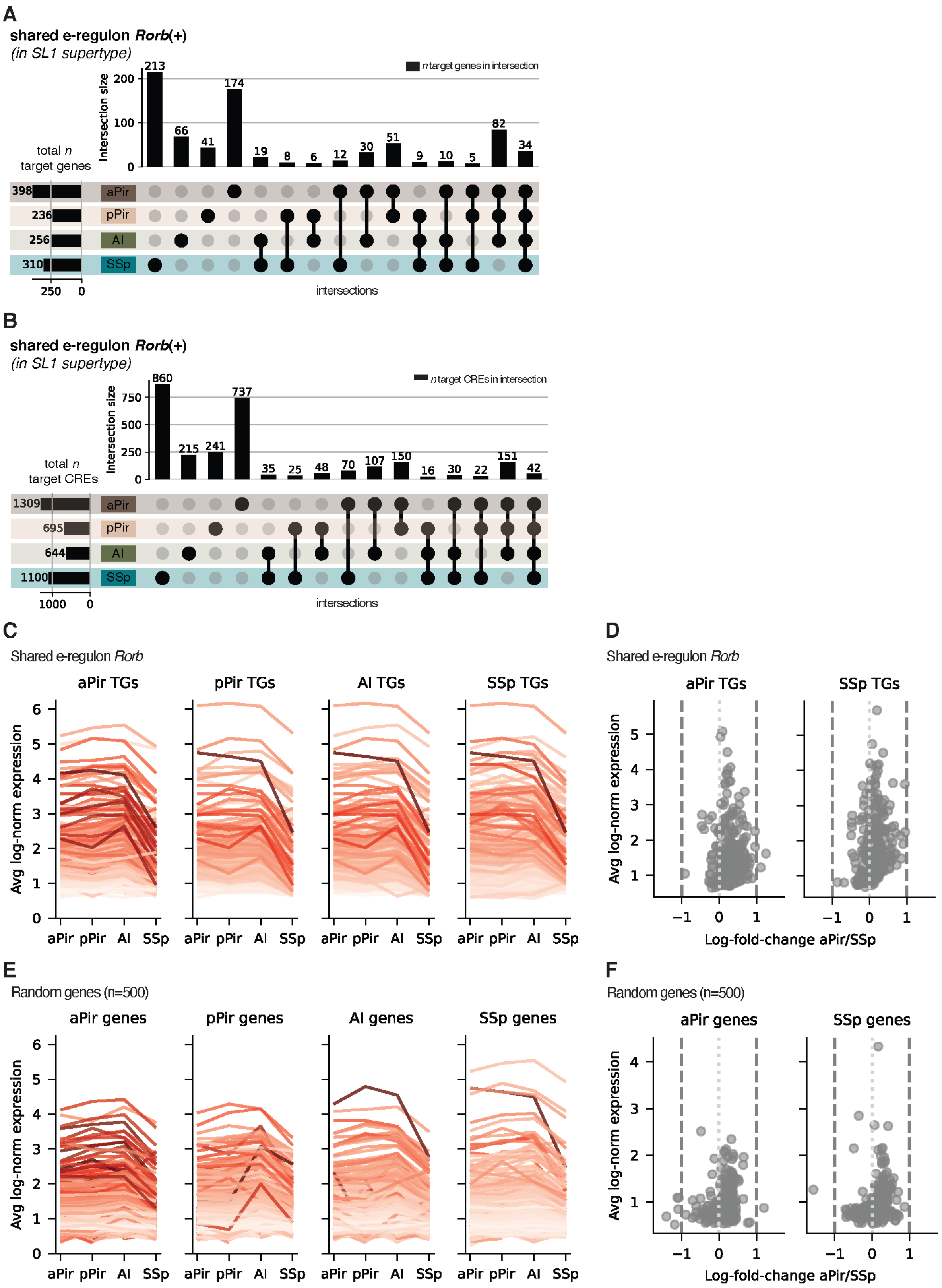
A
shared e-regulon Rorb(+)
(in SL1 supertype)
n target genes in intersection
Intersection size
total n
target genes
aPir
pPir
AI
SSp
intersections
B
shared e-regulon Rorb(+)
(in SL1 supertype)
n target CREs in intersection
total n
target CREs
C
Shared e-regulon Rorb
aPir TGs
pPir TGs
AI TGs
SSp TGs
Avg log-norm expression
D
Shared e-regulon Rorb
Log-fold-change aPir/SSp
E
Random genes (n=500)
aPir genes
pPir genes
AI genes
SSp genes
F
Random genes (n=500)

**Fig. S11. Area-specific target genes and target enhancers of the e-regulon *Rorb*.**

**(A)** Upset plot of the intersection of target genes for the e-regulon *Rorb*(+), which is shared across aPir, pPir, AI and SSp. The vertical bars show the number of target genes in the corresponding intersection of the matrix below. The horizontal bars show the total number of target genes for each cortical area. The 9% overlap between aPir and SSp target genes (mentioned in the main text), is a Jaccard similarity index. It is equivalent to computing in the upset plot the (relative) size of a combination of intersections. **(B)** Upset plot of the intersection of target CREs for the e-regulon *Rorb*(+), which is shared across aPir, pPir, AI and SSp. See panel **(A)** for details. **(C)** From left to right, it is shown the average log-normalization expression of target genes (TGs) of *Rorb* identified in aPir, pPir, AI, and SSp datasets. For each plot, the average expression of TGs is shown also across areas. **(D)** Log-fold-change of gene expression between target genes identified in aPir and SSp datasets for the e-regulon *Rorb*. The log-fold-change remains within a 2-fold change. The average log- normalization expression (y-axis) is only used to spread data points. **(E)** From left to right, it is shown the average log- normalization expression of a random set of genes (n=500) for aPir, pPir, AI, and SSp. For each plot, the average expression of the set of random genes is shown also across areas. Patterns of expression between areas observed in **(C)** are also visible here. **(F)** Log-fold-change of gene expression between set of random genes in aPir and SSp datasets. Log-fold-change remains within a 2-fold change, and indicates that *Rorb* target genes shown in **(D)** behave as any other set of random genes.

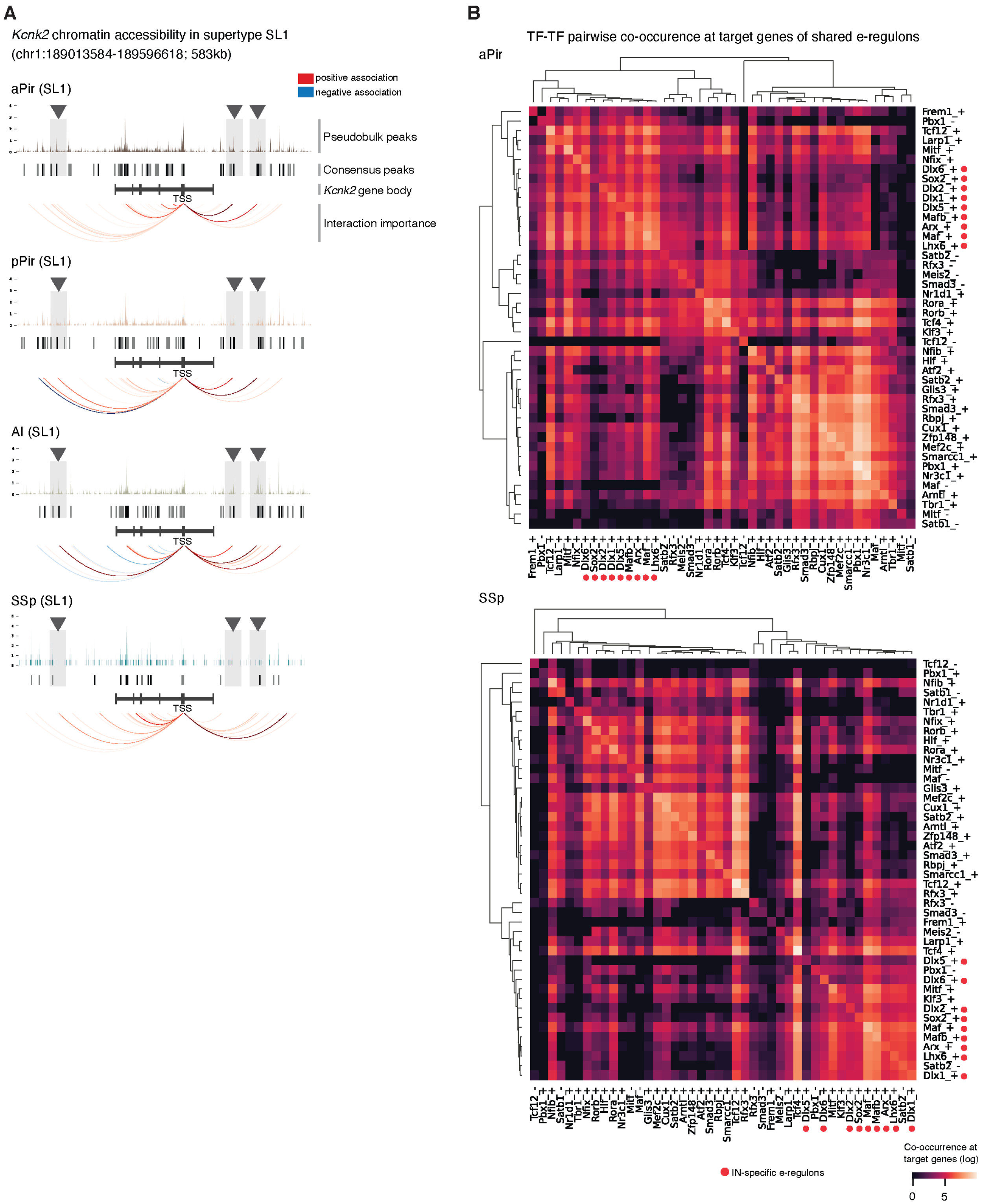
A
Kcnk2 chromatin accessibility in supertype SL1
(chr1:189013584-189596618; 583kb)
aPir (SL1)
positive association
negative association
Pseudobulk peaks
Consensus peaks
Kcnk2 gene body
Interaction importance
TSS
pPir (SL1)
AI (SL1)
SSp (SL1)
B
TF-TF pairwise co-occurence at target genes of shared e-regulons
aPir
SSp
IN-specific e-regulons
Co-occurrence at target genes (log)
0
5

**Fig. S12. Example of differential chromatin accessibility and TF combinations between shared e-regulons.**

**(A)** Genomic tracks showing chromatin accessibility in the supertype SL1 of the *Rorb*-specific target gene *Kcnk2,* which is also shared across aPir, pPir, AI, and SSp as is *Rorb*. From top to bottom rows of each area-specific track is shown: pseudobulk peaks; consensus peaks; gene body with exons as vertical bars; peaks identified as CREs indicated by arcs that go from the CRE to the TSS of the gene CREs are associated with. A positive association between a CRE and the gene has a red hue, while a negative association has a blue hue. Arcs that don't have corresponding consensus peaks represent interactions occurring in other supertypes. Triangles mark differences in CREs between areas. **(B)** Heatmaps showing TF-TF pairwise co-occurrence between e-regulons shared across cortical areas at the target genes they are predicted to regulate. Top is shown for aPir, bottom for SSp. For each e-regulon, the e-GRN-wide presence or absence of their TFs at target genes was quantified. The scale bar represents the logarithm of the co-occurrences. IN-specific e-regulons are marked with a red dot.

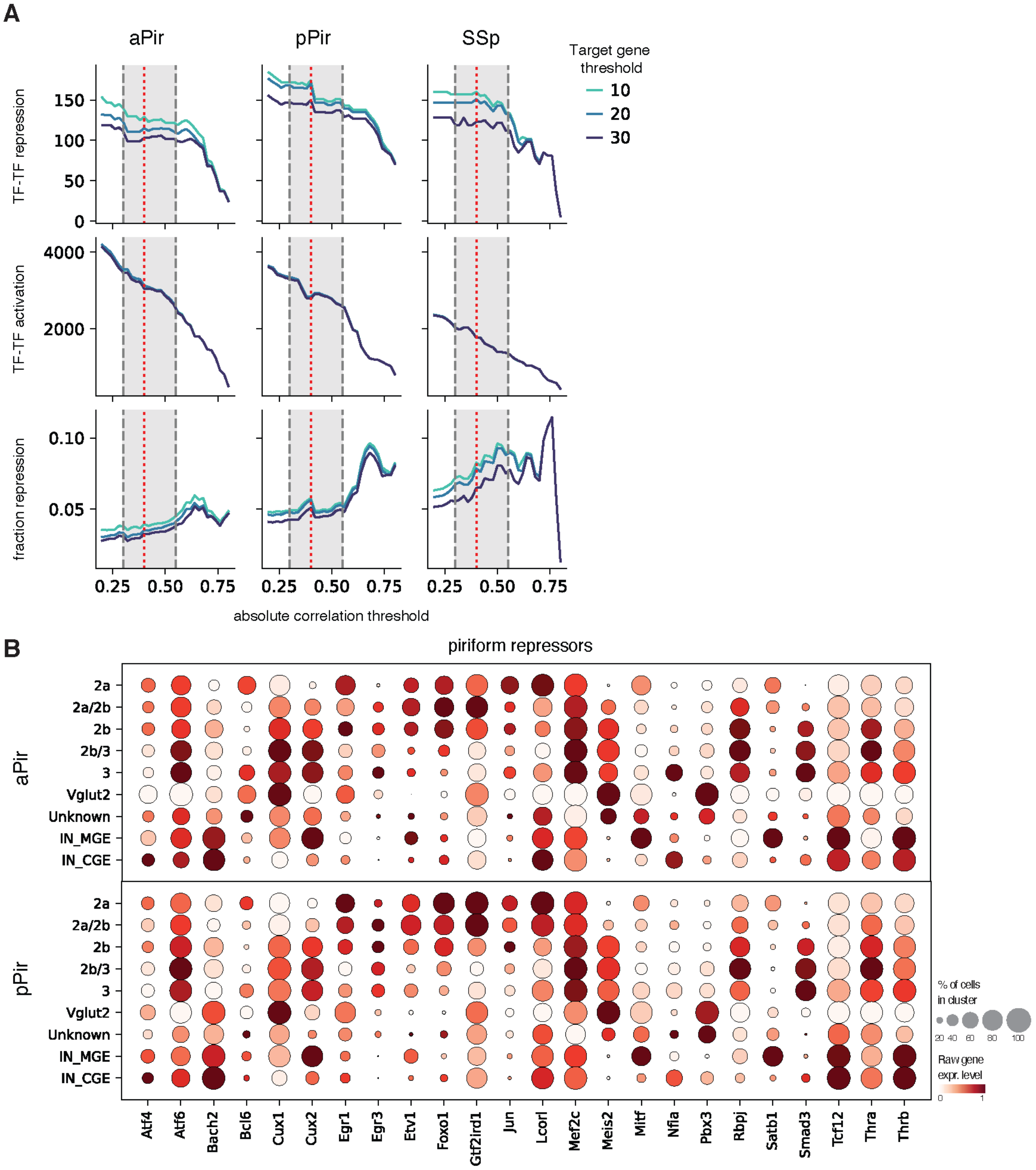


**Fig. S13. Definition of TF-TF interactions and piriform repressors.**

**(A)** Visualization of how repressive (top) and activating (bottom) TF-TF interactions are defined per cortical area, whose layer information is also available (aPir, pPir, SSp). (Anti)correlations for each e-regulon are computed between the TF, its target genes and its target CREs. An absolute correlation value is chosen between minimum and maximum absolute correlations (for activating) and anti-correlations (for repressive) thresholds (gray area). Minimum and maximum thresholds are selected based on the pattern stability of the correlations. An (anti)correlation value of 0.4 (red line) and minimum number of target genes 10 are used for all areas. Only TFs expressed in at least 20% of neurons belonging to a particular cortical layer are considered. **(B)** Dot plots showing gene expression levels of piriform repressors shown in **Fig. 2** in aPir (top) and pPir (bottom) datasets. Piriform neurons are divided by cortical layer (SL and Pyr cells), neuron type (INs and Vglut2 cells) or unknown group if layer information was not available.

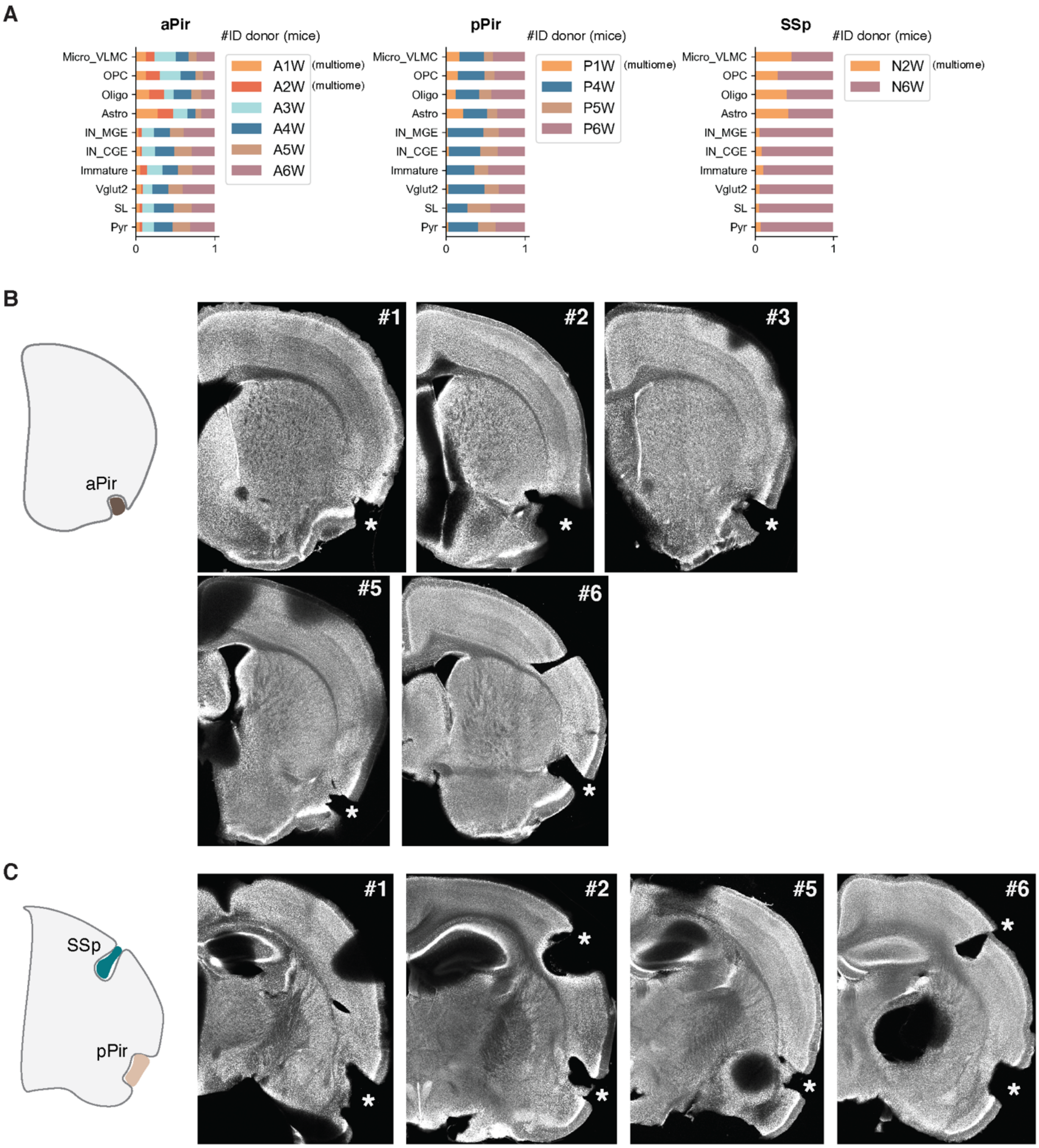
A
aPir
#ID donor (mice)
Micro_VLMC
OPC
Oligo
Astro
IN_MGE
IN_CGE
Immature
Vglut2
SL
Pyr
0
1
A1W (multiome)
A2W (multiome)
A3W
A4W
A5W
A6W
pPir
P1W (multiome)
P4W
P5W
P6W
SSp
N2W (multiome)
N6W
B
aPir
#1
#2
#3
#5
#6
C
SSp
pPir
#1
#2
#5
#6

**Fig. S14. Relative abundance of cell types and histological assessment across biological replicates of wild-derived mice.**

**(A)** Stacked bar plots representing the relative abundance of main cell types across biological replicates (donors) integrated from single-nucleus RNA (sn-RNA seq) and multiome (sn-multiome seq) sequencing experiments. Replicates IDs 1 and 2 derive from sn-multiome seq experiments, replicates IDs 3, 4, 5 and 6 derive from sn-RNA seq experiments. From left to right: aPir replicates, indicated by A; pPir replicates, indicated by P; SSp replicates, indicated by N. Numbers correspond to the ID of mice. W: wild. **(B)** Post-hoc histological assessment of aPir dissections from anterior coronal sections of adult wild-derived mice ordered by ID mouse number. Asterisks indicate the micro-dissected area. Neurotrace counterstain in gray. **(C)** Same as in **(B)** but for pPir and SSp.

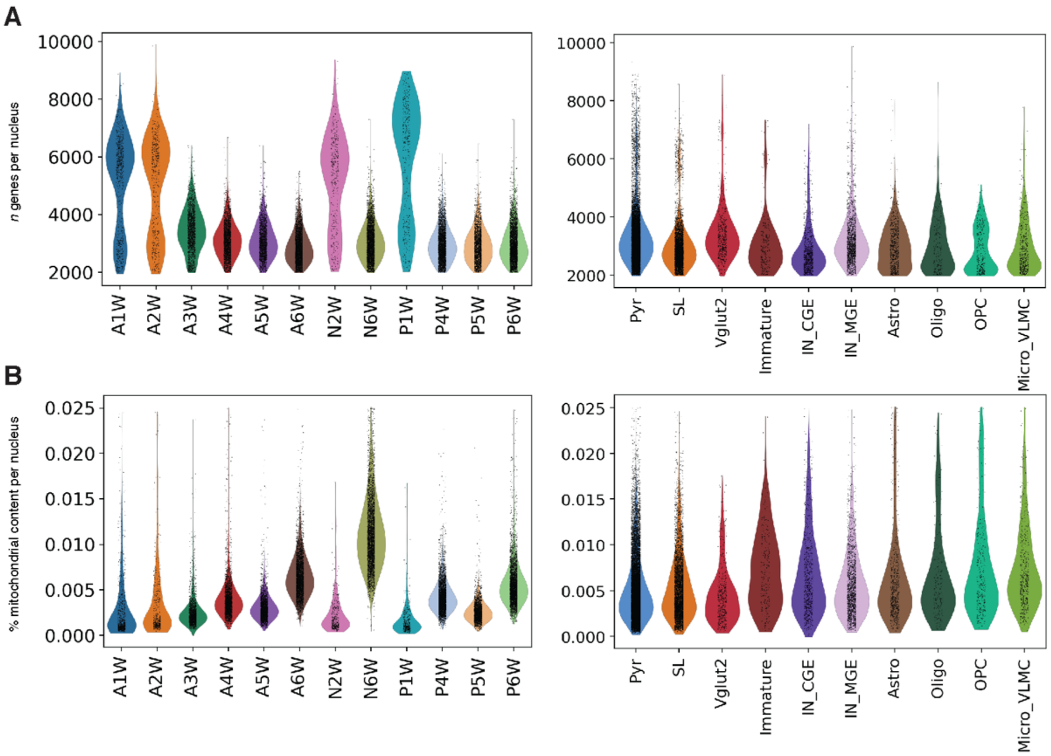


**Fig. S15. Quality control of single nucleus RNA sequencing data of wild-derived mice.**

**(A)** Left: number of genes per nucleus quantified for each biological replicate. A indicates aPir replicates, P indicates pPir, N indicates SSp. Numbers correspond to the ID of mice. Right: number of genes per nucleus identified for each main cell type. **(B)** Same as in **(A)** but for the percentage of mitochondrial content per nucleus.

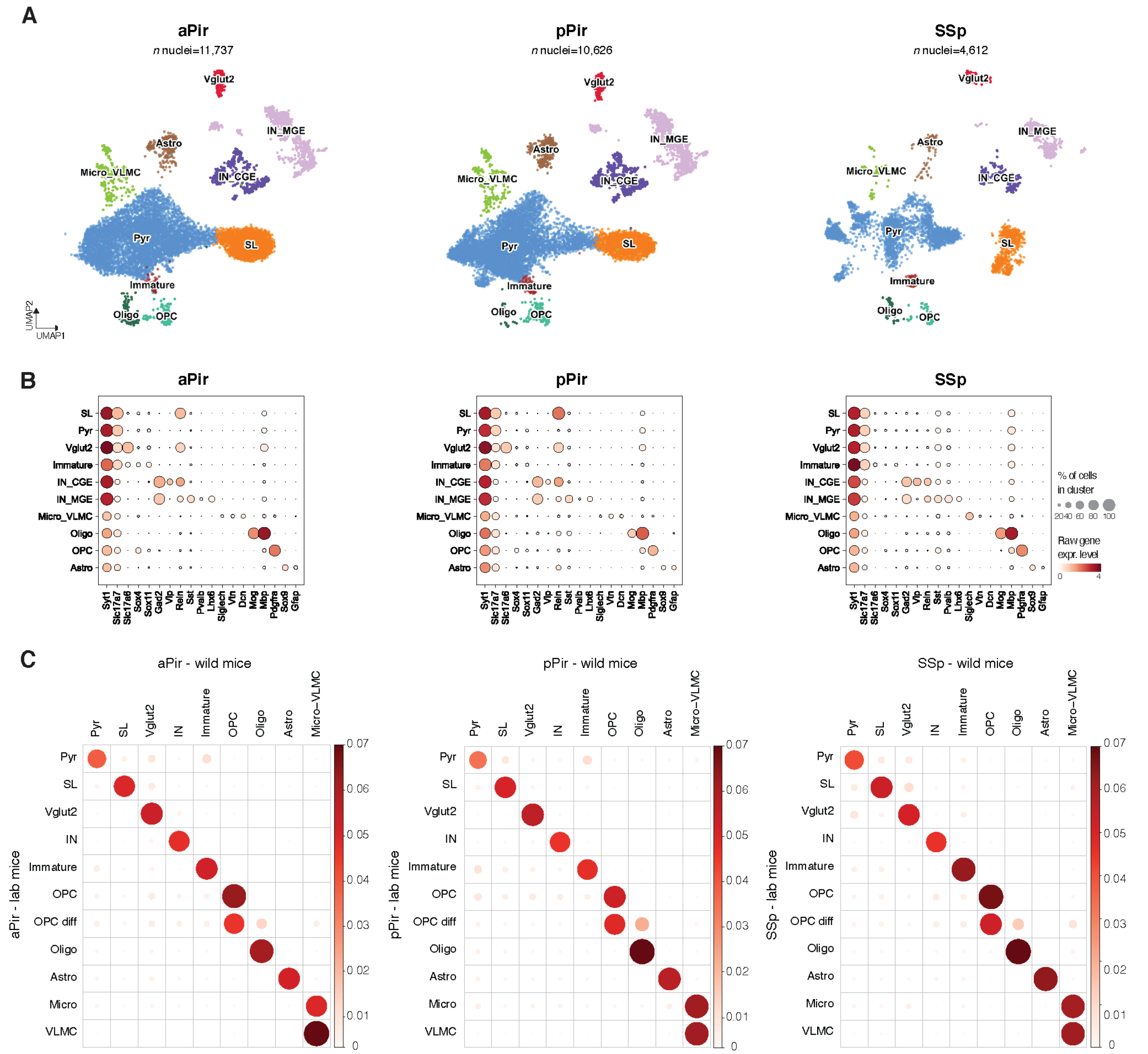


**Fig. S16. Transcriptomically-defined cell types across cortical areas of wild-derived mice.** **(A)** From left to right: UMAP representations of aPir, pPir, and SSp datasets color-coded by main cell types. **(B)** Dot plots showing gene expression levels of representative markers for each cell type across the three cortical areas, from left to right: aPir, pPir, and SSp. **(C)** Dot plots showing optimal transport (OT) alignment of main cell types between lab and wild datasets for each cortical area, from left to right: aPir, pPir, and SSp. Color and size of dots indicate the probability of alignment.

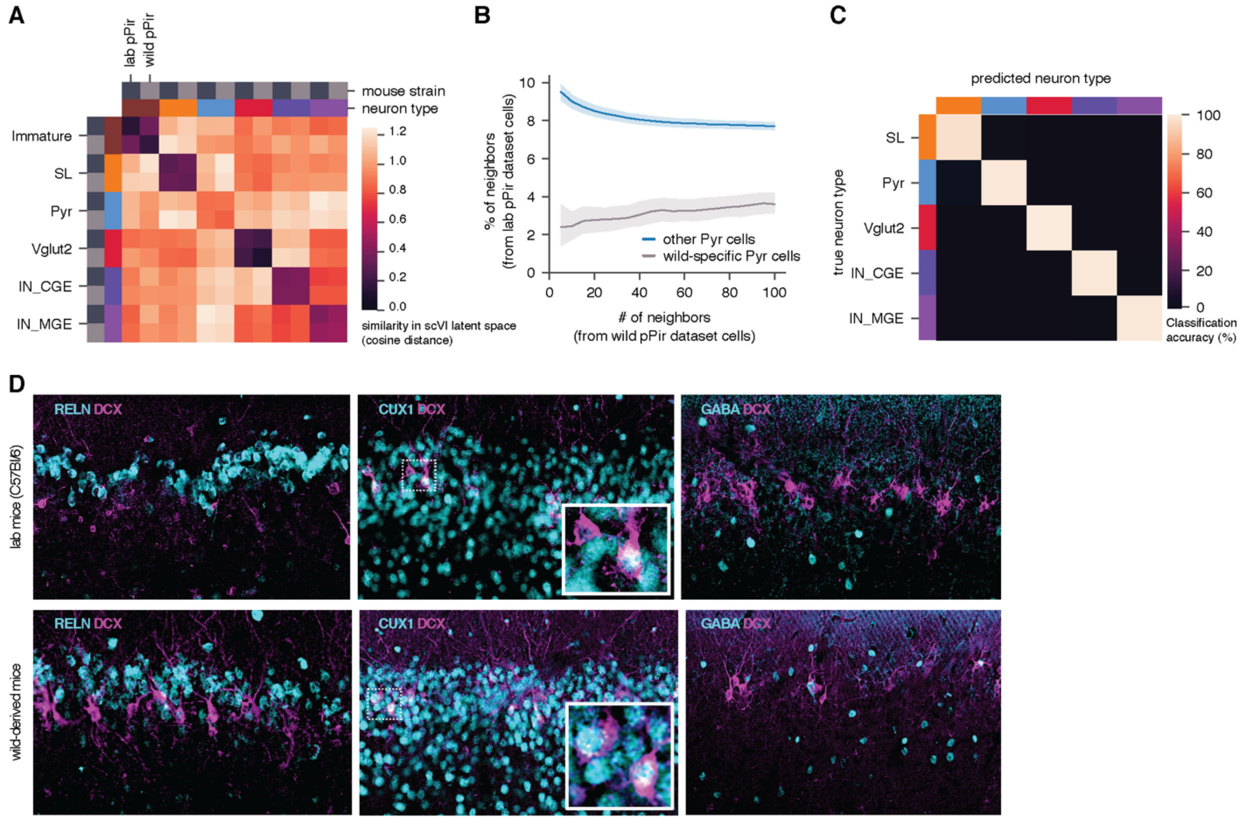


**Fig. S17. Relatedness between pyramidal and immature neurons.**

**(A)** Heatmap plot showing similarity (as cosine distance) between lab and wild neuron types upon integrating lab and wild pPir datasets using scVI as complementary approach to the optimal transport integration. Pyramidal cells were on average less similar to each other compared to the other neuron types. **(B)** Nearest neighbor graph from 5 to 100 neighbors showing the percentage of neighbors in the scVI integration shown in **(A)** as lab dataset cells, calculated for OT-identified misaligned pyramidal neurons (wild-specific pyramidal cells) and for OT-identified aligned pyramidal neurons (other pyramidal cells, molecularly similar cells between lab and wild datasets in the OT integration). **(C)** Linear Support Vector Classifier (SVC) distinguishes each neuron type from combined lab and wild pPir datasets with 98.8% accuracy. **(D)** Immunohistochemistry using lab (top) and wild-derived mice (bottom) of generic markers for piriform neuronal populations (in cyan), namely RELN for semilunar cells, CUX1 for pyramidal cells, GABA for INs, co-stained with DCX (in magenta), a canonical marker for immature neurons. Boxes with higher magnification show co-expression of CUX1 with DCX in both lab and wild-derived mice. Scale bar, 100 μm.

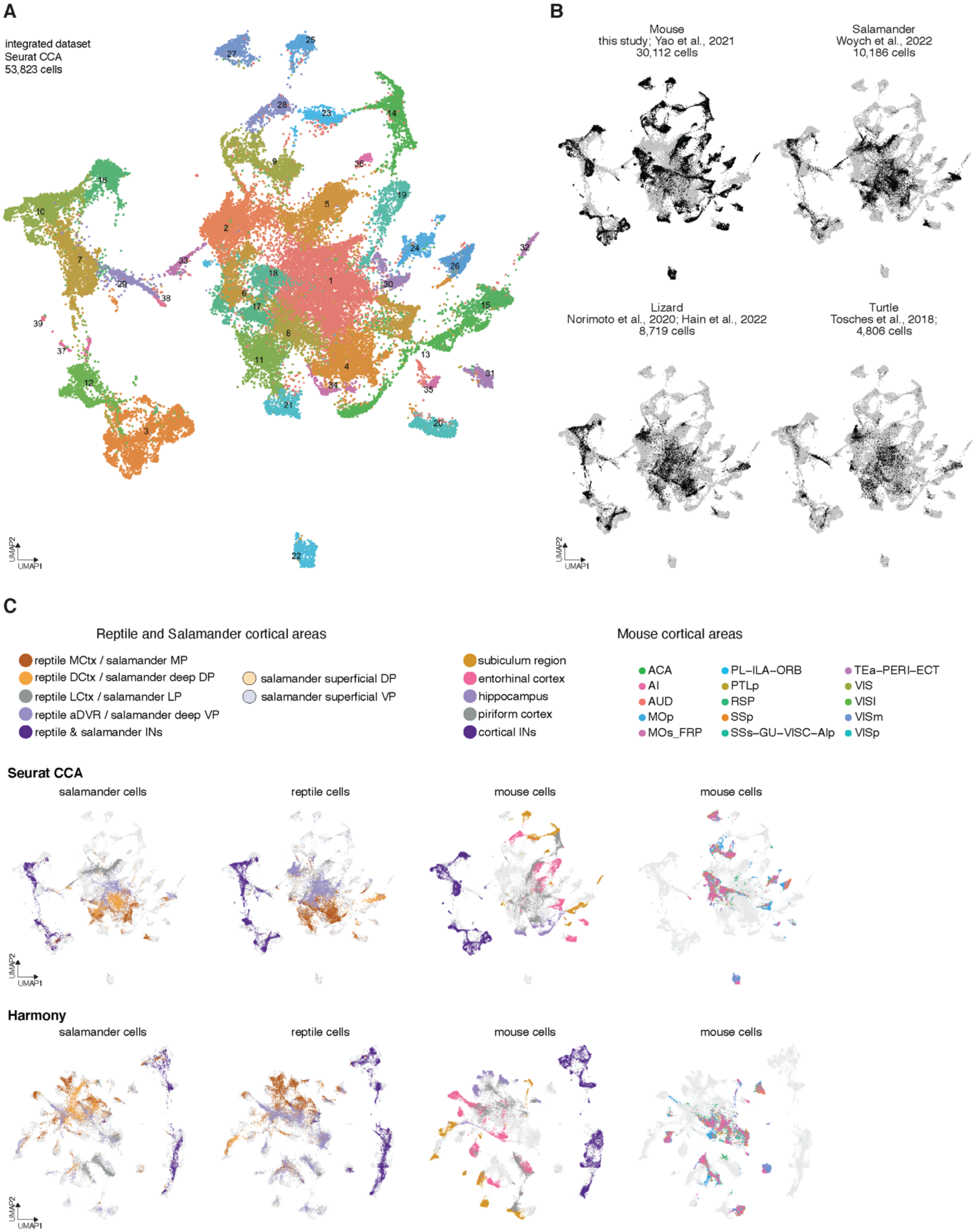
A
integrated dataset
Seurat CCA
53,823 cells
UMAP2
UMAP1
B
Mouse
this study; Yao et al., 2021
30,112 cells
Salamander
Woych et al., 2022
10,186 cells
Lizard
Norimoto et al., 2020; Hain et al., 2022
8,719 cells
Turtle
Tosches et al., 2018;
4,806 cells
C
Reptile and Salamander cortical areas
reptile MCtx / salamander MP
reptile DCtx / salamander deep DP
reptile LCtx / salamander LP
reptile aDVR / salamander deep VP
reptile & salamander INs
salamander superficial DP
salamander superficial VP
subiculum region
entorhinal cortex
hippocampus
piriform cortex
cortical INs
Mouse cortical areas
ACA
AI
AUD
MOp
MOs_FRP
PL–ILA–ORB
PTLp
RSP
SSp
SSs–GU–VISC–AIp
TEa–PERI–ECT
VIS
VISl
VISm
VISp
Seurat CCA
salamander cells
reptile cells
mouse cells
mouse cells
Harmony
salamander cells
reptile cells
mouse cells
mouse cells

**Fig. S18. Integration of neurons from cortical regions of mice, reptiles and salamander.**

**(A)** UMAP representation of the integrated sc-RNA seq datasets of mouse, turtle, lizard, and salamander pallia (medial, dorsal, lateral, and ventral cortical regions) color-coded by new seurat integrated clusters. Datasets were integrated using Canonical Correlation Analysis (CCA) in seurat. **(B)** UMAP representations of the integrated sc-RNA seq datasets shown in **(A)** highlighting in black the contribution of each species. **(C)** Comparison of two manifold integration algorithms using the datasets described in **(B)**. Integration methods used were CCA in seurat (top row) and Harmony in scanpy (bottom row), whose results were then visualized using UMAP embeddings. Similar integration results were obtained using CCA and Harmony. For each method, the first three columns show UMAP representations of species-specific cells color-coded by pallial origin of glutamatergic neurons or by INs. Turtle and lizard datasets are combined as reptile. Fourth column shows UMAP representations of mouse cells color-coded by other cortical regions. MCtx, DCtx, LCtx: medial, dorsal, lateral cortex; MP, DP, LP, VP: medial, dorsal, lateral, ventral pallium; aDVR: anterior dorsal ventricular ridge; INs: inhibitory neurons. Mouse cortical areas: ACA: anterior cingulate; AIp: posterior agranular insular; AI: agranular insular; AUD: auditory; ECT: ectorhinal; GU: gustatory; ILA: infralimbic; Mop: primary motor; MOs_FRP: secondary motor frontal pole; ORB: orbital; PAR: parasubiculum; PERI: perirhinal; PL: prelimbic; POST: postsubiculum; PRE: presubiculum; ProS: prosubiculum; PTLp: posterior parietal association; RSP: retrosplenial; SSp: primary somatosensory; SSs: secondary somatosensory; SUB: subiculum; Tea: temporal association; VIS: visual; VISC: visceral; VISl: lateral visual; VISp: primary visual.

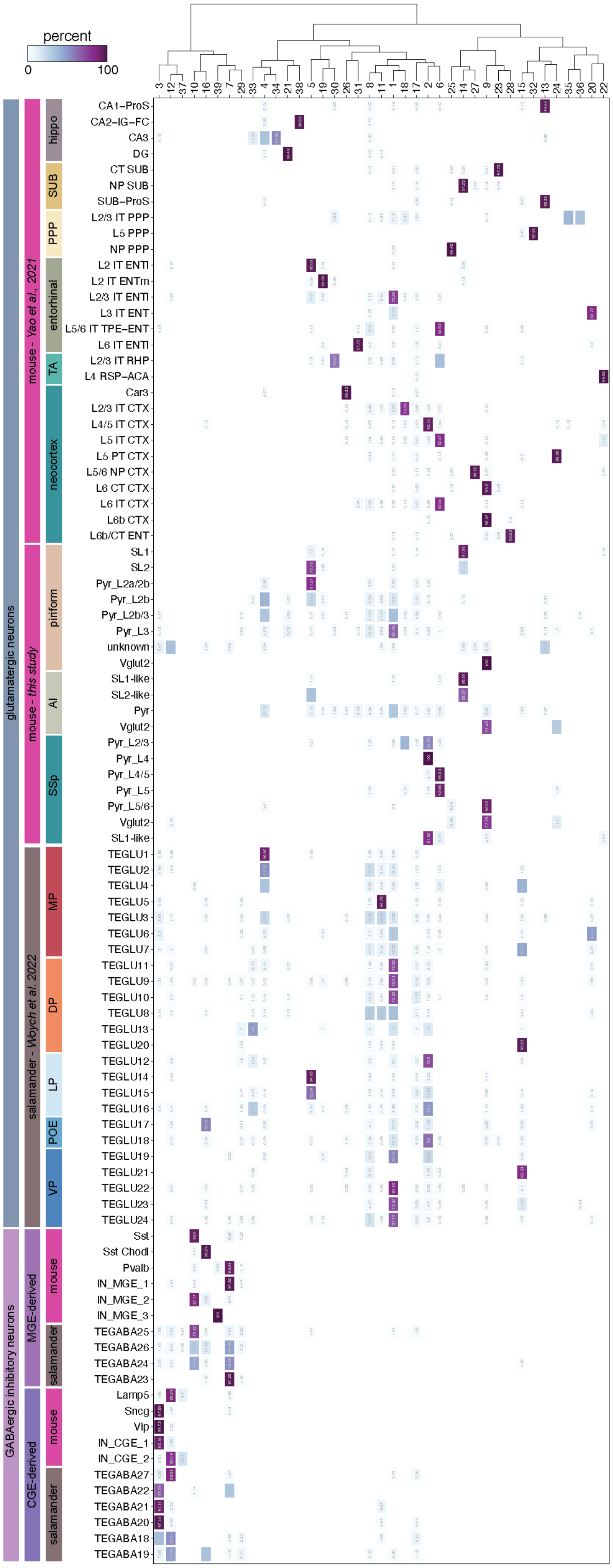
percent
0
100
glutamatergic neurons
GABAergic inhibitory neurons
mouse - Yao et al., 2021
mouse - this study
salamander - Woych et al. 2022
MGE-derived
CGE-derived
hippo
SUB
PPP
entorhinal
TA
neocortex
piriform
AI
SSp
MP
DP
LP
POE
VP
mouse
salamander
CA1–ProS
CA2–IG–FC
CA3
DG
CT SUB
NP SUB
SUB–ProS
L2/3 IT PPP
L5 PPP
NP PPP
L2 IT ENTl
L2 IT ENTm
L2/3 IT ENTl
L3 IT ENT
L5/6 IT TPE–ENT
L6 IT ENTl
L2/3 IT RHP
L4 RSP–ACA
Car3
L2/3 IT CTX
L4/5 IT CTX
L5 IT CTX
L5 PT CTX
L5/6 NP CTX
L6 CT CTX
L6 IT CTX
L6b CTX
L6b/CT ENT
SL1
SL2
Pyr_L2a/2b
Pyr_L2b
Pyr_L2b/3
Pyr_L3
unknown
Vglut2
SL1-like
SL2-like
Pyr
Vglut2
Pyr_L2/3
Pyr_L4
Pyr_L4/5
Pyr_L5
Pyr_L5/6
Vglut2
SL1-like
TEGLU1
TEGLU2
TEGLU4
TEGLU5
TEGLU3
TEGLU6
TEGLU7
TEGLU11
TEGLU9
TEGLU10
TEGLU8
TEGLU13
TEGLU20
TEGLU12
TEGLU14
TEGLU15
TEGLU16
TEGLU17
TEGLU18
TEGLU19
TEGLU21
TEGLU22
TEGLU23
TEGLU24
Sst
Sst Chodl
Pvalb
IN_MGE_1
IN_MGE_2
IN_MGE_3
TEGABA25
TEGABA26
TEGABA24
TEGABA23
Lamp5
Sncg
Vip
IN_CGE_1
IN_CGE_2
TEGABA27
TEGABA22
TEGABA21
TEGABA20
TEGABA18
TEGABA19
3 12 37 10 16 39 7 29 33 4 34 21 38 5 19 30 26 31 8 11 1 18 17 2 6 25 14 27 9 23 28 15 32 13 24 35 36 20 22

**Fig. S19. Quantification of co-clustering between mouse and salamander neurons.**

Heatmap showing quantification of co-clustering in the integrated clusters between mouse and salamander cells. The dendrogram at the top of the heatmap shows transcriptomic similarity between integrated clusters. Neuronal clusters are grouped by glutamatergic or inhibitory neurons. Glutamatergic clusters are further grouped by datasets, and each dataset by cortical areas. INs are further grouped by developmental origin (MGE- or CGE-derived INs) and by species. Rectangles indicate co-clustering of neurons (rows) in the integrated clusters (columns). Color of the rectangle represents the percentage of neurons in the integrated cluster. Hippo: hippocampus; SUB: subiculum; PPP: para-, pre-, pos-subiculum; TA: transition area; AI: agranular insular cortex; SSp: primary somatosensory cortex; MP, DP, LP, VP: medial, dorsal, lateral, ventral pallium; POE: post-olfactory eminence; MGE: medial ganglionic eminence; CGE: caudal ganglionic eminence.

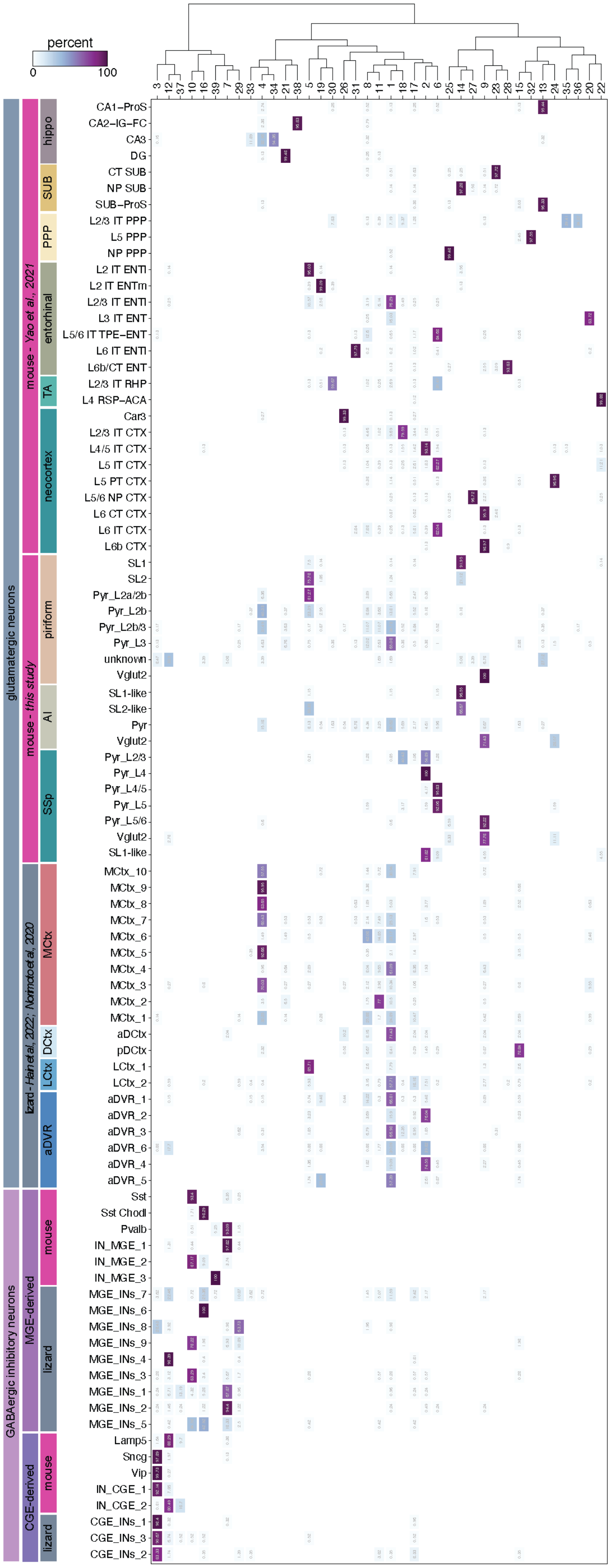
percent
0
100
3 12 37 10 16 39 7 29 33 4 34 21 38 5 19 30 26 31 8 11 1 18 17 2 6 25 14 27 9 23 28 15 32 13 24 35 36 20 22
glutamatergic neurons
GABAergic inhibitory neurons
mouse - Yao et al., 2021
mouse - this study
lizard - Hain et al., 2022; Norimoto et al., 2020
MGE-derived
CGE-derived
hippo
SUB
PPP
entorhinal
TA
neocortex
piriform
AI
SSp
MCtx
DCtx
LCtx
aDVR
mouse
lizard
CA1−ProS
CA2−IG−FC
CA3
DG
CT SUB
NP SUB
SUB−ProS
L2/3 IT PPP
L5 PPP
NP PPP
L2 IT ENTl
L2 IT ENTm
L2/3 IT ENTl
L3 IT ENT
L5/6 IT TPE−ENT
L6 IT ENTl
L6b/CT ENT
L2/3 IT RHP
L4 RSP−ACA
Car3
L2/3 IT CTX
L4/5 IT CTX
L5 IT CTX
L5 PT CTX
L5/6 NP CTX
L6 CT CTX
L6 IT CTX
L6b CTX
SL1
SL2
Pyr_L2a/2b
Pyr_L2b
Pyr_L2b/3
Pyr_L3
unknown
Vglut2
SL1-like
SL2-like
Pyr
Vglut2
Pyr_L2/3
Pyr_L4
Pyr_L4/5
Pyr_L5
Pyr_L5/6
Vglut2
SL1-like
MCtx_10
MCtx_9
MCtx_8
MCtx_7
MCtx_6
MCtx_5
MCtx_4
MCtx_3
MCtx_2
MCtx_1
aDCtx
pDCtx
LCtx_1
LCtx_2
aDVR_1
aDVR_2
aDVR_3
aDVR_6
aDVR_4
aDVR_5
Sst
Sst Chodl
Pvalb
IN_MGE_1
IN_MGE_2
IN_MGE_3
MGE_INs_7
MGE_INs_6
MGE_INs_8
MGE_INs_9
MGE_INs_4
MGE_INs_3
MGE_INs_1
MGE_INs_2
MGE_INs_5
Lamp5
Sncg
Vip
IN_CGE_1
IN_CGE_2
CGE_INs_1
CGE_INs_3
CGE_INs_2

**Fig. S20. Quantification of co-clustering between mouse and lizard neurons.**

Heatmap showing quantification of co-clustering in the integrated clusters between mouse and lizard cells. The dendrogram at the top of the heatmap shows transcriptomic similarity between integrated clusters. Neuronal clusters are grouped by glutamatergic or inhibitory neurons. Glutamatergic clusters are further grouped by datasets, and each dataset by cortical areas. INs are further grouped by developmental origin (MGE- or CGE-derived INs) and by species. Rectangles indicate co-clustering of neurons (rows) in the integrated clusters (columns). Color of the rectangle represents the percentage of neurons in the integrated cluster. Hippo: hippocampus; SUB: subiculum; PPP: para-, pre-, pos-subiculum; TA: transition area; AI: agranular insular cortex; SSp: primary somatosensory cortex; MCtx, DCtx, LCtx: medial, dorsal, lateral cortex; aDVR: anterior dorsal ventricular ridge; MGE: medial ganglionic eminence; CGE: caudal ganglionic eminence.

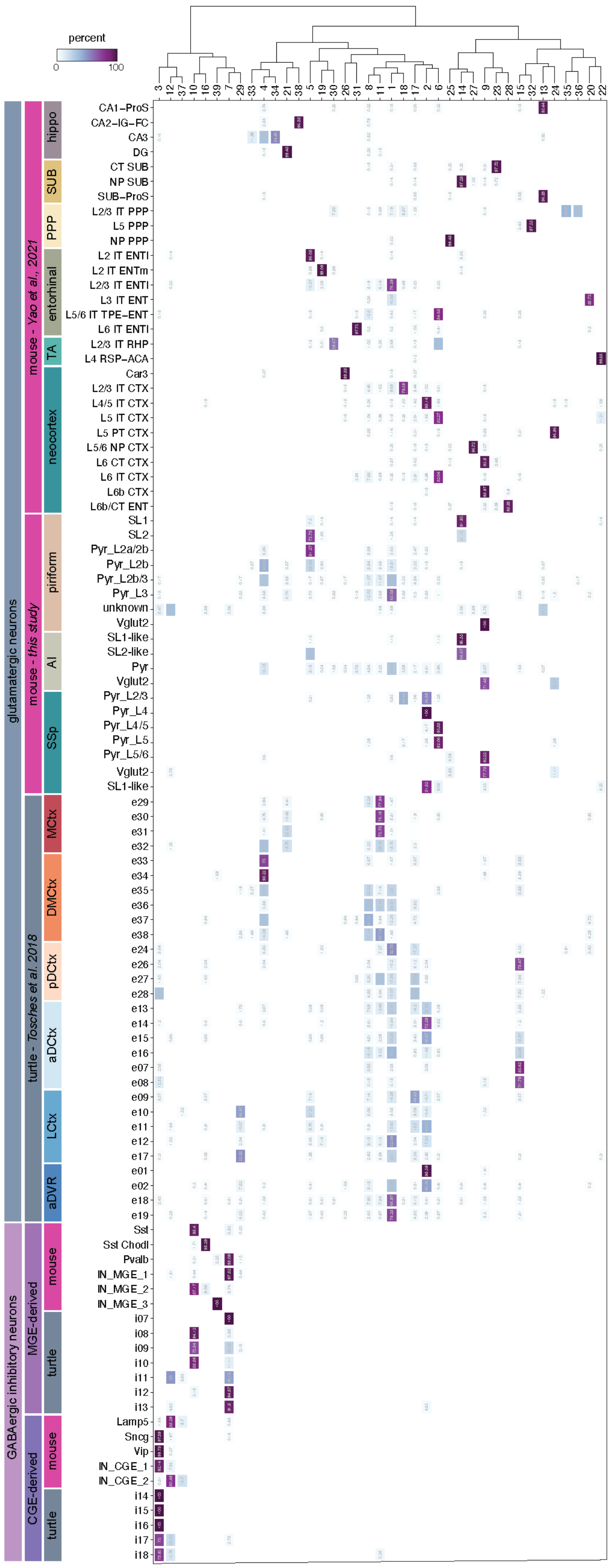
percent
0
100
3 12 37 10 16 39 7 29 33 4 34 21 38 5 19 30 26 31 8 11 1 18 17 2 6 25 14 27 9 23 28 15 32 13 24 35 36 20 22
glutamatergic neurons
GABAergic inhibitory neurons
mouse - Yao et al., 2021
mouse - this study
turtle - Tosches et al., 2018
MGE-derived
CGE-derived
hippo
SUB
PPP
entorhinal
TA
neocortex
piriform
AI
SSp
MCtx
DMCtx
pDCtx
aDCtx
LCtx
aDVR
mouse
turtle
mouse
turtle
CA1–ProS
CA2–IG–FC
CA3
DG
CT SUB
NP SUB
SUB–ProS
L2/3 IT PPP
L5 PPP
NP PPP
L2 IT ENTl
L2 IT ENTm
L2/3 IT ENTl
L3 IT ENT
L5/6 IT TPE–ENT
L6 IT ENTl
L2/3 IT RHP
L4 RSP–ACA
Car3
L2/3 IT CTX
L4/5 IT CTX
L5 IT CTX
L5 PT CTX
L5/6 NP CTX
L6 CT CTX
L6 IT CTX
L6b CTX
L6b/CT ENT
SL1
SL2
Pyr_L2a/2b
Pyr_L2b
Pyr_L2b/3
Pyr_L3
unknown
Vglut2
SL1-like
SL2-like
Pyr
Vglut2
Pyr_L2/3
Pyr_L4
Pyr_L4/5
Pyr_L5
Pyr_L5/6
Vglut2
SL1-like
e29
e30
e31
e32
e33
e34
e35
e36
e37
e38
e24
e26
e27
e28
e13
e14
e15
e16
e07
e08
e09
e10
e11
e12
e17
e01
e02
e18
e19
Sst
Sst Chodl
Pvalb
IN_MGE_1
IN_MGE_2
IN_MGE_3
i07
i08
i09
i10
i11
i12
i13
Lamp5
Sncg
Vip
IN_CGE_1
IN_CGE_2
i14
i15
i16
i17
i18

**Fig. S21. Quantification of co-clustering between mouse and turtle neurons.**

Heatmap showing quantification of co-clustering in the integrated clusters between mouse and turtle cells. The dendrogram at the top of the heatmap shows transcriptomic similarity between integrated clusters. Neuronal clusters are grouped by glutamatergic or inhibitory neurons. Glutamatergic clusters are further grouped by datasets, and each dataset by cortical areas. INs are further grouped by developmental origin (MGE- or CGE-derived INs) and by species. Rectangles indicate co-clustering of neurons (rows) in the integrated clusters (columns). Color of the rectangle represents the percentage of neurons in the integrated cluster. Hippo: hippocampus; SUB: subiculum; PPP: para-, pre-, pos-subiculum; TA: transition area; AI: agranular insular cortex; SSp: primary somatosensory cortex; MCtx, DMCtx, aDCtx and pDCtx, LCtx: medial, dorsomedial, anterior and posterior dorsal, lateral cortex; aDVR: anterior dorsal ventricular ridge; MGE: medial ganglionic eminence; CGE: caudal ganglionic eminence.

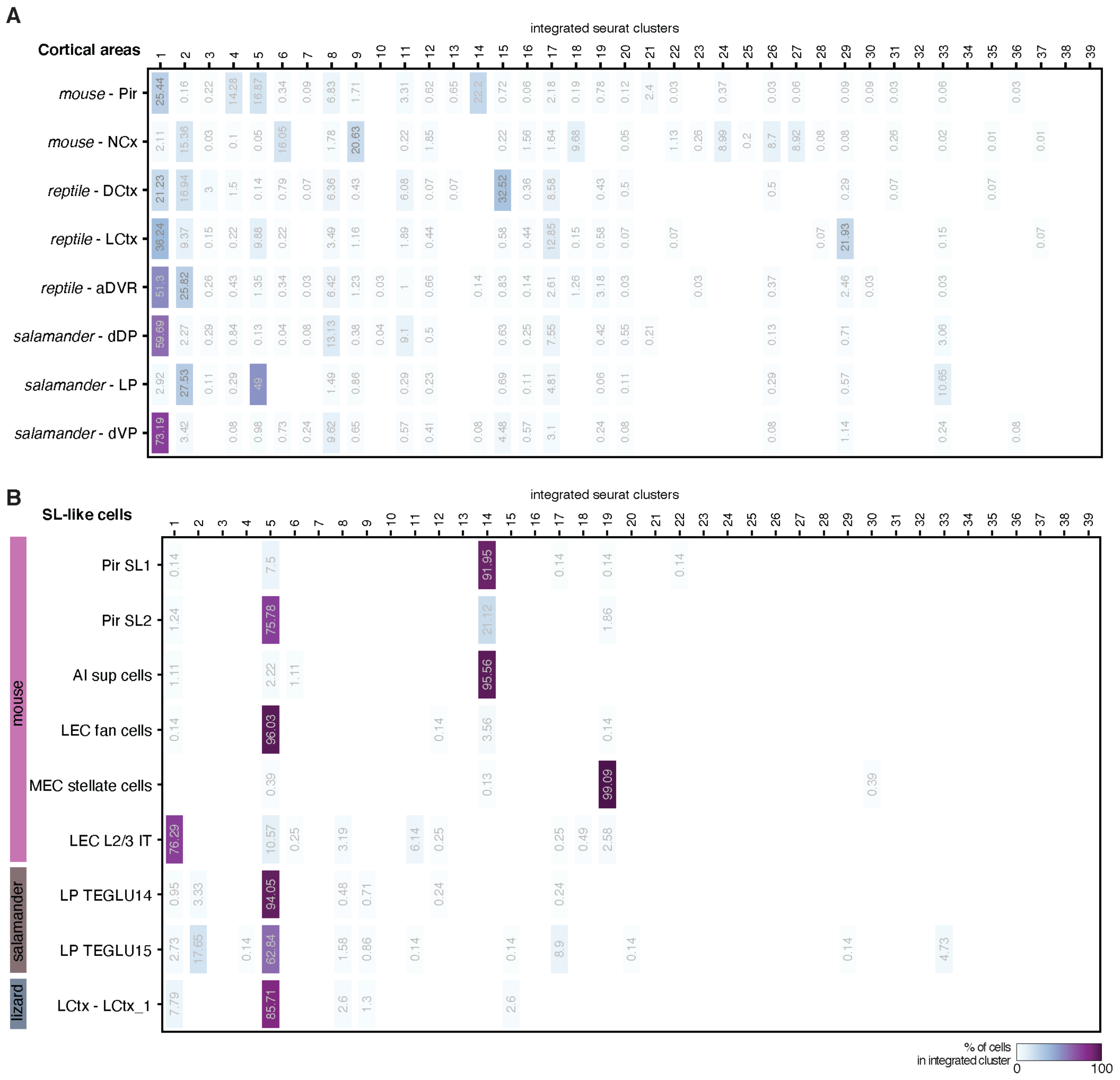


**Fig. S22. Quantification of co-clustering between glutamatergic neurons of mice, reptiles, and salamander at different resolutions.**

**(A)** Heatmap showing broad quantification of co-clustering in the integrated clusters between glutamatergic neurons from mouse and non-mammalian cortical areas, highlighting greater transcriptomic similarity of piriform glutamatergic neurons to those of non-mammals than to those of the neocortex. Pir: piriform; NCx: neocortex; DCtx, LCtx: dorsal, lateral cortex; aDVR: anterior dorsal ventricular ridge; dDP, LP, dVP: deep dorsal, lateral, deep ventral pallium. **(B)** Heatmap showing quantification of co-clustering between piriform semilunar cells and glutamatergic neurons from other cortical areas of mouse, lizard and salamander that co-clustered with piriform semilunar cells (SL-like cells). LEC, MEC: lateral and medial entorhinal cortex. In **(A)** and **(B)**, rectangles indicate co-clustering of neurons (rows) in the integrated clusters (columns). Color of the rectangle represents the percentage of neurons in the integrated cluster.

A

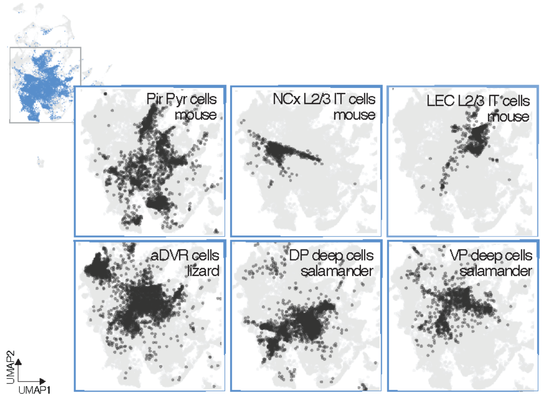


B **Pyr-like cells**

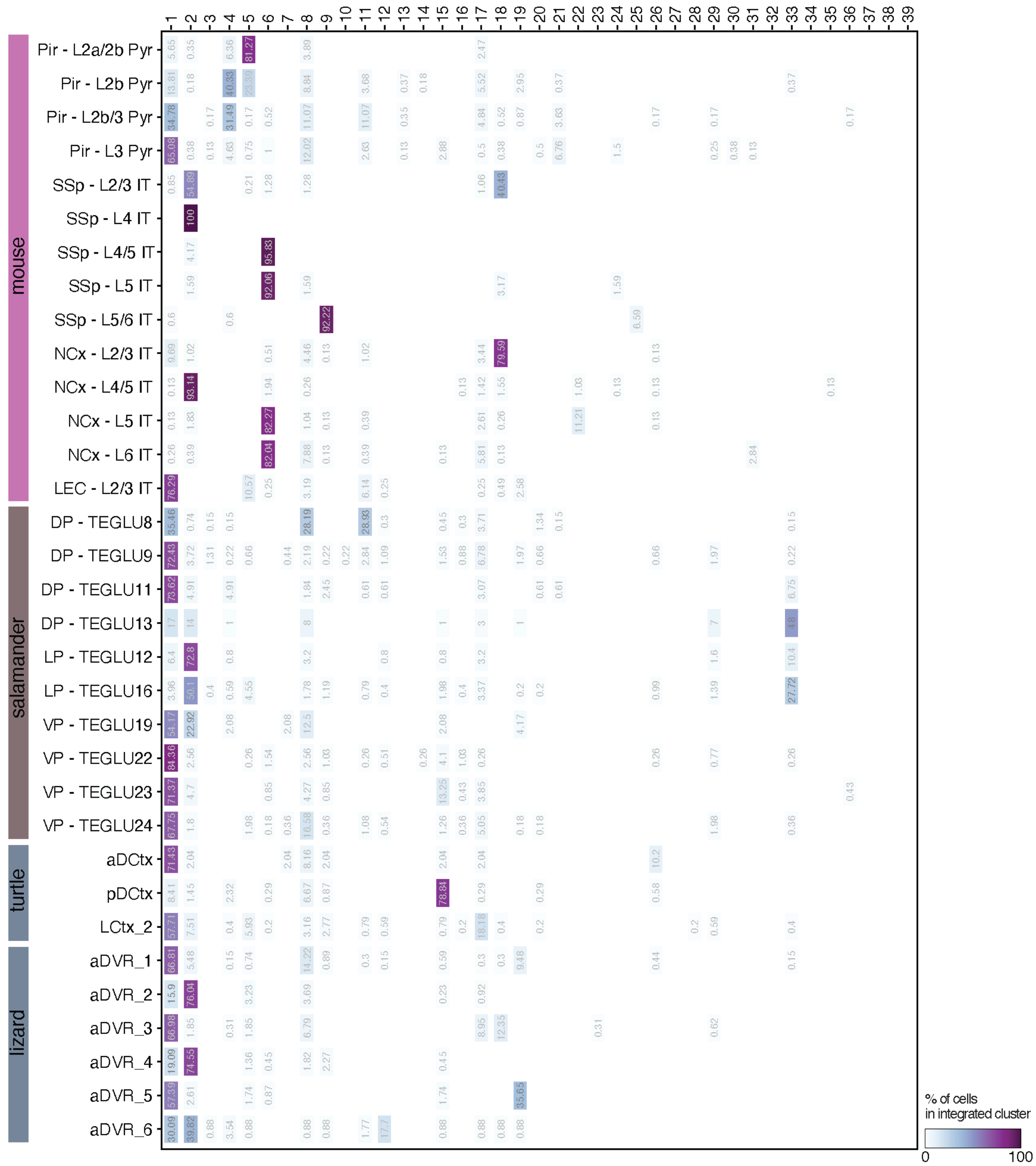

**Fig. S23. Quantification of co-clustering between piriform pyramidal (Pyr) neurons and Pyr-like neurons from other cortical areas of mouse, reptiles and salamander.**

**(A)** Top insert: UMAP representation of the integrated datasets of mice, reptiles and salamander showing in blue all integrated clusters that comprise piriform pyramidal cells. Rest: UMAP representations of neurons from mouse and non-mammalian cortical areas that co-clustered with piriform pyramidal cells in the integrated clusters. **(B)** Heatmap showing quantification of co-clustering between piriform pyramidal cells and glutamatergic neurons from other cortical areas of mouse, lizard, turtle, and salamander that co-clustered with piriform pyramidal cells (Pyr-like cells). Rectangles indicate co-clustering of neurons (rows) in the integrated clusters (columns). Color of the rectangle represents the percentage of neurons in the integrated cluster. Pir: piriform; NCx L2/3 IT: neocortex layer 2/3 intratelencephalic; LEC 2/3 IT: lateral entorhinal cortex layer 2/3 intratelencephalic; aDVR: anterior dorsal ventricular ridge; DP, LP, VP: dorsal, ventral pallium; DCtx, LCtx: dorsal, lateral cortex; SSp: primary somatosensory cortex.

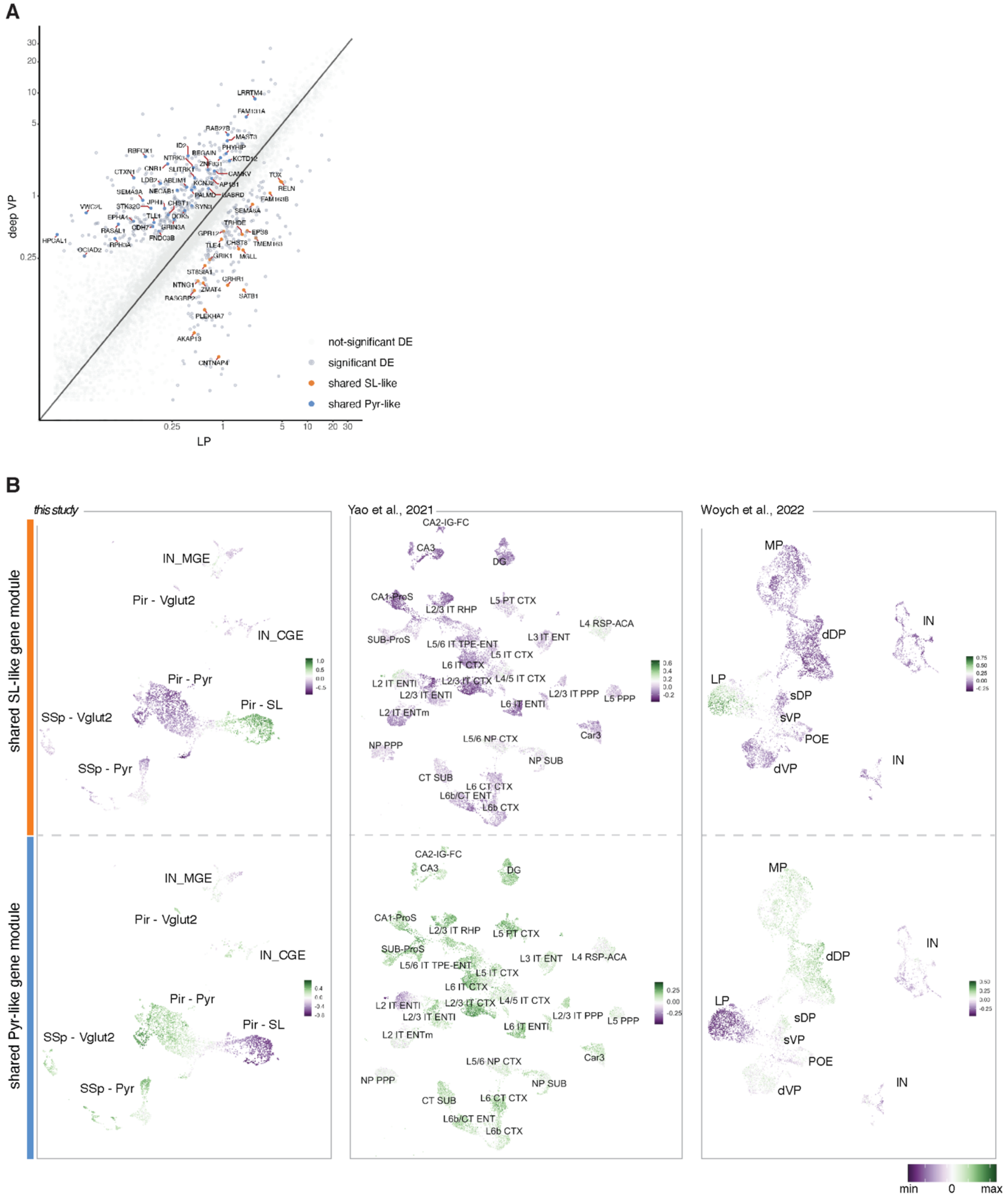
A
deep VP
LP
not-significant DE
significant DE
shared SL-like
shared Pyr-like
LRRTM4
FAM131A
RAB27B
MAST3
PHYHIP
KCTD12
CAMKV
TOX
RELN
SEMA6A
TRHDE
EPS8
TMEM163
CHST8
GRIK1
MGLL
TLE4
GPR12
ST8SIA1
CRHR1
NTNG1
ZMAT4
SATB1
RASGRP2
PLEKHA7
AKAP13
CNTNAP4
RBFOX1
CTXN1
CNR1
NTRK1
SLITRK1
LDB2
ABLIM1
SEMA3A
NECAB1
STK32C
JPH1
CHST1
SYN1
VWC2L
EPHA4
TLL1
CDH7
GRIN3A
RASAL1
FNDC3B
HPCAL1
RPH3A
OCIAD2
B
this study
Yao et al., 2021
Woych et al., 2022
shared SL-like gene module
shared Pyr-like gene module
IN_MGE
Pir - Vglut2
IN_CGE
Pir - Pyr
Pir - SL
SSp - Vglut2
SSp - Pyr
CA2-IG-FC
CA3
DG
CA1-ProS
L2/3 IT RHP
L5 PT CTX
L4 RSP-ACA
SUB-ProS
L5/6 IT TPE-ENT
L3 IT ENT
L5 IT CTX
L6 IT CTX
L2 IT ENTl
L2/3 IT CTX
L4/5 IT CTX
L2/3 IT ENTl
L2/3 IT PPP
L5 PPP
L6 IT ENTl
L2 IT ENTm
Car3
L5/6 NP CTX
NP PPP
NP SUB
CT SUB
L6 CT CTX
L6b/CT ENT
L6b CTX
MP
IN
dDP
LP
sDP
sVP
POE
dVP
min
0
max

**Fig. S24. SL-like and Pyr-like gene signatures across areas and species.**

**(A)** Scatter plot of differentially expressed genes (DEGs) between cells from LP and VP. Each dot represents the mean gene expression value. Orange and blue dots depict shared genes enriched both in semilunar and LP neurons, or in pyramidal and VP neurons, respectively. Other statistically significant DEGs are shown in dark gray, while non-statistically significant DEGs in light gray. **(B)** UMAP representations of SL-like (top) and Pyr-like (bottom) gene module scores across datasets, from left to right: *this study* (mouse - Pir, AI, SSp); Yao et al., 2021 (mouse - dorsal-medial-lateral cortex); Woych et al., 2022 (salamander - MP, LP, superficial(s) and deep(d) DP and VP, POE). Scale bar indicates the score of the gene signatures. A positive score indicates that the set of genes in the module are expressed in a particular cluster more highly compared to the average expression across all clusters.
